# Discovery of small molecule antagonists of human Retinoblastoma Binding Protein 4 (RBBP4)

**DOI:** 10.1101/2022.01.05.475077

**Authors:** Sumera Perveen, Carlos A. Zepeda-Velázquez, David McLeod, Richard Marcellus, Mohammed Mohammed, Megha Abbey, Deeba Ensan, Dimitrios Panagopoulos, Viacheslav Trush, Elisa Gibson, Aiping Dong, Levon Halabelian, Gennady Poda, Julie Owen, Ahmed Aman, Taira Kiyota, Ahmed Mamai, Cheryl H. Arrowsmith, Peter J. Brown, Matthieu Schapira, Rima Al-awar, Masoud Vedadi

**Author notes:** To whom correspondence should be addressed: Masoud Vedadi; Rima Al-awar. Contributed equally (Sumera Perveen, Carlos A. Zepeda-Velázquez).

## Abstract

RBBP4 is a nuclear WD40 motif-containing protein widely implicated in various cancers and a putative drug target. It interacts with multiple proteins within diverse complexes such as nucleosome remodeling and deacetylase (NuRD) complex and polycomb repressive complex 2 (PRC2), as well as histone H3 and H4 through two distinct binding sites. B-cell lymphoma/leukemia 11A (BCL11A), friend of GATA-1 (FOG-1), plant homeodomain finger protein 6 (PHF6) and histone H3 bind to the top of the donut-shaped seven-bladed β-propeller fold of RBBP4, while suppressor of zeste 12 (SUZ12), metastasis associated protein 1 (MTA1) and histone H4 bind to a pocket on the side of the WD40 repeats of this protein. Here, we report the discovery of the first small molecule antagonists of the RBBP4 top pocket, competing with interacting peptides from proteins such as BCL11A and histone H3. We also determined the first crystal structure of RBBP4 in complex with a small molecule (OICR17251), paving the path for structure-guided design and optimization towards more potent antagonists.

## Introduction

Retinoblastoma binding protein 4 (RBBP4) and 7 (RBBP7) also known as retinoblastoma associated protein 48 (RbAp48) and 46 (RbAp46), respectively, are highly homologous nuclear WD40 motif-containing proteins.^1–2^ RBBP4 and 7 (RBBP4/7) have been widely implicated in various cancers and are valuable drug targets. RBBP4 is overexpressed in several cancers including hepatocellular carcinoma^3^, and acute myeloid leukemia^4–5^. Reduction of RBBP4 expression using siRNA results in the suppression of tumorigenicity. RBBP4 expression is upregulated in human thyroid carcinoma, and its downregulation suppresses tumorigenicity.^6–7^ Decreases in RBBP4 expression in glioblastoma cells also enhance sensitivity towards the chemotherapeutic temozolomide. ^8^ Elevated levels of RBBP4/7 in non-small cell lung cancer and inhibition of lung cancer cells migration upon RBBP7 knockdown have been reported.^9–10^ RBBP7 is also overexpressed in other cancers such as renal cell carcinoma, medulloblastoma and breast cancer.^11–13^ Upregulation of RBBP7 in bladder cancer results in an invasive cell phenotype, that can be ameliorated through a reduction in its expression.^14^

RBBP4 and RBBP7 are highly homologous proteins (89% sequence identity) with similar structures. Both are donut-shaped proteins, each with a seven-bladed β-propeller fold typical of WD40-repeat proteins. Distinctive features of these proteins include a prominent *N*-terminal α-helix, a negatively charged PP-loop, and a short C-terminal α-helix.^15^ Two binding sites have been identified on these two proteins, a c-site on the top of the WD40 repeats (**Suppl. Fig. 1A**)^16–18^ and another located on the side of the WD40 domain between the *N*-terminal α-helix and PP-loop (Ser-347 to Glu-364 in RBBP7) (**Suppl. Fig. 1B**)^18–20^. RBBP4/7 are components of several multi-protein complexes involved in chromatin remodeling, histone post-translational modifications, and regulation of gene expression suggesting diverse functions for RBBP4/7. Such complexes include nucleosome remodeling and deacetylase (NuRD) complex^21^, switch independent 3A (Sin3A)^22^, and polycomb repressive complex 2 (PRC2)^23–24^. Some interaction partners of RBBP4/7 include metastasis associated protein 1 (MTA1)^21, 25^, also a component of NuRD complex, and suppressor of zeste 12 (SUZ12)^18, 24^ within the PRC2 complex. Structural studies have shown that these interactions, along with binding to histone H4^20^, are through the side pocket on the WD40 domain of RBBP4/7 and Nurf55 (*Drosophila* homolog of RBBP4/7) (**Suppl. Fig. 1B**). In addition, the C-terminal α-helix may be involved in binding of the peptides to this site. This pocket is unique to RBBP4/7 and has both negatively charged and hydrophobic areas (**Suppl. Fig. 2**). As a result, hydrophobic interactions play important roles in Su(z)12-Nurf55 binding. It has been shown that binding of H4, MTA1 and Su(z)12 to RBBP4/7 and Nurf55 are mutually exclusive.^18–19^ Interestingly, RBBP4/7 can bind to MTA1, histone H3 or FOG-1 independently. However, within the NuRD complex, the presence of MTA1 can destabilize the interaction of RBBP4/7 with histones H3-H4^19^.

Crystal structures of RBBP4 in complex with PHF6^16^, FOG-1^17^, and Nurf55 with histone H3^18^ peptides reveal these ligands bind to the same position in the c-site on top of the WD40 repeats, forming multiple hydrogen bonds. This site is negatively charged, creating a surface perfectly suited for interaction with the positively charged residues of the peptides (**Suppl. Fig. 2**). In all these interactions, an arginine side chain of the peptide (Arg-2 in H3, Arg-163 in PHF6 and Arg-4 in FOG-1) is directly inserted into the negatively charged central channel of RBBP4/7 and forms a hydrogen bond with Glu-231 (Glu-235 in Nurf55) (PDB ID: 2YBA, 4R7A and 2XU7). Furthermore, all peptides adopt the same binding pose (**Suppl. Fig. 1A**). This observation suggests that PHF6, FOG-1 and H3 interactions with RBBP4/7 and Nurf55 are highly conserved.

BCL11A, a transcription factor, has been shown to interact with the RBBP4 top pocket ^26^. The crystal structure revealed that BCL11A (**Suppl. Fig. 1A**) binds to the c-site similar to H3 ^18^, PHF6^16^, and FOG-1^17^. In BCL11A, residues 4-13 are involved in binding to the top of the β-propeller fold of RBBP4 in a similar manner as residues 2-11 of the histone H3 with arginine 4 making a hydrogen bond with Glu-231 of RBBP4 (PDB ID: 5VTB).

Here we report the development of Fluorescence Polarization (FP)-based peptide displacement assays that we used to screen RBBP4 against a library of 20,000 diverse small molecules to identify antagonists of BCL11A and H3 interactions with the top binding site.

Through structure-activity relationship (SAR) studies, we identified OICR17251 as an antagonist of the RBBP4-H3 interaction with a K_disp_ of 23 ± 1 μM, and further characterized its interaction through surface plasmon resonance (SPR) and determination of co-crystal structure. This discovery paves the way towards the development of more potent antagonists of RBBP4 interactions with various proteins, providing a promising starting point for the development of cancer therapeutics.

## Results and Discussion

The crystal structures of RBBP4 in complex with BCL11A (aa 2-16)^26^, H3 (aa 1-21)^18^ and MTA1 (aa 656-686) ^19^ peptides were previously reported. We used these peptides to develop fluorescence polarization (FP)-based peptide displacement assays for screening RBBP4 in a 384-well format to identify small molecules that bind to the RBBP4 top (BCL11A and H3 peptides) and side (MTA1 peptide) pockets (**Suppl. Table 1 and 2**). In these assays, binding of the fluorescein-labeled peptides to RBBP4 was assessed by monitoring the increase in the fluorescence polarization signal^27^. The unlabeled peptides with the same amino acid sequences were used to displace the labeled peptides as controls (**Fig. 1 and Suppl. Fig. 3**). The K_D_ values were determined for all three FITC-labeled peptides (**Fig. 1A, C, E**), and confirmation of their displacement by their unlabeled versions indicated that these assays can be used for small molecule screening (**Fig. 1B, D, F**). The FITC-BCL11A (2-16) peptide was displaced from RBBP4 by unlabeled BCL11A (2-16) (**Fig. 1B**) and H3 (1-21) (**Suppl. Fig. 3**) with K_disp_ values of 1.2 ± 0.009 µM and 2.5 ± 0.03 µM, respectively. Similarly, unlabeled BCL11A (2-16) displaced FITC-H3 (1-21) peptide with K_disp_ of 0.9 ± 0.094 µM (**Fig. 1D**), indicating that both BCL11A (2-16) and H3 (1-21) compete for binding to the top pocket of RBBP4 as expected. Additionally, unlabeled MTA1 (656-686) peptide displaced FITC-MTA (656-686) from the side pocket of RBBP4 with K_disp_ of 0.24 ± 0.02 µM (**Fig. 1F**). To confirm the reproducibility and suitability of the assay for RBBP4 screening, the displacement assays were performed in a 384-well format and Z′-factors of 0.69, 0.59 and 0.55 were determined for displacement with BCL11A (2-16), H3 (1-21) and MTA (656-686) peptides, respectively (**Suppl. Fig. 4**).

**Figure 1.**
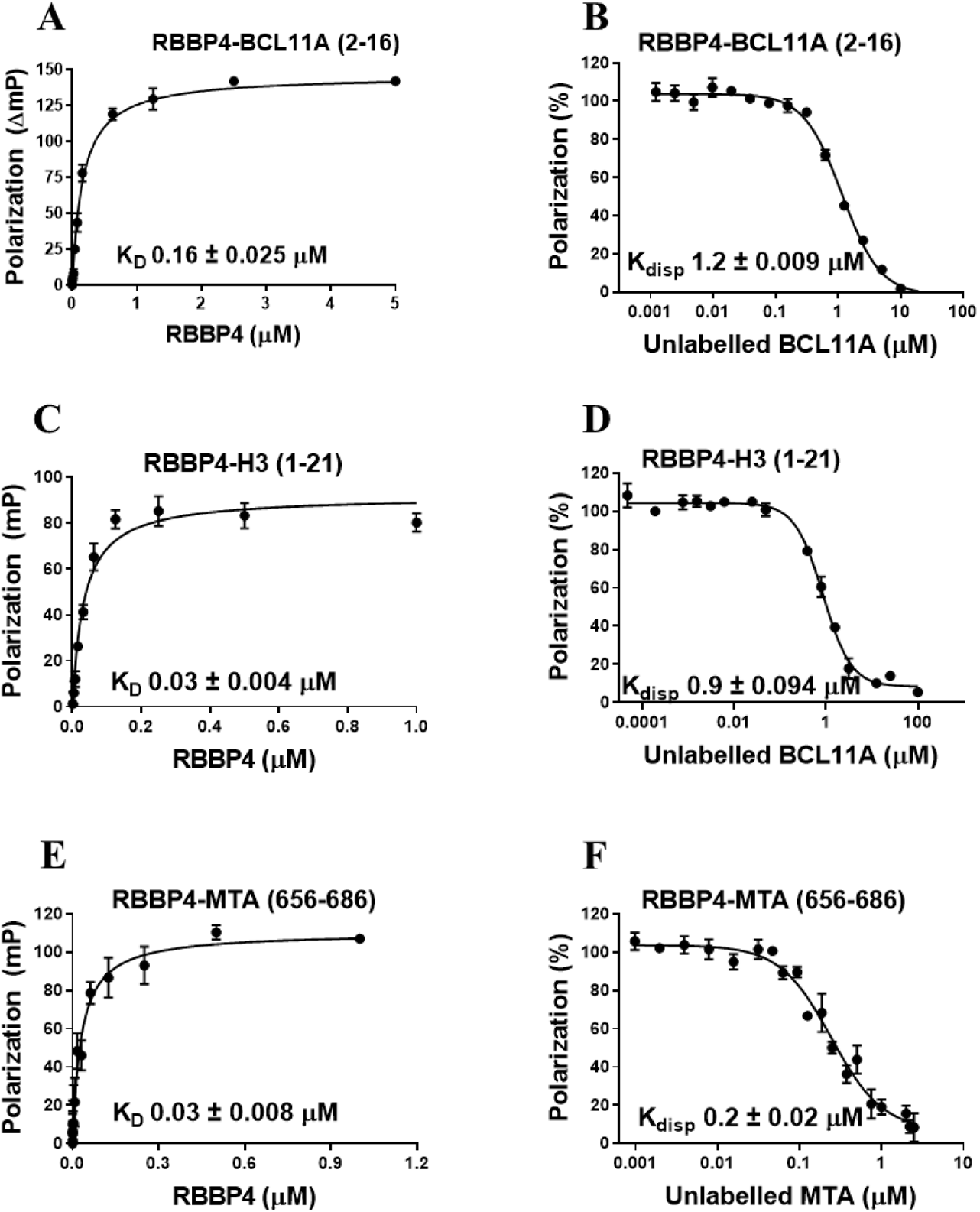
Characterization of RBBP4 interaction with peptides. Binding of the FITC-labeled peptides to RBBP4 and their displacement by unlabeled version of each peptide for (A, B) BCL11A(2-16), (C, D) H3(1-21), and (E, F) MTA(656-686) peptides are presented. The K_D_ (A, C, and E) and K_disp_ (B, D, and F) values are presented on each related figure. All experiments were performed in triplicate, and data are shown as the mean ± standard deviation.

The strong correlation between RBBP4 and cancer ^3, 6, 9, 11, 13^ motivated us to screen our internal collection of 20,000 diverse small molecules to identify binding antagonists to either the canonical top or side pockets. Screening campaigns were performed in parallel using the suitable peptides for each pocket. Screening compounds of interest were triaged, and hits were confirmed by K_disp_ determination. From these efforts, four singletons and one structural cluster were identified as binders of the canonical site (top pocket). The structural cluster consisted of compounds containing a deazapurine core with an aliphatic amine side chain. OICR14833 was then identified to bind to the top pocket of RBBP4 by displacing BCL11A (2-16) and H3 (1-21) with K_disp_ of 114 ± 13 μM (**Fig. 2A**) and 74 ± 7 μM (**Fig. 2B**), respectively. Additionally, OICR14833 was shown to displace the MTA1 656-686 peptide from the side binding pocket of RBBP4 with a K_disp_ of 26 ± 0.3 μM (**Fig. 2C**). Weak binding of OICR14833 to RBBP4 was also confirmed by SPR (**Fig. 3A, B and Fig. 4**). However, the K_D_ value was not calculated due to its apparent weak binding and linear fitting (**Fig. 3A**).

**Figure 2.**
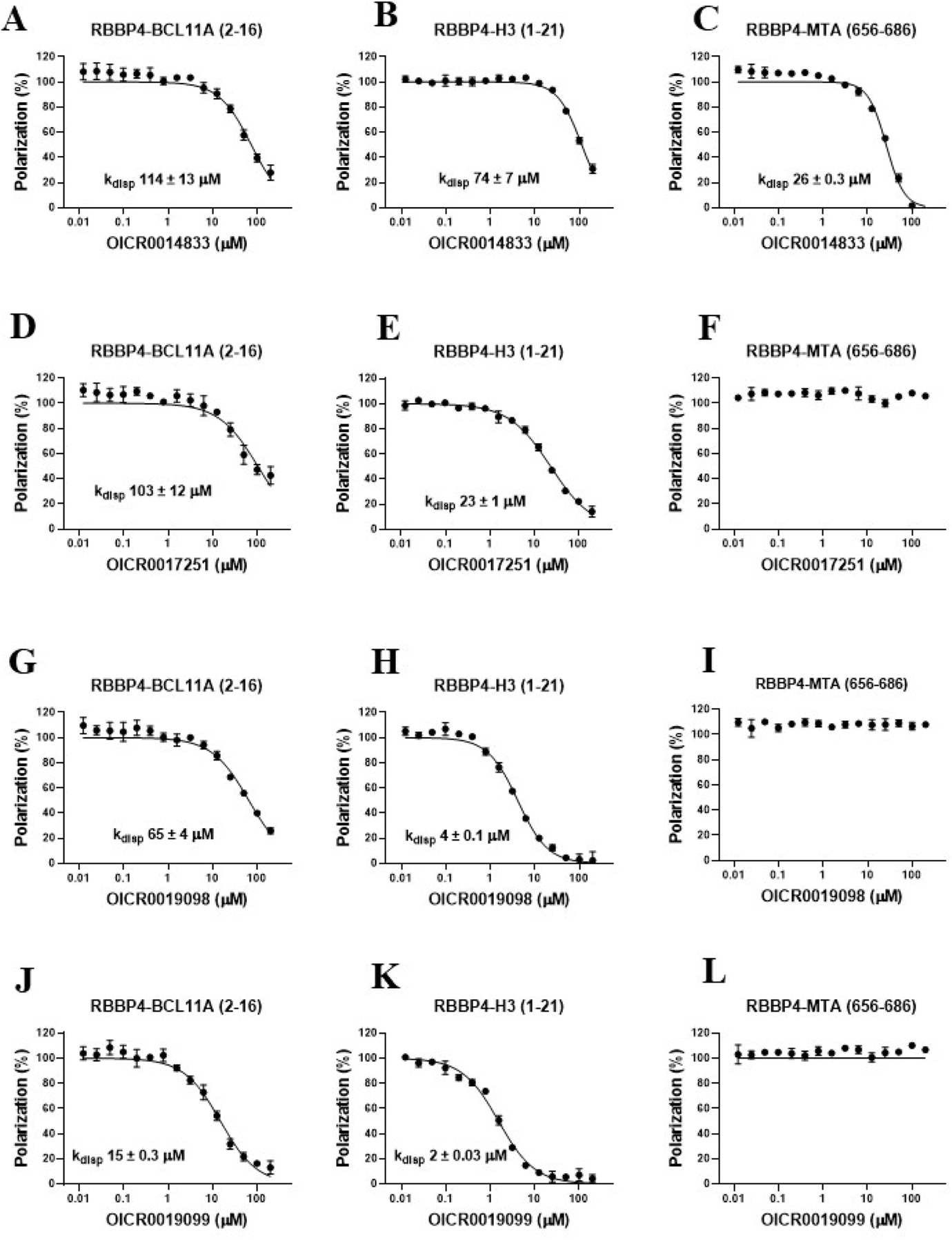
Assessing the binding of OICR14833, OICR17251, OICR19098, and OICR19099 to RBBP4 by peptide displacement assay. All three compounds were tested for displacement of (A, D, G, J) FITC-BCL11A (2-16), and (B, E, H, and K) FITC-H3 (1-21) peptides from the top pocket, and (C, F, I, and L) FITC-MTA (656-686) peptide from the side pocket. The K_disp_ values are presented on each figure. All experiments were performed in triplicate, and data are shown as the mean ± standard deviation.

**Figure 3:**
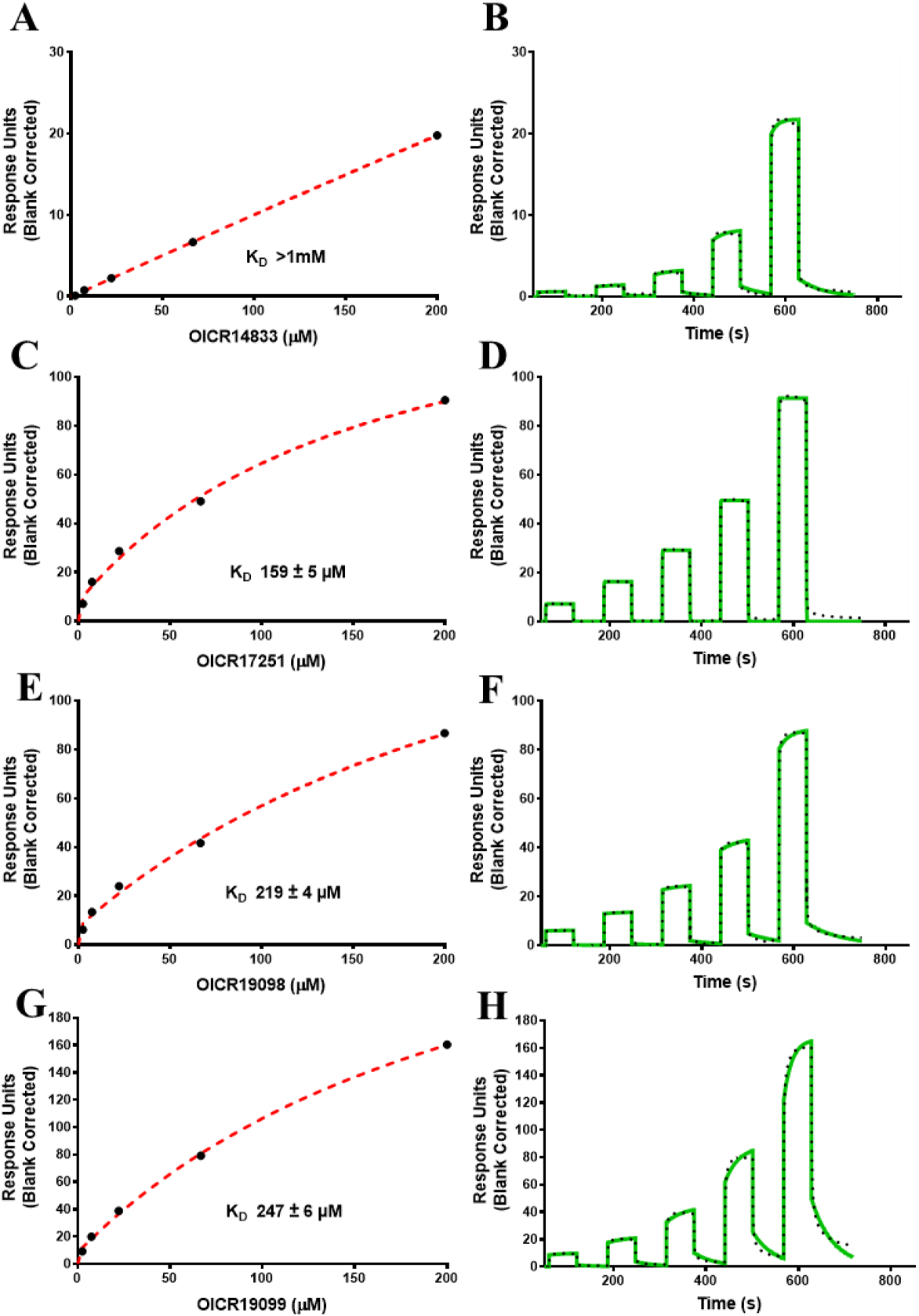
Orthogonal confirmation of ligand binding by SPR. Representative binding curves and sensorgrams for (A, B) OICR14833, (C, D) OICR17251, (E, F) OICR19098 and (G, H) OICR19099 are presented. For each compound, the steady state response (black circles; A, C, E and G) with the steady state 1:1 binding model fitting (red dashed line), and the sensorgrams (solid green; B, D, F, H) are shown with the kinetic fit (black dots). Experiments were performed in triplicate, and data are shown as the mean ± standard deviation. The K_D_ values are presented on each figure.

**Figure 4:**
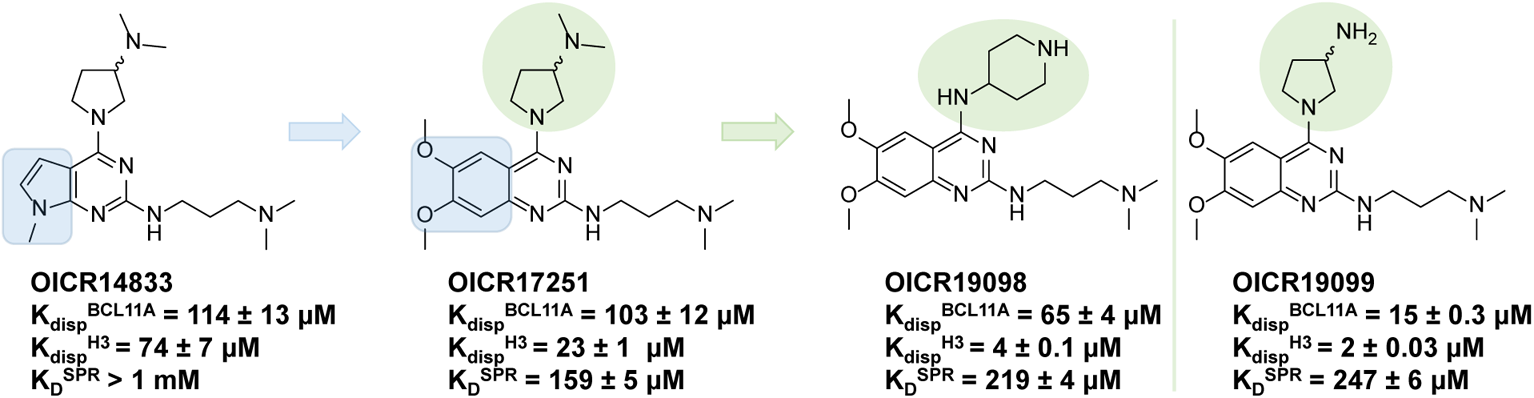
Evolution of SAR from initial screening hit. Design strategy for the hit expansion: Replacement of deazapurine core of OICR14833 (K_disp_ of 114 ± 13 μM and 74 ± 7 μM against RBBP4-BCL11A and RBBP4-H3, respectively) with an ortho-dimethoxy quinazoline in OICR1725, resulted in increased potency with a K_disp_ of 103 ± 12 μM and 23 ± 1 μM against RBBP4-BCL11A and RBBP4-H3, respectively. Furthermore, replacement of the dimethylaminopyrrolidine moiety at position 4 of the quinazoline core of OICR17251 with a piperidin-4-amine in OICR19098 further increased potency with a K_disp_ of 65 ± 4 μM and 4 ± 0.1 μM against RBBP4-BCL11A and RBBP4-H3, respectively. Finally, replacement of the dimethylaminopyrrolidine moiety at position 4 of the quinazoline core of OICR17251 with an aminopyrrolidine in OICR19099 further increased potency with a K_disp_ of 15 ± 3 μM and 2 ± 0.03 μM against RBBP4-BCL11A and RBBP4-H3, respectively.

OICR14833 (**Fig. 4**) was selected from the structural cluster for further characterization and optimization by SAR studies. More than 40 analogs of OICR14833 were synthesized and surveyed. Specifically, we examined cases where the deazapurine core was replaced by other fused heteroaromatic scaffolds. Interestingly, diaminoquinazolines and diaminoquinolines were found to be active in the peptide displacement assay. Although not significantly more potent, all 40 analogs were tested in our optimized peptide-displacement assay. Out of 40 compounds, 23 displaced both BCL11A (2-16) and H3 (1-21), suggesting binding to the top pocket of RBBP4 (**Table 1**). Out of the 23 hits, 6 showed weak affinities in the high micromolar range (K_disp_ > 80 μM against RBBP4-H3) (**Table 1**). We also ratified their specificity by checking their displacement against MTA1 and verified that all compounds only bind to the top pocket, and none displaced MTA1 peptide from the RBBP4 side pocket. An *ortho*-dimethoxy quinazoline, OICR17251 (**Fig. 4**) was one of the most potent ligands with a K_disp_ of 103 ± 12 μM and 23 ± 1 μM against RBBP4-BCL11A (**Fig. 2D**) and RBBP4-H3 (**Fig. 2E**), respectively. The binding of OICR17251 to RBBP4 was also validated by SPR (K_D_ of 159 ± 5 μM) (**Fig. 3C & 3D**).

**Table 1.**
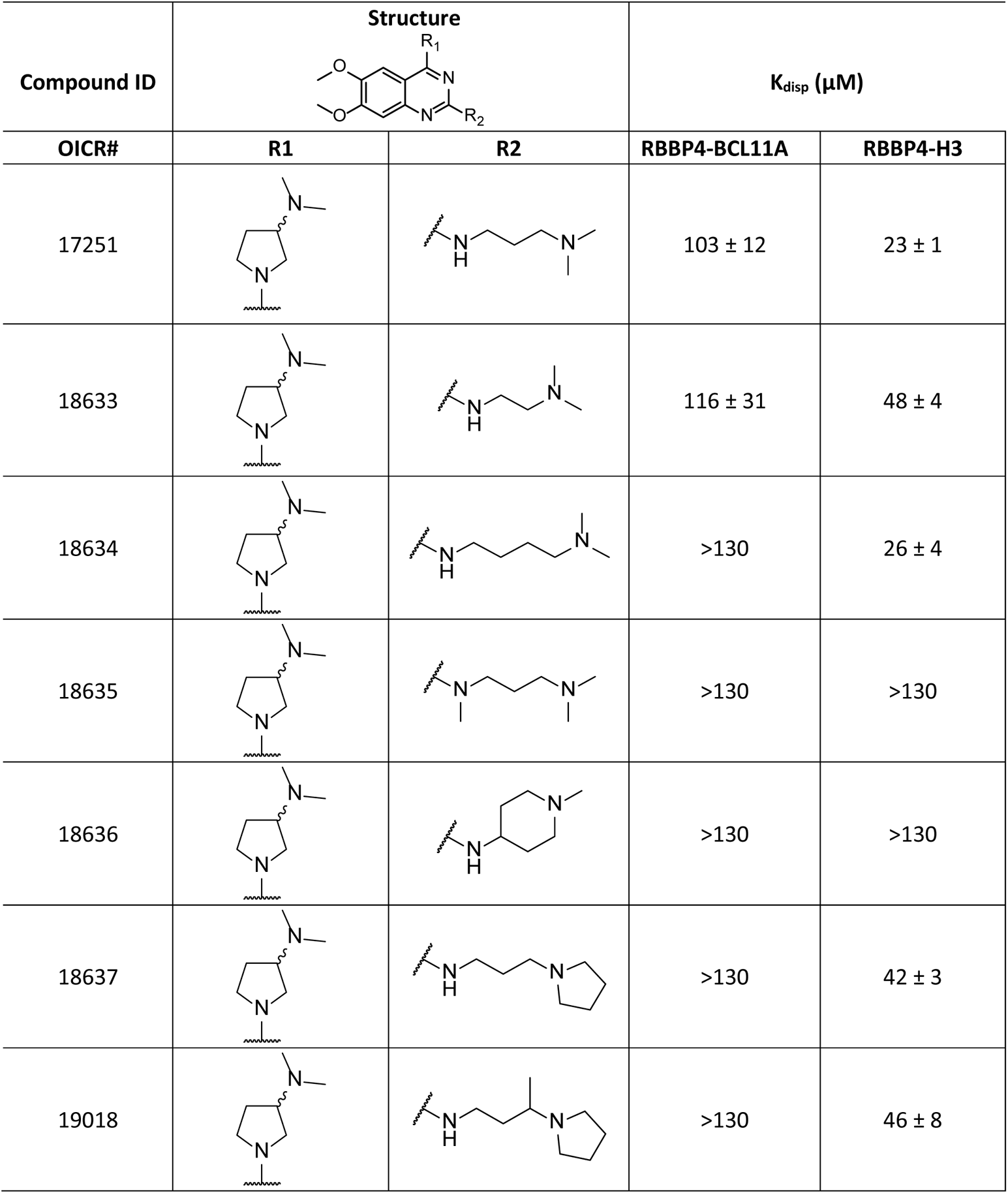
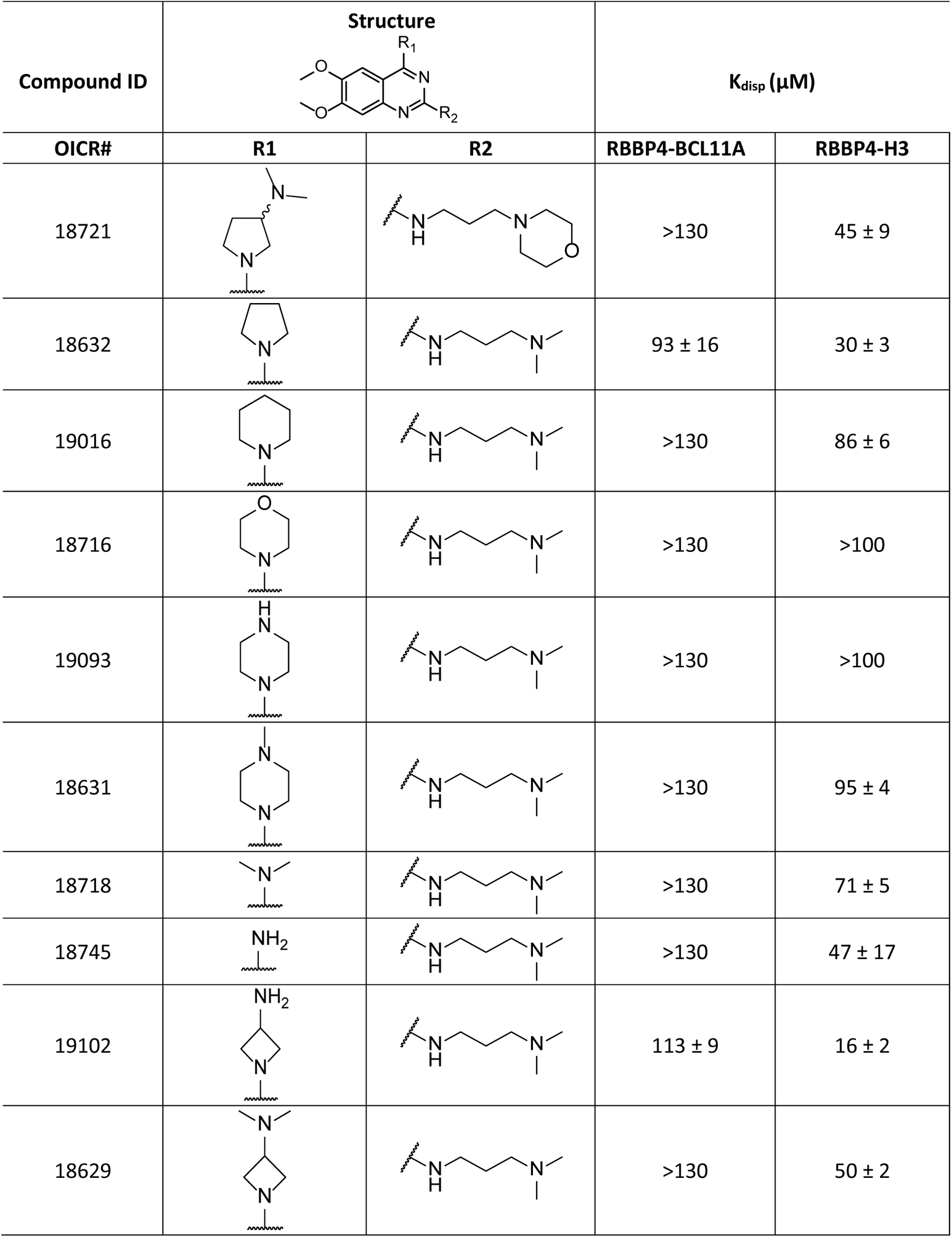
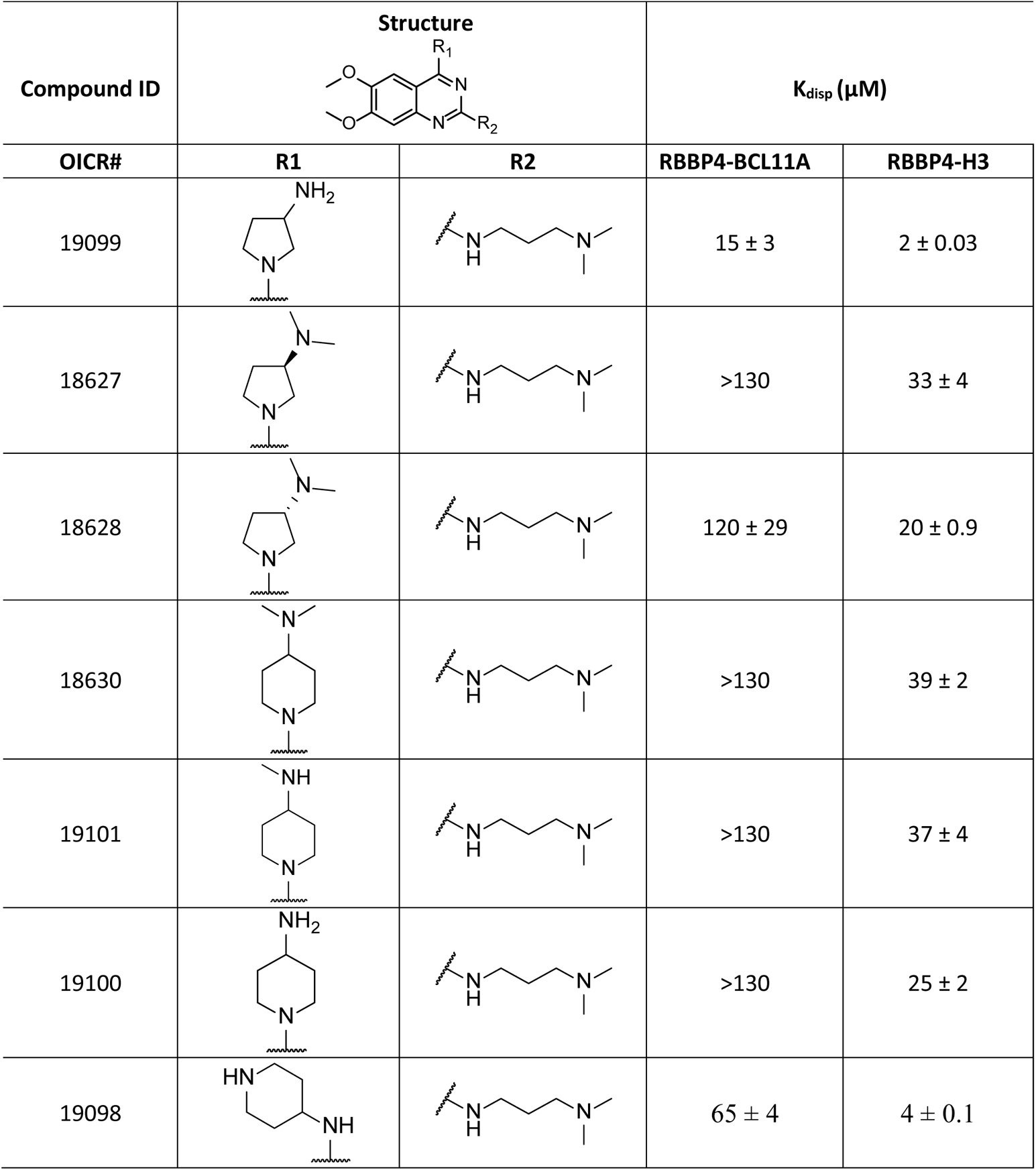
Characterization of compounds of interest. K_disp_ values were determined using BCL11A (2-16), and H3 (1-21) peptides. All values represent the mean of three independent experiments ± standard deviation.

| Compound ID | Structure | | $K_{\text{disp}}$ ($\mu\text{M}$) | |
| --- | --- | --- | --- | --- |
|  | R1 | R2 | RBBP4-BCL11A | RBBP4-H3 |
| 17251 | | | 103 ± 12 | 23 ± 1 |
| 18633 | | | 116 ± 31 | 48 ± 4 |
| 18634 | | | >130 | 26 ± 4 |
| 18635 |  |  | >130 | >130 |
| 18636 |  |  | >130 | >130 |
| 18637 | | | >130 | 42 ± 3 |
| 19018 | | | >130 | 46 ± 8 |

Table 1. Continued
| Compound ID | Structure | | $K_{\text{disp}}$ ($\mu\text{M}$) | |
| --- | --- | --- | --- | --- |
|  | R1 | R2 |  |  |
| 18721 |  |  | >130 | 45 ± 9 |
| 18632 |  |  | 93 ± 16 | 30 ± 3 |
| 19016 |  |  | >130 | 86 ± 6 |
| 18716 |  |  | >130 | >100 |
| 19093 |  |  | >130 | >100 |
| 18631 |  |  | >130 | 95 ± 4 |
| 18718 |  |  | >130 | 71 ± 5 |
| 18745 |  |  | >130 | 47 ± 17 |
| 19102 |  |  | 113 ± 9 | 16 ± 2 |
| 18629 |  |  | >130 | 50 ± 2 |

Table 1. Continued
| Compound ID | Structure | | $K_{\text{disp}}$ ($\mu\text{M}$) | |
| --- | --- | --- | --- | --- |
|  | R1 | R2 |  |  |
| 19099 |  |  | 15 ± 3 | 2 ± 0.03 |
| 18627 |  |  | >130 | 33 ± 4 |
| 18628 |  |  | 120 ± 29 | 20 ± 0.9 |
| 18630 |  |  | >130 | 39 ± 2 |
| 19101 |  |  | >130 | 37 ± 4 |
| 19100 |  |  | >130 | 25 ± 2 |
| 19098 |  |  | 65 ± 4 | 4 ± 0.1 |

To confirm the competitive mode of action observed in the fluorescence polarization assay, we solved the crystal structure of compound OICR17251 in complex with RBBP4 (PDB: 7M40). The antagonist occupies the central cavity of the WDR domain, which is also exploited by FOG1^17^, H3^18^ and PHF6^16^ peptides (**Fig. 5, Suppl. Fig. 5**). While ambiguous, the electron density is compatible with a binding pose where the ligand is engaged in a set of interactions that recapitulate those formed by a FOG-1 peptide previously co-crystallized with RBBP4 ^17^. In this putative model, the aromatic system of OICR17251 is anchored in the cavity, sandwiched in a π-stacking arrangement between Tyr-181 and Phe-321, and engaged in a hydrogen-bond with Glu-231, recapitulating interactions made by Arg-4 of FOG1^17^. The terminal dimethyl ammonium group of OICR17251 mimics FOG1 Arg-3 in making an electrostatic interaction with Glu-319. Electron density for the dimethylaminopyrrolidine group from the pyrrolidine group is missing, suggesting that this group can adopt multiple conformational states in the context of the bound molecule. These results suggest a structural mechanism where OICR17251 directly competes with peptides exploiting the top pocket of RBBP4.

**Figure 5:**
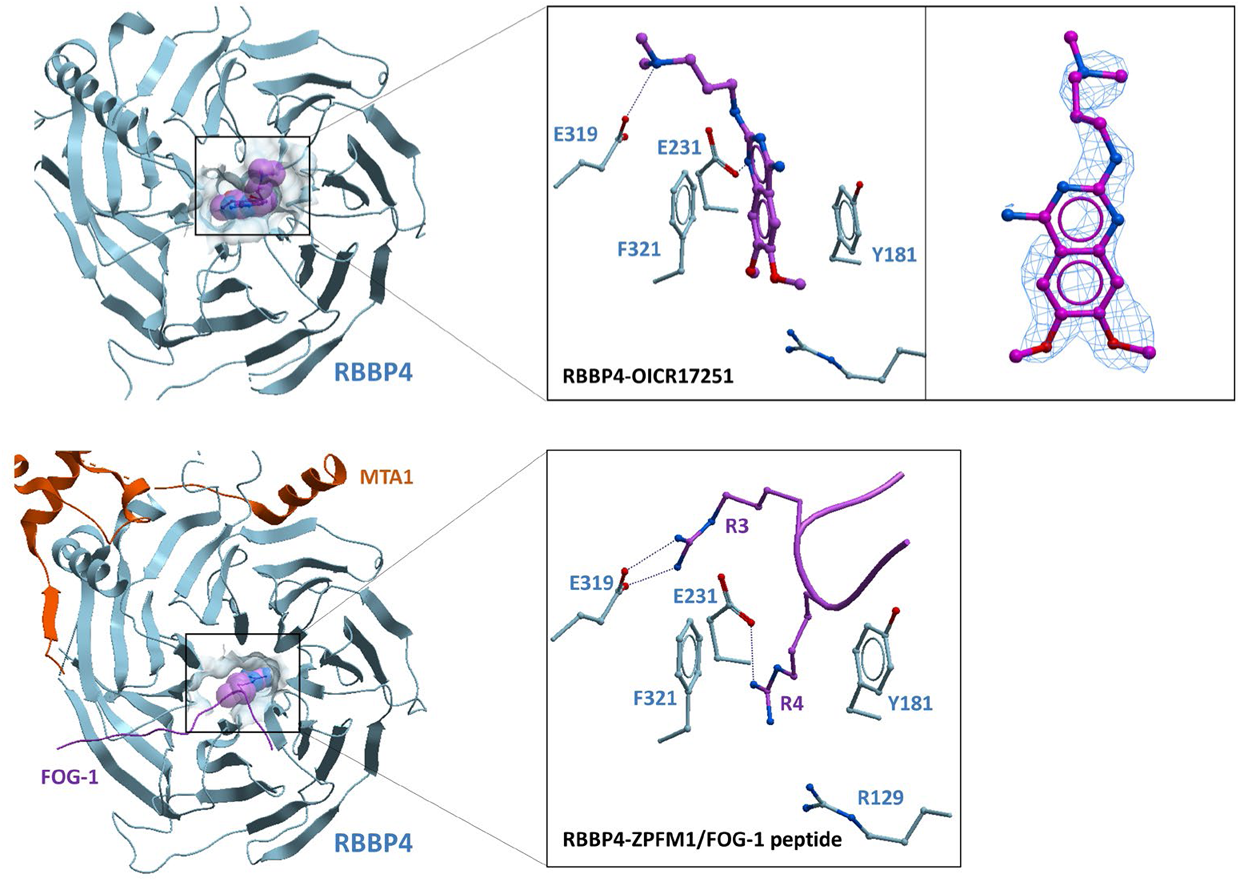
Crystal structure of the RBBP4 in complex with inhibitor OICR17251. Top: OICR17251 occupies the central pocket of the RBBP4 WDR domain. Insert: Probable binding pose (left), suggested by an incomplete mFo-DFc electron density contoured at level 2.5 with Molsoft’s ICM (right). Bottom: The central pocket is exploited by a FOG-1 peptide (purple, PDB code 2XU7), while a MTA1 peptide (orange) interacts with the side of the WDR (PDB code 6G16).

Crystallization of OICR17251-bound RBBP4, allowed us to gain additional insight into the mode of binding. The solvent-exposed dimethylamine group of the aliphatic chain of OICR17251 is predicted to be protonated at physiological pH. The dimethylamine group (stemming from the pyrrolidine) could not be fully resolved in the X-ray structure, due to the presence of both enantiomers, however, its position indicates that there could be a second electrostatic hydrogen bonding interaction between the side chain carboxylate of Glu-319 and the protonated dimethylamine. The dimethoxy moiety of the ligand occupies the bottom of the pocket, in proximity to the guanidinium group of Arg-129.

We initially focused on the installation of flexible and basic substituents to increase the interaction of the compound with the surrounding negatively charged environment (**Fig. 4**). In particular, incorporation in OICR19098 and OICR19099 of a piperidine-4-amine and a 3-amino-pyrrolidine substitution, respectively, provided our best hits in terms of biochemical potency, which may be driven by electrostatic interactions with E395 (**Fig. 4**). OICR19098 showed K_disp_ of 65 ± 4 μM and 4 ± 0.1 μM against RBBP4-BCL11A and RBBP4-H3, respectively (**Fig. 2G, 2H & 2I**), and binding to RBBP4 with SPR K_D_ of 219 ± 4 μM (**Fig. 3E & 3F**). OICR19099 showed improved K_disp_ of 15 ± 0.3 μM and 2 ± 0.03 μM against RBBP4-BCL11A and RBBP4-H3, respectively (**Fig. 2J, 2K & 2L**), and binding to RBBP4 with SPR K_D_ of 247 ± 6 μM (**Fig. 3G and 3H**).

As already stated, the installation of the two methoxy moieties at positions 6 and 7 were found to be central to the development of a potent inhibitor (**Fig. 4**). Analogues devoid of the methoxy groups at these positions were found to be less potent and completely insoluble in water. Substitution with different methoxy bioisosteres lead to a substantial decrease in potency (data not shown). Computational analysis showed that this moiety freezes the conformation of the inhibitor, projecting the methyl groups away from each other in the same plane and making the oxygen lone pairs face each other. Having those two lone pairs in that conformation gives the compound an improved hydrogen bonding ability with Arg-129.

## Conclusion

In summary, we report the discovery of the first RBBP4 antagonists through medium-throughput screening using novel peptide-displacement assays. Conducting small-scale SAR studies indicated that the aromatic system of the ligand occupies the top binding pocket of RBBP4 through non-covalent and electrostatic interactions. We believe that the ligands reported here, and the co-crystal structure deposited in the PDB (7M40) through this study will provide insight to accelerate optimization of these types of antagonists through more comprehensive SAR studies.

## EXPERIMENTAL SECTION

### Protein Expression and Purification

Full-length human RBBP4 (residues 1–425) was cloned into the baculovirus expression vector pFBOH-LIC (GenBank EF456740) encoding an N-terminal His6 tag, and a tobacco etch virus protease cleavage site. The protein was expressed in *Sf9* cells infected with baculovirus using the Bac-to-Bac^TM^ expression methodology (Invitrogen). Harvested *Sf9* cells were re-suspended in 20 mM Tris-HCl (pH 7.5) containing 500 mM NaCl, 5 mM imidazole, 5% glycerol and 1x protease inhibitor cocktail (100x protease inhibitor stock in 70% ethanol; 0.25 mg/ml Aprotinin, 0.25 mg/ml Leupeptin, 0.25 mg/ml Pepstatin A and 0.25 mg/ml E-64) or Roche cOmplete^TM^ EDTA-free protease inhibitor cocktail tablet (Millipore-Sigma). NP-40 (final concentration of 0.5%) and 15 µl Benzonase nuclease (In-House) was added to the cell suspension, followed by 30 min of rotation and further sonication at frequency of 7 (10” on/10” off) for 4 min (Sonicator 3000, Misoni). The crude extract was clarified by high-speed centrifugation (60 min at 36,000 g at 4 ℃) using a Beckman Coulter Centrifuge. Clarified lysate was then incubated for one hour at 4 °C with Ni-NTA (Qiagen) in batch adsorption format. The resin was then transferred to an open column (Bio-Rad) where it was washed with binding buffer followed by washing buffer containing 20 mM Tris-HCl (pH 7.5), 500 mM NaCl, 5% glycerol, 5 mM imidazole and 30 mM imidazole, respectively. Protein was then eluted with 20 mM Tris-HCl (pH 7.5), 500 mM NaCl, 5% glycerol and 250 mM imidazole. For further purification, the eluate was loaded onto a Resource Q ion exchange column (GE Healthcare) followed by Gel Filtration on a HiLoad Superdex 75 26/60 (GE Healthcare) column pre-equilibrated in 20 mM Tris (pH 8), 200 mM NaCl, 5% glycerol, 1 mM TCEP. Fractions containing pure protein were pooled, concentrated and flash frozen by immersion in liquid nitrogen, and kept at −80 °C.

### Crystallization

Purified RBBP4 (1-425) at a concentration of 13 mg/mL was buffer exchanged to 10 mM Tris pH 8, 100 mM NaCl and incubated with 10 mM OICR17251 for 1 h on ice. Co-crystals were obtained by the hanging-drop vapor-diffusion method at 18 °C. Each drop included 2 μL of the RBBP4-OICR17251 complex and 2 μL of reservoir solution which contained 23% PEG3350, 0.2 M NH_4_OAc, 0.1 M BisTris pH 5.5. The crystals were formed after 5 days. Using a cryoprotectant solution containing the reservoir solution supplemented with 10% (v/v) ethylene glycol, single crystals were flash frozen in liquid nitrogen and stored.

### Data Collection, Structure Determination, and Refinement

X-ray diffraction data for RBBP4-OICR17251 co-crystals were collected at 100 K at the Argonne National Laboratory Advanced Photon Source beamline 19-ID with a wavelength of 0.979180 Å from Pilatus3 X 6M detector. Diffraction data were processed and scaled with the XDS^28^ and AIMLESS^29^. The structure was solved by molecular replacement using human RBBP4 (PDB ID: 6BW3) as the search model with the program Phaser^30^ and constraints for compound were generated by GRADE. The structural models were refined using REFMAC^31^ and manually checked with COOT^32^ for model building and visualization. Data collection and refinement statistics are presented **in Suppl. Table 3**.

### Fluorescence polarization assay

All experiments were performed in a total volume of 20 μL in 384-well black, low volume, round-bottom polypropylene microplates (Greiner). To obtain the K_D_ values for RBBP4 interaction with the peptides, protein was titrated in a reaction mixture in the optimized buffer composition and FITC-peptide (**Suppl. Table 1)**. The reaction mixture in the plate was incubated at room temperature for 30 min. Fluorescence polarization was measured using a Biotek Synergy H1 microplate reader (Biotek) at excitation and emission wavelengths of 485 and 528 nm, respectively. The data were analyzed with GraphPad Prism 8.2.0 (GraphPad Software Inc., San Diego).

### Fluorescence polarization-based peptide displacement assay

All the K displacement (K_disp_) values were measured using fluorescence polarization-based peptide displacement assays performed using the unlabeled peptides with identical sequence as the FITC-labeled peptide or ligands (**Suppl. Table 2)**. The unlabeled peptide was titrated in the optimized buffer containing FITC-labeled peptide and protein, while ligands were titrated in optimized buffer containing 2% (v/v) DMSO for 2-fold dilution titration series. Here, a protein concentration below saturation (corresponding to 80% of polarization signal in the binding assay) was used. Experiments were performed in 384-well black, low volume, Greiner plates, and the FP was measured after incubation at room temperature for 30 min. The data were analyzed with GraphPad Prism, K_disp_ and Hill slopes were calculated.

### Surface Plasmon Resonance (SPR)

SPR studies were performed using a Biacore T200 (GE Health Sciences) at 25 °C. Biotinylated RBBP4 was captured onto a flow cell of a streptavidin-conjugated SA chip at approximately 4000 response units (RU) according to manufacturer’s protocol while another flow cell was left empty for reference subtraction. Serial dilutions of compounds (dilution factor of 0.33 was used to yield 5 concentrations) were prepared with 200 μM as the highest compound concentration in HBS-P buffer (10 mM Hepes pH 7.4, 150 mM NaCl, 0.005% Tween) and 2% DMSO. Kinetic determination experiments were performed using single-cycle kinetics with 60 s on time, and off time of 180 s at a flow rate of 40 μL min^−1^. Buffer containing 2% DMSO was used for blank injections; and buffers containing 1 to 3 % DMSO were used for buffer corrections. K_D_ values were calculated by using steady state affinity fitting and the Biacore T200 Evaluation software.

### Chemical Synthesis and Compound Characterization

#### General procedure A

To 2,4-dichloro-6,7-dimethoxyquinazoline (150 mg, 0.579 mmol, 1 equiv.) in tetrahydrofuran (THF, ∼0.1 molar, 5 mL) were added *N*,*N*-diisopropylethylamine (DIPEA, 0.302 mL, 1.737 mmol, 3 equiv.) and the corresponding primary or secondary amine (1.158 mmol, 2 equiv.) at room temperature. The solution was stirred for 16 h. Completion of the reaction was monitored by LCMS and the volatiles were evaporated. CH_2_Cl_2_ (10 mL) and 10% K_2_CO_3_ solution were added (10 mL), upon combination of organic and aqueous solutions, an insoluble precipitate appears, which floats between the two layers. The organic layer was separated, and the aqueous layer extracted with CH_2_Cl_2_ (2 x 10 mL). The organic phases were combined alongside the insoluble precipitate. Note: if emulsion appears, the container was centrifuged at 500 RPM for 5 min. The remaining CH_2_Cl_2_ was evaporated to obtain a yellow-brown semisolid that was suspended in water and freeze dried for 2 days to obtain the final solid product that was used in the following step without further purification.

#### General procedure B

A suspension of palladium(II) acetate (2.65 mg, 0.012 mmol, 0.05 equiv.) and 2,2’-*bis*(diphenylphosphino)-1,1’-binaphthalene (14.72 mg, 0.024 mmol, 0.1 equiv.) in 1,4-dioxane (∼0.3 molar, 0.800 mL) were gently warmed with a heat gun with vigorous stirring resulting in a clear red-orange solution. This solution was added to a mixture of the 2-chloroquinazolin-4-amine (Procedure A, 1 equiv.), the desired alkylamine (3 equiv.) and Cs_2_CO_3 anhyd._ (2 equiv.). The reaction mixture was heated to 130 °C under N_2_ atmosphere for 16 h. The reaction was monitored by LCMS. The volatiles in the reaction mixture were evaporated and the crude was diluted with water (5 mL).

Purification was performed via RP Biotage column, (98-2% to 80-20%, H_2_O (0.1% formic acid); ACN (0.1% formic acid); using a gradient for 15 min and isocratic for 5 min using a double RP 12 g column), direct loading using acetonitrile (ACN) as solvent. After evaporation of the desired fraction the (protected) product was obtained as the formic acid salt. The protecting group was removed (when a protecting group is present) by dissolving the sample in CH_2_Cl_2_ and adding 0.75-1.00 mL of trifluoroacetic acid (TFA) and stirring for 1 h. The volatiles were removed, and the product was desalted by passing it through a Biotage^®^ SCX-2 cation exchange resin (0.5 g) washing with MeOH (10 mL) and eluting with NH_4_OH (3% w/v in MeOH, 10 mL). The solvent was removed, and product was suspended in water and freeze dried for 2 days to obtain the final product.

### 1-(2-Chloro-6,7-dimethoxyquinazolin-4-yl)-N,N-dimethylpyrrolidin-3-amine

The title compound was synthesized according to the general procedure **A**. 2,4-Dichloro-6,7-dimethoxyquinazoline (310 mg, 1.197 mmol), tetrahydrofuran (2 mL), *N,N*-diisopropylethylamine (155 mg, 1.197 mmol) and 3-(dimethylamino)pyrrolidine (137 mg, 1.197 mmol). A white powder was obtained 325 mg (80% yield).^1^H NMR (500 MHz, CDCl_3_) δ 7.42 (s, 1H), 7.14 (s, 1H), 4.14 – 4.04 (m, 2H), 4.00 – 3.93 (m, 7H), 3.81 (dd, *J* = 10.4, 8.4 Hz, 1H), 2.86 – 2.76 (m, 1H), 2.35 (s, 6H), 2.29 – 2.20 (m, 1H), 2.03 – 1.92 (m, 1H). MS (ESI): m/z = 337.5 [M + H]^+^.

### *N*^1^-(4-(3-(Dimethylamino)pyrrolidin-1-yl)-6,7-dimethoxyquinazolin-2-yl)-*N*^3^,*N*^3^-dimethylpropane-1,3-diamine (OICR17251)

The title compound was synthesized according to the general procedure **B**. Palladium(II) acetate (1.333 mg, 5.94 µmol), 2,2’-*bis*(diphenylphosphino)-1,1’-binaphthalene (7.39 mg, 0.012 mmol), 1,4-dioxane (1 mL), 1-(2-chloro-6,7-dimethoxyquinazolin-4-yl)-*N*,*N*-dimethylpyrrolidin-3-amine (40 mg, 0.119 mmol), 3-(dimethylamino)-1-propylamine (18.20 mg, 0.178 mmol) and Cs_2_CO_3 anhyd_ (77 mg, 0.238 mmol). A yellow oil was obtained 40 mg (83% yield). ^1^H NMR (500 MHz, CDCl_3_) δ 7.28 (s, 1H), 6.86 (s, 1H), 5.04 (s, 1H), 4.04 – 3.98 (m, 1H), 3.95 – 3.88 (m, 5H), 3.86 (s, 3H), 3.77 – 3.68 (m, 1H), 3.54 – 3.43 (m, 2H), 2.78 – 2.70 (m, 1H), 2.39 – 2.34 (m, 2H), 2.32 (s, 6H), 2.22 (s, 6H), 2.20 – 2.16 (m, 1H), 1.93 – 1.84 (m, 1H), 1.81 – 1.74 (m, 2H); LCMS Column 1, RT = 0.88 min, MS (ESI): m/z = 403.56 [M + 1]^+^, Purity (UV^254^) = 100%; HRMS (ESI) for C_21_H_35_N_6_O_2_ [M + H]^+^: m/z = calcd, 403.2821; found, 403.2819.

### *N*^1^-(4-(3-(Dimethylamino)pyrrolidin-1-yl)-6,7-dimethoxyquinazolin-2-yl)-*N*^2^,*N*^2^-dimethylethane-1,2-diamine (OICR18633)

The title compound was synthesized according to the general procedure **B**. Palladium(II) acetate (1.616 mg, 7.20 µmol), 2,2’-*bis*(diphenylphosphino)-1,1’-binaphthalene (8.97 mg, 0.014 mmol), 1,4-dioxane (480 µL), 1-(2-chloro-6,7-dimethoxyquinazolin-4-yl)-*N,N*-dimethylpyrrolidin-3-amine (48.5 mg, 0.144 mmol), *N,N*-dimethylethylenediamine 95% (38.1 mg, 0.432 mmol) and Cs_2_CO_3 anhyd_ (94 mg, 0.288 mmol). A yellow powder was obtained 35 mg (63% yield). ^1^H NMR (500 MHz, MeOD) δ 7.44 (s, 1H), 6.84 (s, 1H), 4.12 – 4.06 (m, 2H), 4.01 (td, *J* = 10.6, 6.6 Hz, 1H), 3.92 (s, 3H), 3.87 (s, 3H), 3.75 (dd, *J* = 10.5, 8.9 Hz, 1H), 3.58 (t, *J* = 6.7 Hz, 2H), 2.92 (td, *J* = 8.8, 4.3 Hz, 1H), 2.67 (t, *J* = 6.7 Hz, 2H), 2.38 (s, 6H), 2.37 (s, 6H), 2.33 – 2.29 (m, 1H), 1.95 – 1.89 (m, 1H); LCMS Column 1, RT = 0.80 min, MS (ESI): m/z = 389.48 [M + 1]^+^, Purity (UV^254^) = 100%; HRMS (ESI) for C_20_H_33_N_6_O_2_ [M + H]^+^: m/z = calcd, 389.2665; found, 389.2660.

### *N*^1^-(4-(3-(Dimethylamino)pyrrolidin-1-yl)-6,7-dimethoxyquinazolin-2-yl)-*N*^4^,*N*^4^-dimethylbutane-1,4-diamine (OICR18634)

The title compound was synthesized according to the general procedure **B.** Palladium(II) acetate (1.660 mg, 7.39 µmol), 2,2’-*bis*(diphenylphosphino)-1,1’-binaphthalene (9.21 mg, 0.015 mmol), 1,4-dioxane (493 µL), 1-(2-chloro-6,7-dimethoxyquinazolin-4-yl)-*N*,*N*-dimethylpyrrolidin-3-amine (49.8 mg, 0.148 mmol), 4-dimethylaminobutylamine (51.5 mg, 0.444 mmol) and Cs_2_CO_3 anhyd_ (96 mg, 0.296 mmol). A yellow semi-solid was obtained 26 mg (38% yield). ^1^H NMR (500 MHz, MeOD) δ 7.45 (s, 1H), 6.87 (s, 1H), 4.14 – 4.06 (m, 2H), 4.01 (td, *J* = 10.7, 6.6 Hz, 1H), 3.92 (s, 3H), 3.88 (s, 3H), 3.78 – 3.74 (m, 1H), 3.45 (dd, *J* = 8.0, 4.4 Hz, 2H), 2.95 – 2.90 (m, 1H), 2.51 (d, *J* = 7.2 Hz, 2H), 2.37 (s, 6H), 2.35 (s, 6H), 1.98 – 1.88 (m, 2H), 1.64 (dd, *J* = 9.3, 5.6 Hz, 4H); LCMS Column 1, RT = 0.84 min, MS (ESI): m/z = 417.54 [M + 1]^+^, Purity (UV^254^) = 94%; HRMS (ESI) for C_22_H_37_N_6_O_2_ [M + H]^+^: m/z = calcd, 417.2978; found, 417.2975.

### *N*^1^-(4-(3-(Dimethylamino)pyrrolidin-1-yl)-6,7-dimethoxyquinazolin-2-yl)-*N*^1^,*N*^3^,*N*^3^-trimethylpropane-1,3-diamine (OICR18635)

The title compound was synthesized according to the general procedure **B.** Palladium(II) acetate (1.750 mg, 7.79 µmol) and 2,2’-*bis*(diphenylphosphino)-1,1’-binaphthalene (9.71 mg, 0.016 mmol) in 1,4-dioxane (520 µL), 1-(2-chloro-6,7-dimethoxyquinazolin-4-yl)-*N,N*-dimethylpyrrolidin-3-amine (52.5 mg, 0.156 mmol), *N,N,N’*-trimethyl-1,3-propanediamine (54.3 mg, 0.468 mmol) and Cs_2_CO_3 anhyd_ (102 mg, 0.312 mmol). An orange semi-solid was obtained 23 mg (35% yield).^1^H NMR (500 MHz, MeOD) δ 7.43 (s, 1H), 6.95 (s, 1H), 4.13 – 4.04 (m, 2H), 4.01 (dd, *J* = 10.4, 6.6 Hz, 1H), 3.92 (d, *J* = 4.9 Hz, 3H), 3.87 (s, 3H), 3.79 – 3.74 (m, 1H), 3.71 (d, *J* = 3.0 Hz, 1H), 3.18 (s, 3H), 2.93 (dd, *J* = 15.0, 7.5 Hz, 1H), 2.53 (s, 2H), 2.38 (d, *J* = 4.9 Hz, 12H), 2.33 – 2.27 (m, 1H), 1.98 – 1.86 (m, 4H); LCMS Column 1, RT = 0.82 min, MS (ESI): m/z = 417.61 [M + 1]^+^, Purity (UV^254^) = 100%; HRMS (ESI) for C_22_H_37_N_6_O_2_ [M + H]^+^: m/z = calcd, 417.2978; found, 417.2975.

### 4-(3-(Dimethylamino)pyrrolidin-1-yl)-6,7-dimethoxy-N-(1-methylpiperidin-4-yl)quinazolin-2-amine (OICR18636)

The title compound was synthesized according to the general procedure **B**. Palladium(II) acetate (1.700 mg, 7.57 µmol), 2,2’-*bis*(diphenylphosphino)-1,1’-binaphthalene (9.43 mg, 0.015 mmol), 1,4-dioxane (505 µL), 1-(2-chloro-6,7-dimethoxyquinazolin-4-yl)-*N,N*-dimethylpyrrolidin-3-amine (51 mg, 0.151 mmol), 4-amino-1-methylpiperidine (51.9 mg, 0.454 mmol) and Cs_2_CO_3 anhyd_ (99 mg, 0.303 mmol). A white powder was obtained 17 mg (27% yield). ^1^H NMR (500 MHz, MeOD) δ 7.49 (s, 1H), 6.92 (s, 1H), 4.20 – 4.09 (m, 2H), 4.09 – 4.03 (m, 1H), 3.95 (s, 3H), 3.91 (s, 3H), 3.79 (dd, *J* = 20.1, 11.2 Hz, 1H), 3.14 – 2.86 (m, 4H), 2.50 (s, 2H), 2.41 (t, *J* = 4.5 Hz, 6H), 2.37 – 2.30 (m, 1H), 2.16 (s, 1H), 1.96 (d, *J* = 8.1 Hz, 2H), 1.74 (d, *J* = 10.1 Hz, 1H). Two sets of rotamers; LCMS Column 1, RT = 0.87 min, MS (ESI): m/z = 415.58 [M + 1]^+^, Purity (UV^254^) = 100%; HRMS (ESI) for C_22_H_35_N_6_O_2_ [M + H]^+^: m/z = calcd, 415.2821; found, 415.2818.

### 4-(3-(Dimethylamino)pyrrolidin-1-yl)-6,7-dimethoxy-N-(3-(pyrrolidin-1-yl)propyl)quinazolin-2-amine (OICR18637)

The title compound was synthesized according to the general procedure **B**. Palladium(II) acetate (1.666 mg, 7.42 µmol), 2,2’-*bis*(diphenylphosphino)-1,1’-binaphthalene (9.24 mg, 0.015 mmol), 1,4-dioxane (495 µL), 1-(2-chloro-6,7-dimethoxyquinazolin-4-yl)-*N,N*-dimethylpyrrolidin-3-amine (50 mg, 0.148 mmol), 1-(3-aminopropyl)pyrrolidine (57.1 mg, 0.445 mmol) and Cs_2_CO_3 anhyd_ (97 mg, 0.297 mmol). A yellow semi-solid was obtained 17 mg (27% yield).^1^H NMR (500 MHz, MeOD) δ 6.70 (d, *J* = 3.7 Hz, 1H), 6.38 (d, *J* = 3.7 Hz, 1H), 3.93 – 3.90 (m, 4H), 3.59 (s, 3H), 3.42 (t, *J* = 6.8 Hz, 2H), 2.56 – 2.53 (m, 4H), 2.48 – 2.44 (m, 2H), 2.34 (s, 3H), 2.27 (s, 6H), 1.85 – 1.79 (m, 2H); LCMS Column 1, RT = 0.94 min, MS (ESI): m/z = 429.57 [M + 1]^+^, Purity (UV^254^) = 100%; HRMS (ESI) for C_23_H_37_N_6_O_2_ [M + H]^+^: m/z = calcd, 429.2978; found, 429.2972.

### 4-((S)-3-(Dimethylamino)pyrrolidin-1-yl)-6,7-dimethoxy-N-(3-(pyrrolidin-1-yl)butyl)quinazolin-2-amine (OICR19018)

The title compound was synthesized according to the general procedure **B**. Palladium(II) acetate (1.700 mg, 7.57 µmol), 2,2’-*bis*(diphenylphosphino)-1,1’-binaphthalene (9.43 mg, 0.015 mmol), 1,4-dioxane (505 µL), (S)-1-(2-chloro-6,7-dimethoxyquinazolin-4-yl)-*N,N*-dimethylpyrrolidin-3-amine (505 µl, 0.151 mmol), 3-(1-pyrrolidinyl)-1-butanamine (64.6 mg, 0.454 mmol) and Cs_2_CO_3_ anhyd (99 mg, 0.303 mmol). A yellow semi-solid was obtained 26 mg (40% yield). ^1^H NMR (500 MHz, MeOD) δ 7.43 (s, 1H), 6.82 (s, 1H), 4.13 – 3.96 (m, 4H), 3.91 (s, 3H), 3.87 (s, 3H), 3.78 – 3.70 (m, 1H), 3.58 – 3.47 (m, 1H), 3.47 – 3.38 (m, 1H), 2.96 – 2.85 (m, 1H), 2.70 (br s, 4H), 2.63 – 2.48 (m, 1H), 2.37 (s, 6H), 2.33 – 2.25 (m, 1H), 2.12 – 1.98 (m, 1H), 1.98 – 1.86 (m, 1H), 1.86 – 1.75 (m, 4H), 1.64 (dt, *J* = 8.9, 4.4 Hz, 1H), 1.26 – 1.19 (m, 3H); LCMS Column 1, RT = 1.02 min, MS (ESI): m/z = 443.69 [M + 1]^+^, Purity (UV^254^) = 100%; HRMS (ESI) for C_24_H_39_N_6_O_2_ [M + H]^+^: m/z = calcd, 443.3134; found, 443.3125.

### 4-(3-(Dimethylamino)pyrrolidin-1-yl)-6,7-dimethoxy-N-(3-morpholinopropyl)quinazolin-2-amine (OICR18721)

The title compound was synthesized according to the general procedure **B**. Palladium(II) acetate (1.700 mg, 7.57 µmol), 2,2’-*bis*(diphenylphosphino)-1,1’-binaphthalene (9.43 mg, 0.015 mmol), 1,4-dioxane (505 µL), 1-(2-chloro-6,7-dimethoxyquinazolin-4-yl)-*N,N*-dimethylpyrrolidin-3-amine (51 mg, 0.151 mmol), 3-morpholinopropylamine (65.5 mg, 0.454 mmol) and Cs_2_CO_3 anhyd_ (99 mg, 0.303 mmol). A yellow semi-solid was obtained 24 mg (36% yield).^1^H NMR (500 MHz, MeOD) δ 7.43 (s, 2H), 6.85 (s, 2H), 4.13 – 4.04 (m, 4H), 4.00 (dd, *J* = 13.2, 6.6 Hz, 2H), 3.92 (s, 6H), 3.87 (s, 6H), 3.73 – 3.69 (m, 11H), 3.47 (t, *J* = 6.7 Hz, 4H), 2.97 – 2.87 (m, 4H), 2.50 (dd, *J* = 9.2, 5.4 Hz, 14H), 2.38 (s, 12H), 1.91 (dd, *J* = 19.7, 9.1 Hz, 3H), 1.84 (dd, *J* = 14.4, 7.2 Hz, 6H). A set of rotamers is observed; LCMS Column 1, RT = 0.97 min, MS (ESI): m/z = 445.52 [M + 1]^+^, Purity (UV^254^) = 100%; HRMS (ESI) for C_23_H_37_N_6_O_3_ [M + H]^+^: m/z = calcd, 445.2927; found, 445.2917.

### 2-Chloro-6,7-dimethoxy-4-(pyrrolidin-1-yl)quinazoline

The title compound was synthesized according to the general procedure **A**. 2,4-Dichloro-6,7-dimethoxyquinazoline (100 mg, 0.386 mmol), tetrahydrofuran (5 mL), *N,N*-diisopropylethylamine (0.202 mL, 1.158 mmol), pyrrolidine 99% (0.064 mL, 0.772 mmol). MS (ESI): m/z = 294.33 [M + H]^+^.

### *N*^1^-(6,7-Dimethoxy-4-(pyrrolidin-1-yl)quinazolin-2-yl)-*N*^3^,*N*^3^-dimethylpropane-1,3-diamine (OICR18632)

The title compound was synthesized according to the general procedure **B**. Palladium(II) acetate (2.64 mg, 0.012 mmol), 2,2’-*bis*(diphenylphosphino)-1,1’-binaphthalene (14.63 mg, 0.023 mmol), 1,4-dioxane (783 µL), 2-chloro-6,7-dimethoxy-4-(pyrrolidin-1-yl)quinazoline (69 mg, 0.235 mmol), 3-(dimethylamino)-1-propylamine (89 µL, 0.705 mmol) and Cs_2_CO_3 anhyd_ (153 mg, 0.470 mmol). An off-white powder was obtained 54 mg (64% yield); ^1^H NMR (500 MHz, MeOD) δ 7.48 (s, 1H), 6.82 (s, 1H), 3.92 (s, 2H), 3.91 (s, 3H), 3.89 (s, 2H), 3.86 (s, 3H), 3.44 (t, *J* = 6.8 Hz, 2H), 2.49 – 2.45 (m, 2H), 2.29 (s, 6H), 2.03 (t, *J* = 6.5 Hz, 4H), 1.83 (dd, *J* = 14.8, 7.3 Hz, 2H); LCMS Column 1, RT = 1.08 min, MS (ESI): m/z = 360.44 [M + 1]^+^, Purity (UV^254^) = 100%; HRMS (ESI) for C_19_H_30_N_5_O_2_ [M + H]^+^: m/z = calcd, 360.2400; found, 360.2397.

### 2-Chloro-6,7-dimethoxy-4-(piperidin-1-yl)quinazoline

The title compound was synthesized according to the general procedure **A**. 2,4-dichloro-6,7-dimethoxyquinazoline (107 mg, 0.413 mmol), tetrahydrofuran (4.5 mL), *N,N*-diisopropylethylamine (216 µL, 1.239 mmol) and piperidine (82 µl, 0.826 mmol). ^1^H NMR (500 MHz, MEOD) δ 7.18 (s, 1H), 7.06 (s, 1H), 3.98 (s, 3H), 3.94 (s, 3H), 3.75 (s, 4H), 1.80 (s, 6H). MS (ESI): m/z = 308.30 [M + H]^+^.

### *N*^1^-(6,7-Dimethoxy-4-(piperidin-1-yl)quinazolin-2-yl)-*N*^3^,*N*^3^-dimethylpropane-1,3-diamine (OICR19016)

The title compound was synthesized according to the general procedure **B**. Palladium(II) acetate (1.787 mg, 7.96 µmol), 2,2’-*bis*(diphenylphosphino)-1,1’-binaphthalene (9.91 mg, 0.016 mmol), 1,4-dioxane (531 µL), 2-chloro-6,7-dimethoxy-4-(piperidin-1-yl)quinazoline (531 µL, 0.159 mmol), 3-(dimethylamino)-1-propylamine (60.1 µL, 0.478 mmol) and Cs_2_CO_3 anhyd_ (104 mg, 0.318 mmol). ^1^H NMR (500 MHz, MEOD) δ 7.06 (s, 1H), 6.86 (s, 1H), 3.92 (s, 3H), 3.87 (s, 3H), 3.57 (br s, 4H), 3.51 – 3.34 (m, 2H), 2.51 – 2.41 (m, 2H), 2.28 (s, 6H), 1.90 – 1.80 (m, 3H), 1.80 – 1.73 (m, 6H); LCMS Column 1, RT = 1.28 min, MS (ESI): m/z = 374.68 [M + 1]^+^, Purity (UV^254^) = 100%; HRMS (ESI) for C_20_H_32_N_5_O_2_ [M + H]^+^: m/z = calcd, 374.2556; found, 374.2549.

### 4-(2-Chloro-6,7-dimethoxyquinazolin-4-yl)morpholine

The title compound was synthesized according to the general procedure **A**. 2,4-dichloro-6,7-dimethoxyquinazoline (100 mg, 0.386 mmol), tetrahydrofuran (5 mL) were added *N,N*-diisopropylethylamine (0.202 mL, 1.158 mmol) and morpholine (0.067 ml, 0.772 mmol). MS (ESI): m/z = 310.27 [M + H]^+^.

### *N*^1^-(6,7-dimethoxy-4-morpholinoquinazolin-2-yl)-*N*^3^,*N*^3^-dimethylpropane-1,3-diamine (OICR18716)

The title compound was synthesized according to the general procedure **B**. Palladium(II) acetate (2.68 mg, 0.012 mmol), 2,2’-*bis*(diphenylphosphino)-1,1’-binaphthalene (14.88 mg, 0.024 mmol), 1,4-dioxane (796 µL), 4-(2-chloro-6,7-dimethoxyquinazolin-4-yl)morpholine (74 mg, 0.239 mmol), 3-(dimethylamino)-1-propylamine (90 µL, 0.717 mmol) and Cs_2_CO_3 anhyd_ (156 mg, 0.478 mmol). A yellow semi-solid was obtained 24 mg (27% yield). ^1^H NMR (500 MHz, MeOD) δ 7.03 (s, 1H), 6.87 (s, 1H), 3.90 (s, 3H), 3.85 (s, 3H), 3.59 – 3.56 (m, 4H), 3.43 (t, *J* = 6.8 Hz, 2H), 2.47 – 2.43 (m, 2H), 2.30 (s, 4H), 2.27 (s, 6H), 1.85 – 1.78 (m, 2H); LCMS Column 1, RT = 1.05 min, MS (ESI): m/z = 376.46 [M + 1]^+^, Purity (UV^254^) = 100%; HRMS (ESI) for C_19_H_30_N_5_O_3_ [M + H]^+^: m/z = calcd, 376.2349; found, 376.2344.

### *tert*-Butyl 4-(2-chloro-6,7-dimethoxyquinazolin-4-yl)piperazine-1-carboxylate

The title compound was synthesized according to the general procedure **A**. 2,4-Dichloro-6,7-dimethoxyquinazoline (150 mg, 0.579 mmol), tetrahydrofuran (5 mL) were added *N,N*-diisopropylethylamine (0.302 mL, 1.737 mmol) and 1-boc-piperazine (216 mg, 1.158 mmol). MS (ESI): m/z = 409.26 [M + H]^+^.

### *N*^1^-(6,7-Dimethoxy-4-(piperazin-1-yl)quinazolin-2-yl)-*N*^3^,*N*^3^-dimethylpropane-1,3-diamine (OICR19093)

The title compound was synthesized according to the general procedure **B**. Palladium(II) acetate (2.75 mg, 0.012 mmol), 2,2’-*bis*(diphenylphosphino)-1,1’-binaphthalene (15.23 mg, 0.024 mmol), 1,4-dioxane (815 µL), *tert*-butyl 4-(2-chloro-6,7-dimethoxyquinazolin-4-yl)piperazine-1-carboxylate (100 mg, 0.245 mmol), 3-(dimethylamino)-1-propylamine (92 µL, 0.734 mmol) and Cs_2_CO_3 anhyd_ (159 mg, 0.489 mmol). An orange powder was obtained 19 mg (21% yield). ^1^H NMR (500 MHz, MeOD) δ 7.06 (s, 1H), 6.89 (s, 1H), 3.93 (s, 3H), 3.88 (s, 3H), 3.60 (s, 4H), 3.46 (t, *J* = 6.7 Hz, 2H), 3.08 – 3.01 (m, 4H), 2.61 – 2.53 (m, 2H), 2.37 (s, 6H), 1.90 – 1.83 (m, 2H); LCMS Column 1, RT = 0.75 min, MS (ESI): m/z = 375.33 [M + 1]^+^, Purity (UV^254^) = 100%; HRMS (ESI) for C_19_H_31_N_6_O_2_ [M + H]^+^: m/z = calcd, 375.2508; found, 375.2502.

### 2-Chloro-6,7-dimethoxy-4-(4-methylpiperazin-1-yl)quinazoline

The title compound was synthesized according to the general procedure **A**. 2,4-Dichloro-6,7-dimethoxyquinazoline (100 mg, 0.386 mmol), tetrahydrofuran (5 mL), *N,N*-diisopropylethylamine (0.202 mL, 1.158 mmol) and 1-methylpiperazine (0.086 mL, 0.772 mmol). MS (ESI): m/z = 323.36 [M + H]^+^.

### *N*^1^-(6,7-Dimethoxy-4-(4-methylpiperazin-1-yl)quinazolin-2-yl)-*N*^3^,*N*^3^-dimethylpropane-1,3-diamine (OICR18631)

The title compound was synthesized according to the general procedure **B**. Palladium(II) acetate (2.57 mg, 0.011 mmol), 2,2’-*bis*(diphenylphosphino)-1,1’-binaphthalene (14.28 mg, 0.023 mmol), 1,4-dioxane (764 µL), 2-chloro-6,7-dimethoxy-4-(4-methylpiperazin-1-yl)quinazoline (74 mg, 0.229 mmol), 3-(dimethylamino)-1-propylamine (87 µL, 0.688 mmol) and Cs_2_CO_3 anhyd_ (149 mg, 0.458 mmol). An orange semi-solid was obtained 36 mg (40% yield). ^1^H NMR (500 MHz, MeOD) δ 7.05 (s, 1H), 6.89 (s, 1H), 3.92 (s, 3H), 3.88 (s, 3H), 3.64 (s, 4H), 3.45 (t, *J* = 6.8 Hz, 2H), 2.68 – 2.65 (m, 4H), 2.50 – 2.46 (m, 2H), 2.38 (s, 3H), 2.29 (s, 6H), 1.87 – 1.81 (m, 2H); LCMS Column 1, RT = 0.80 min, MS (ESI): m/z = 389.55 [M + 1]^+^, Purity (UV^254^) = 100%; HRMS (ESI) for C_20_H_33_N_6_O_2_ [M + H]^+^: m/z = calcd, 389.2665; found, 389.2658.

### 2-Chloro-6,7-dimethoxy-N,N-dimethylquinazolin-4-amine

The title compound was synthesized according to the general procedure **A**. 2,4-Dichloro-6,7-dimethoxyquinazoline (100 mg, 0.386 mmol), tetrahydrofuran (5 mL), *N,N*-diisopropylethylamine (0.202 mL, 1.158 mmol) and dimethylamine, 2.0 M solution in methanol (0.386 mL, 0.772 mmol). MS (ESI): m/z = 268.08 [M + H]^+^.

### *N*^2^-(3-(Dimethylamino)propyl)-6,7-dimethoxy-*N*^4^,*N*^4^-dimethylquinazoline-2,4-diamine (OICR18718)

The title compound was synthesized according to the general procedure **B**. Palladium(II) acetate (2.097 mg, 9.34 µmol), 2,2’-*bis*(diphenylphosphino)-1,1’-binaphthalene (11.63 mg, 0.019 mmol), 1,4-dioxane (623 µL), 2-chloro-6,7-dimethoxy-*N,N*-dimethylquinazolin-4-amine (50 mg, 0.187 mmol), 3-(dimethylamino)-1-propylamine (70.5 µL, 0.560 mmol) and Cs_2_CO_3 anhyd_ (122 mg, 0.374 mmol). A yellow semi-solid was obtained 21 mg (34% yield).^1^H NMR (500 MHz, MeOD) δ 7.28 (s, 1H), 6.86 (s, 1H), 3.92 (s, 3H), 3.87 (s, 3H), 3.46 (t, *J* = 6.8 Hz, 2H), 3.28 (s, 6H), 2.55 – 2.50 (m, 2H), 2.33 (s, 6H), 1.86 (dt, *J* = 14.4, 7.0 Hz, 2H); LCMS Column 1, RT = 1.11 min, MS (ESI): m/z = 334.41 [M + 1]^+^, Purity (UV^254^) = 100%; HRMS (ESI) for C_17_H_28_N_5_O_2_ [M + H]^+^: m/z = calcd, 334.2243; found, 334.2238.

### *N*^2^-(3-(Dimethylamino)propyl)-6,7-dimethoxyquinazoline-2,4-diamine (OICR18745)

The title compound was synthesized according to the general procedure **B**. Palladium(II) acetate (2.342 mg, 10.43 µmol), 2,2’-*bis*(diphenylphosphino)-1,1’-binaphthalene (12.99 mg, 0.021 mmol), 1,4-dioxane (695 µL), 4-amino-2-chloro-6,7-dimethoxyquinazoline (50 mg, 0.209 mmol), 3-(dimethylamino)-1-propylamine (79 µL, 0.626 mmol) and Cs_2_CO_3 anhyd_ (136 mg, 0.417 mmol). A white semi-solid was obtained 9.5 mg (15% yield). Preparative HPLC purification. ^1^H NMR (500 MHz, MeOD) δ 7.33 (s, 1H), 6.83 (s, 1H), 3.92 (s, 3H), 3.88 (s, 3H), 3.43 (t, *J* = 6.8 Hz, 2H), 2.52 – 2.47 (m, 2H), 2.31 (s, 6H), 1.86 – 1.80 (m, 2H); LCMS Column 1, RT = 0.85 min, MS (ESI): m/z = 306.44 [M + 1]^+^, Purity (UV^254^) = 100%; HRMS (ESI) for C_15_H_24_N_5_O_2_ [M + H]^+^: m/z = calcd, 306.1930; found, 306.1924.

### *tert*-Butyl (1-(2-chloro-6,7-dimethoxyquinazolin-4-yl)azetidin-3-yl)carbamate

The title compound was synthesized according to the general procedure **A**. 2,4-Dichloro-6,7-dimethoxyquinazoline (150 mg, 0.579 mmol), tetrahydrofuran (5 ml), *N,N*-diisopropylethylamine (0.302 mL, 1.737 mmol) and 3-(BOC-amino)-azetidine (199 mg, 1.158 mmol). MS (ESI): m/z = 395.27 [M + H]^+^.

### *N*^1^-(4-(3-aminoazetidin-1-yl)-6,7-dimethoxyquinazolin-2-yl)-*N*^3^,*N*^3^-dimethylpropane-1,3-diamine (OICR19102)

The title compound was synthesized according to the general procedure **B**. Palladium(II) acetate (2.84 mg, 0.013 mmol), 2,2’-bis(diphenylphosphino)-1,1’-binaphthalene (15.77 mg, 0.025 mmol), 1,4-dioxane (844 µL), *tert*-butyl (1-(2-chloro-6,7-dimethoxyquinazolin-4-yl)azetidin-3-yl)carbamate (100 mg, 0.253 mmol), 3-(dimethylamino)-1-propylamine (96 µL, 0.760 mmol) and Cs_2_CO_3 anhyd_ (165 mg, 0.507 mmol). An off-white powder was obtained 24 mg (24% yield). ^1^H NMR (500 MHz, MeOD) δ 7.12 (s, 1H), 6.84 (s, 1H), 4.71 (t, *J* = 8.3 Hz, 2H), 4.16 (dd, *J* = 9.1, 5.2 Hz, 2H), 3.98 – 3.93 (m, 1H), 3.91 (s, 3H), 3.86 (s, 3H), 3.44 (t, *J* = 6.7 Hz, 2H), 2.53 – 2.49 (m, 2H), 2.33 (s, 6H), 1.87 – 1.81 (m, 2H); LCMS Column 1, RT = 0.75 min, MS (ESI): m/z = 361.34 [M + 1]^+^, Purity (UV^254^) = 96%; HRMS (ESI) for C_18_H_29_N_6_O_2_ [M + H]^+^: m/z = calcd, 361.2352; found, 361.2350.

### 1-(2-Chloro-6,7-dimethoxyquinazolin-4-yl)-N,N-dimethylazetidin-3-amine

The title compound was synthesized according to the general procedure **A**. 2,4-Dichloro-6,7-dimethoxyquinazoline (100 mg, 0.386 mmol), tetrahydrofuran (5 mL) were added *N,N*-diisopropylethylamine (0.202 mL, 1.158 mmol) and 3-(dimethylamino)azetidine dihydrochloride (134 mg, 0.772 mmol). MS (ESI): m/z = 323.36 [M + H]^+^.

### *N*^1^-(4-(3-(Dimethylamino)azetidin-1-yl)-6,7-dimethoxyquinazolin-2-yl)-*N*^3^,*N*^3^-dimethylpropane-1,3-diamine (OICR18629)

The title compound was synthesized according to the general procedure **B**. Palladium(II) acetate (2.504 mg, 0.011 mmol), 2,2’-*bis*(diphenylphosphino)-1,1’-binaphthalene (13.89 mg, 0.022 mmol), 1,4-dioxane (744 µL), 1-(2-chloro-6,7-dimethoxyquinazolin-4-yl)-*N,N*-dimethylazetidin-3-amine (72 mg, 0.223 mmol), 3-(dimethylamino)-1-propylamine (84 µL, 0.669 mmol) and Cs_2_CO_3 anhyd_ (145 mg, 0.446 mmol). An orange semi-solid was obtained 51 mg (59% yield). ^1^H NMR (500 MHz, MeOD) δ 7.10 (s, 1H), 6.84 (s, 1H), 4.55 (t, *J* = 8.1 Hz, 2H), 4.29 (dd, *J* = 8.9, 5.1 Hz, 2H), 3.91 (s, 3H), 3.87 (d, *J* = 2.7 Hz, 3H), 3.44 (t, *J* = 6.8 Hz, 2H), 2.51 – 2.47 (m, 2H), 2.31 (s, 6H), 2.27 (s, 6H), 1.86 – 1.80 (m, 2H); LCMS Column 1, RT = 0.81 min, MS (ESI): m/z = 389.55 [M + 1]^+^, Purity (UV^254^) = 100%; HRMS (ESI) for C_20_H_33_N_6_O_2_ [M + H]^+^: m/z = calcd, 389.2665; found, 389.2656.

### *tert*-butyl (1-(2-chloro-6,7-dimethoxyquinazolin-4-yl)pyrrolidin-3-yl)carbamate

The title compound was synthesized according to the general procedure **A**. 2,4-Dichloro-6,7-dimethoxyquinazoline (150 mg, 0.579 mmol), tetrahydrofuran (5 mL) were added *N,N*-diisopropylethylamine (0.302 mL, 1.737 mmol) and 3-(*tert*-butoxycarbonylamino)pyrrolidine (216 mg, 1.158 mmol). MS (ESI): m/z = 409.26 [M + H]^+^.

### *N*^1^-(4-(3-aminopyrrolidin-1-yl)-6,7-dimethoxyquinazolin-2-yl)-*N*^3^,*N*^3^-dimethylpropane-1,3-diamine (OICR19099)

The title compound was synthesized according to the general procedure **B**. Palladium(II) acetate (2.75 mg, 0.012 mmol), 2,2’-*bis*(diphenylphosphino)-1,1’-binaphthalene (15.23 mg, 0.024 mmol), 1,4-dioxane (815 µL), *tert*-butyl (1-(2-chloro-6,7-dimethoxyquinazolin-4-yl)pyrrolidin-3-yl)carbamate (100 mg, 0.245 mmol), 3-(dimethylamino)-1-propylamine (92 µL, 0.734 mmol) and Cs_2_CO_3 anhyd_ (159 mg, 0.489 mmol). A yellow powder was obtained 22 mg (23% yield).^1^H NMR (500 MHz, MeOD) δ 7.49 (s, 1H), 6.84 (s, 1H), 4.12 (dt, *J* = 10.7, 6.3 Hz, 2H), 3.92 (s, 3H), 3.88 (s, 3H), 3.72 – 3.63 (m, 2H), 3.46 (t, *J* = 6.7 Hz, 2H), 2.57 – 2.50 (m, 2H), 2.45 (dd, *J* = 13.3, 6.1 Hz, 1H), 2.34 (s, 6H), 2.22 (dd, *J* = 12.7, 6.0 Hz, 1H), 1.91 (dd, *J* = 12.6, 6.2 Hz, 1H), 1.89 – 1.83 (m, 2H); LCMS Column 1, RT = 0.76 min, MS (ESI): m/z = 375.41 [M + 1]^+^, Purity (UV^254^) = 96%; HRMS (ESI) for C_19_H_31_N_6_O_2_ [M + H]^+^: m/z = calcd, 375.2508; found, 375.2507.

### (R)-1-(2-Chloro-6,7-dimethoxyquinazolin-4-yl)-*N*,*N*-dimethylpyrrolidin-3-amine

The title compound was synthesized according to the general procedure **A**. 2,4-Dichloro-6,7-dimethoxyquinazoline (100 mg, 0.386 mmol), tetrahydrofuran (5 mL), *N,N*-diisopropylethylamine (0.202 mL, 1.158 mmol) and (3R)-(+)-3-(dimethylamino)pyrrolidine (0.098 mL, 0.772 mmol). MS (ESI): m/z = 337.42 [M + H]^+^.

### (R)-*N*^1^-(4-(3-(dimethylamino)pyrrolidin-1-yl)-6,7-dimethoxyquinazolin-2-yl)-*N*^3^,*N*^3^-dimethylpropane-1,3-diamine (OICR18627)

The title compound was synthesized according to the general procedure **B**. Palladium(II) acetate (2.466 mg, 10.99 µmol), 2,2’-*bis*(diphenylphosphino)-1,1’-binaphthalene (13.68 mg, 0.022 mmol), 1,4-dioxane (732 µL), (R)-1-(2-chloro-6,7-dimethoxyquinazolin-4-yl)-*N,N*-dimethylpyrrolidin-3-amine (74 mg, 0.220 mmol), 3-(dimethylamino)-1-propylamine (83 µL, 0.659 mmol) and Cs_2_CO_3 anhyd_ (143 mg, 0.439 mmol). An orange semi-solid was obtained 24 mg (27% yield). ^1^H NMR (500 MHz, MeOD) δ 7.43 (s, 1H), 6.84 (s, 1H), 4.11 – 4.04 (m, 2H), 3.92 (s, 3H), 3.87 (s, 3H), 3.75 (t, *J* = 9.4 Hz, 1H), 3.45 (t, *J* = 6.5 Hz, 2H), 2.90 (d, *J* = 7.1 Hz, 2H), 2.51 – 2.48 (m, 2H), 2.37 (s, 6H), 2.31 (s, 6H), 1.84 (s, 2H), 1.76 (dd, *J* = 14.9, 7.4 Hz, 2H); LCMS Column 1, RT = 0.80 min, MS (ESI): m/z = 403.54 [M + 1]^+^, Purity (UV^254^) = 100%; HRMS (ESI) for C_21_H_35_N_6_O_2_ [M + H]^+^: m/z = calcd, 403.2821; found, 403.2814.

### (S)-1-(2-Chloro-6,7-dimethoxyquinazolin-4-yl)-*N*,*N*-dimethylpyrrolidin-3-amine

The title compound was synthesized according to the general procedure **A**. 2,4-Dichloro-6,7-dimethoxyquinazoline (100 mg, 0.386 mmol), tetrahydrofuran (5 mL) were added *N,N*-diisopropylethylamine (0.202 mL, 1.158 mmol) and (S)-(-)-3-(dimethylamino)pyrrolidine (0.098 mL, 0.772 mmol). MS (ESI): m/z = 337.35 [M + H]^+^.

### (S)-*N*^1^-(4-(3-(Dimethylamino)pyrrolidin-1-yl)-6,7-dimethoxyquinazolin-2-yl)-*N*^3^,*N*^3^-dimethylpropane-1,3-diamine (OICR18628)

The title compound was synthesized according to the general procedure **B**. Palladium(II) acetate (2.333 mg, 10.39 µmol), 2,2’-*bis*(diphenylphosphino)-1,1’-binaphthalene (12.94 mg, 0.021 mmol), 1,4-dioxane (693 µL), (S)-1-(2-chloro-6,7-dimethoxyquinazolin-4-yl)-*N,N*-dimethylpyrrolidin-3-amine (70 mg, 0.208 mmol), 3-(dimethylamino)-1-propylamine (78 µL, 0.623 mmol) and Cs_2_CO_3 anhyd_ (135 mg, 0.416 mmol). An orange semi-solid was obtained 45 mg (54% yield). ^1^H NMR (500 MHz, MeOD) δ 7.43 (s, 1H), 6.84 (s, 1H), 4.12 – 4.05 (m, 2H), 3.92 (s, 3H), 3.87 (s, 3H), 3.75 (t, *J* = 9.3 Hz, 1H), 3.45 (t, *J* = 5.3 Hz, 2H), 2.91 (s, 2H), 2.52 – 2.49 (m, 2H), 2.37 (s, 6H), 2.32 (s, 6H), 1.87 – 1.83 (m, 2H), 1.82 – 1.71 (m, 2H); LCMS Column 1, RT = 0.81 min, MS (ESI): m/z = 403.47 [M + 1]^+^, Purity (UV^254^) = 100%; HRMS (ESI) for C_21_H_35_N_6_O_2_ [M + H]^+^: m/z = calcd, 403.2821; found, 403.2812.

### 1-(2-Chloro-6,7-dimethoxyquinazolin-4-yl)-N,N-dimethylpiperidin-4-amine

The title compound was synthesized according to the general procedure **A**. 2,4-Dichloro-6,7-dimethoxyquinazoline (100 mg, 0.386 mmol), tetrahydrofuran (5 mL), *N,N*-diisopropylethylamine (0.202 mL, 1.158 mmol) and 4-(dimethylamino)piperidine (0.109 mL, 0.772 mmol). MS (ESI): m/z = 351.41 [M + H]^+^.

### *N*^1^-(4-(4-(dimethylamino)piperidin-1-yl)-6,7-dimethoxyquinazolin-2-yl)-*N*^3^,*N*^3^-dimethylpropane-1,3-diamine (OICR18630)

The title compound was synthesized according to the general procedure **B**. Palladium(II) acetate (2.336 mg, 10.40 µmol), 2,2’-bis(diphenylphosphino)-1,1’-binaphthalene (12.96 mg, 0.021 mmol), 1,4-dioxane (694 µL), 1-(2-chloro-6,7-dimethoxyquinazolin-4-yl)-*N,N*-dimethylpiperidin-4-amine (73 mg, 0.208 mmol), 3-(dimethylamino)-1-propylamine (79 µL, 0.624 mmol) and Cs_2_CO_3 anhyd_ (136 mg, 0.416 mmol). A yellow semi-solid was obtained 36 mg (41% yield).^1^H NMR (500 MHz, MeOD) δ 6.70 (d, *J* = 3.7 Hz, 1H), 6.38 (d, *J* = 3.7 Hz, 1H), 3.93 – 3.90 (m, 4H), 3.59 (s, 3H), 3.42 (t, *J* = 6.8 Hz, 2H), 2.56 – 2.53 (m, 4H), 2.48 – 2.44 (m, 2H), 2.34 (s, 3H), 2.27 (s, 6H), 1.85 – 1.79 (m, 2H); LCMS Column 1, RT = 0.83 min, MS (ESI): m/z = 417.54 [M + 1]^+^, Purity (UV^254^) = 100%; HRMS (ESI) for C_22_H_37_N_6_O_2_ [M + H]^+^: m/z = calcd, 417.2978; found, 417.2970.

### *tert*-Butyl (1-(2-chloro-6,7-dimethoxyquinazolin-4-yl)piperidin-4-yl)(methyl)carbamate

The title compound was synthesized according to the general procedure **A**. 2,4-Dichloro-6,7-dimethoxyquinazoline (150 mg, 0.579 mmol), tetrahydrofuran (5 mL) were added *N,N*-diisopropylethylamine (0.302 mL, 1.737 mmol) and *tert*-butyl *N*-methyl-*N*-(piperidin-4-yl)carbamate (248 mg, 1.158 mmol). MS (ESI): m/z = 437.25 [M + H]^+^.

### *N*^1^-(6,7-dimethoxy-4-(4-(methylamino)piperidin-1-yl)quinazolin-2-yl)-*N*^3^,*N*^3^-dimethylpropane-1,3-diamine (OICR19101)

The title compound was synthesized according to the general procedure **B**. Palladium(II) acetate (2.57 mg, 0.011 mmol), 2,2’-*bis*(diphenylphosphino)-1,1’-binaphthalene (14.25 mg, 0.023 mmol), 1,4-dioxane (763 µL), *tert*-butyl (1-(2-chloro-6,7-dimethoxyquinazolin-4-yl)piperidin-4-yl)(methyl)carbamate (100 mg, 0.229 mmol), 3-(dimethylamino)-1-propylamine (86 µL, 0.687 mmol) and Cs_2_CO_3 anhyd_ (149 mg, 0.458 mmol). A yellow powder was obtained 31 mg (33% yield). ^1^H NMR (500 MHz, MeOD) δ 7.05 (s, 1H), 6.87 (s, 1H), 4.20 (d, *J* = 13.0 Hz, 2H), 3.92 (s, 3H), 3.88 (s, 3H), 3.45 (t, *J* = 6.7 Hz, 2H), 3.09 (t, *J* = 12.4 Hz, 2H), 2.77 (t, *J* = 10.7 Hz, 1H), 2.51 (t, *J* = 7.5 Hz, 2H), 2.47 (s, 3H), 2.31 (s, 6H), 2.08 (d, *J* = 11.1 Hz, 2H), 1.88 – 1.81 (m, 2H), 1.60 (q, *J* = 11.2 Hz, 2H); LCMS Column 1, RT = 0.77 min, MS (ESI): m/z = 403.47 [M + 1]^+^, Purity (UV^254^) = 100%; HRMS (ESI) for C_21_H_35_N_6_O_2_ [M + H]^+^: m/z = calcd, 403.2821; found, 403.2817.

### *tert*-Butyl (1-(2-chloro-6,7-dimethoxyquinazolin-4-yl)piperidin-4-yl)carbamate

The title compound was synthesized according to the general procedure **A**. 2,4-Dichloro-6,7-dimethoxyquinazoline (150 mg, 0.579 mmol), tetrahydrofuran (5 mL), *N,N*-diisopropylethylamine (0.302 mL, 1.737 mmol) and 4-(*N*-Boc-amino)piperidine (232 mg, 1.158 mmol). MS (ESI): m/z = 423.33 [M + H]^+^.

### *N*^1^-(4-(4-aminopiperidin-1-yl)-6,7-dimethoxyquinazolin-2-yl)-*N*^3^,*N*^3^-dimethylpropane-1,3-diamine (OICR19100)

The title compound was synthesized according to the general procedure **B**. Palladium(II) acetate (2.65 mg, 0.012 mmol), 2,2’-*bis*(diphenylphosphino)-1,1’-binaphthalene (14.72 mg, 0.024 mmol), 1,4-dioxane (788 µL), *tert*-butyl (1-(2-chloro-6,7-dimethoxyquinazolin-4-yl)piperidin-4-yl)carbamate (100 mg, 0.236 mmol), 3-(dimethylamino)-1-propylamine (89 µL, 0.709 mmol) and Cs_2_CO_3 anhyd_ (154 mg, 0.473 mmol). A yellow powder was obtained 35 mg (38% yield). ^1^H NMR (500 MHz, MeOD) δ 7.05 (s, 1H), 6.87 (s, 1H), 4.18 (d, *J* = 13.1 Hz, 2H), 3.92 (s, 3H), 3.88 (s, 3H), 3.45 (t, *J* = 6.7 Hz, 2H), 3.11 (t, *J* = 13.0 Hz, 2H), 3.05 (dd, *J* = 9.5, 5.6 Hz, 1H), 2.53 – 2.48 (m, 2H), 2.32 (s, 6H), 2.00 (d, *J* = 10.6 Hz, 2H), 1.85 (q, *J* = 14.4, 7.0 Hz, 2H), 1.65 (dd, *J* = 20.8, 11.3 Hz, 2H); LCMS Column 1, RT = 0.76 min, MS (ESI): m/z = 389.40 [M + 1]^+^, Purity (UV^254^) = 100%; HRMS (ESI) for C_20_H_33_N_6_O_2_ [M + H]^+^: m/z = calcd, 389.2665; found, 389.2663.

### *tert*-Butyl 4-((2-chloro-6,7-dimethoxyquinazolin-4-yl)amino)piperidine-1-carboxylate

The title compound was synthesized according to the general procedure **A**. 2,4-Dichloro-6,7-dimethoxyquinazoline (150 mg, 0.579 mmol), tetrahydrofuran (5 mL), *N,N*-diisopropylethylamine (0.302 mL, 1.737 mmol) and 4-amino-1-Boc-piperidine (232 mg, 1.158 mmol). MS (ESI): m/z = 423.40 [M + H]^+^.

### *N*^2^-(3-(Dimethylamino)propyl)-6,7-dimethoxy-*N*^4^-(piperidin-4-yl)quinazoline-2,4-diamine (OICR19098)

The title compound was synthesized according to the general procedure **B**. Palladium(II) acetate (2.65 mg, 0.012 mmol), 2,2’-*bis*(diphenylphosphino)-1,1’-binaphthalene (14.72 mg, 0.024 mmol), 1,4-dioxane (788 µL), *tert*-butyl 4-((2-chloro-6,7-dimethoxyquinazolin-4-yl)amino)piperidine-1-carboxylate (100 mg, 0.236 mmol), 3-(dimethylamino)-1-propylamine (89 µL, 0.709 mmol) and Cs_2_CO_3 anhyd_ (154 mg, 0.473 mmol). A white powder was obtained 15 mg (17% yield).^1^H NMR (500 MHz, MeOD) δ 7.43 (s, 1H), 6.80 (s, 1H), 4.33 (t, *J* = 11.2 Hz, 1H), 3.91 (s, 3H), 3.90 (s, 3H), 3.45 (t, *J* = 6.7 Hz, 2H), 3.18 (d, *J* = 14.0 Hz, 2H), 2.78 (t, *J* = 11.6 Hz, 2H), 2.53 – 2.47 (m, 2H), 2.31 (s, 6H), 2.11 (d, *J* = 11.7 Hz, 2H), 1.84 (q, *J* = 14.1, 6.9 Hz, 2H), 1.65 (qd, *J* = 12.6, 3.7 Hz, 2H); LCMS Column 1, RT = 0.76 min, MS (ESI): m/z = 389.40 [M + 1]^+^, Purity (UV^254^) = 100%; HRMS (ESI) for C_20_H_33_N_6_O_2_ [M + H]^+^: m/z = calcd, 389.2665; found, 389.2665.

## Contribution of authors

S.P. and C.Z.-V. contributed equally to this work. Experimental Design: S.P., C.Z.-V., D.M., R.M., M.M., M.A., D.E., D.P., A.A., T.K., R.A.-A., M.V; Crystallization and Structure Determination: S.P., A.D., M.V; Protein purification: E.G., M.V; Data analysis and Review: S.P., C.Z.-V., D.M., R.M., M.M., M.A., D.E., D.P., A.A., T.K., V.T., L.H., G.P., J.O., A..M., C.H. A., P.J.B., M.S., R.A.-A., M.V; Manuscript Writing: S.P., C.Z.-V., M.S., and M.V.; Project Team Supervision: R.A.-A., M.V. All authors have reviewed and approved the final version of the manuscript.

## Acknowledgement

The Structural Genomics Consortium is a registered charity (no: 1097737) that receives funds from Bayer AG, Boehringer Ingelheim, Bristol Myers Squibb, Genentech, Genome Canada through Ontario Genomics Institute [OGI-196], EU/EFPIA/OICR/McGill/KTH/Diamond Innovative Medicines Initiative 2 Joint Undertaking [EUbOPEN grant 875510], Janssen, Merck KGaA (aka EMD in Canada and US), Pfizer and Takeda. Funding for the OICR is provided by the government of Ontario. RBBP4 PDB entry 7M40 data was collected on APS beamline 19ID.

