## Supplemental figures and Tables for "Discovery of small molecule antagonists of human Retinoblastoma Binding Protein 4 (RBBP4)"

[†]Contributed equally

**Table of contents** **Page**


| Entry | Page |
|---|---|
| **Table 1. Peptides used in peptide displacement assays.** | **3** |
| **Table 2. Optimized buffer conditions for the peptide displacement assays.** | **3** |
| **Table 3. Crystallographic data collection and refinement statistics.** | **4** |
| **Figure 1. Crystal structures of RBBP4 and 7 in complex with interacting peptides.** | **5** |
| **Figure 2. Binding pockets of RBBP4.** | **6** |
| **Figure 3. H3 (1-21) competes with FITC-BCL11A for binding to RBBP4 top pocket.** | **7** |
| **Figure 4. Z′-factor determination.** | **7** |
| **Figure 5. Representation of ligand binding to RBBP4.** | **8** |
| **Supplementary Material and Methods.** | **9** |

**Table S1. Peptides used in peptide displacement assays.** FITC-labelled and unlabeled peptides used for developing FP-based peptide displacement assays are listed.

| Peptide | Tag | Sequence | Function |
|---|---|---|---|
| BCL11A (2-16) | FITC | SRRKQGKPQHLSKRE | Binds to RBBP4 Top pocket |
| H3 (1-21) | FITC | ARTKQTARKSTGGKAPRKQLA | Binds to RBBP4 Top pocket |
| MTA1 (656-686) | FITC | DVFYMATEETRKIRKLLSSSETKRAARRPYK | Binds to RBBP4 Side pocket |
| BCL11A (2-16) | Unlabeled | SRRKQGKPQHLSKRE | Displaces FITC-BCL11A(2-16) and FITC-H3(1-15) |
| H3 (1-21) | Unlabeled | ARTKQTARKSTGGKAPRKQLA | Displaces FITC-BCL11A(2-16) |
| MTA1 (656-686) | Unlabeled | DVFYMATEETRKIRKLLSSSETKRAARRPYK | Displaces FITC-MTA1 (656-686) |

**Table S2. Optimized buffer conditions for the FP-based peptide displacement assays.**

| Interaction | Buffer | Protein (nM) | Peptide (nM) |
|---|---|---|---|
| RBBP4-BCL11A (2-16) | 50 mM Tris pH 7.5,150 mM NaCl, 0.01% TX100 | 300 | 10 |
| RBBP4-H3 (1-21) | 10 mM Tris, pH 8.0, 0.005 % NP-40 | 200 | 5 |
| RBBP4-MTA1 (656-686) | 50 mM Sodium phosphate, pH 7.5, 150 mM KCl, 0.01 % NP-40 | 150 | 10 |

**Table S3. Crystallographic data collection and refinement statistics.**

| **PDB ID: 7M40** | |
|---|---|
| **Data collection** | |
| Space group | P21 |
| Wavelength (Å) | 0.97918 |
| a, b, c (Å) | 75.820 59.857 102.011 |
| α, β, γ (°) | 90.000 93.732 90.000 |
| Resolution (Å) | 50.0-1.88 (1.91-1.88)* |
| Rmerge (%) | 6.5% (73.5%)* |
| I/sigma | 20.9 (1.32)* |
| CC 1/2 | 0.970 (0.869)* |
| Redundancy | 5.4 (4.9)* |
| Completeness (%) | 98.0% (93.8%)* |
| **Refinement** | |
| Resolution (Å) | 46.99–1.88 |
| No. reflections | 72452 |
| $R_{work}$ / $R_{free}$(%) | 18.4/23.6 |
| No. atoms/B-factors [$Å^2$] | |
| Protein | 6063/43.2 |
| Ligand | 45/69.9 |
| Water | 337/43.2 |
| Others | n/a |
| R.m.s. deviations | |
| Bond lengths (Å) | 0.010 |
| Bond angles (°) | 1.380 |

***The values in parentheses refer to statistics in the highest resolution bin.**

A

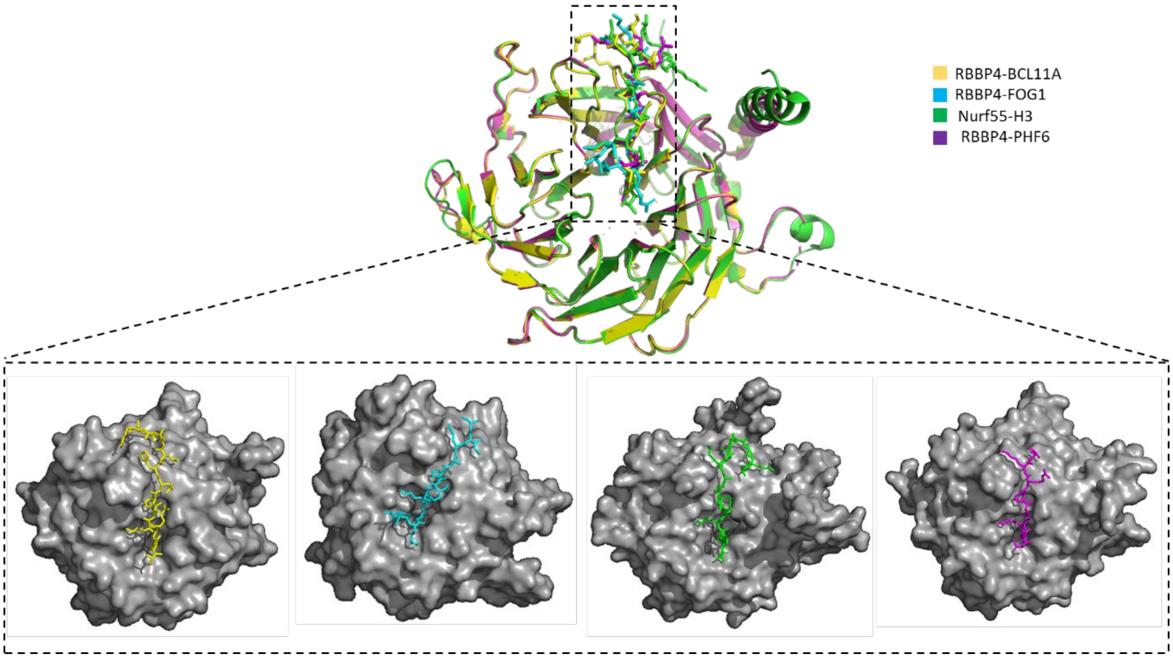


B

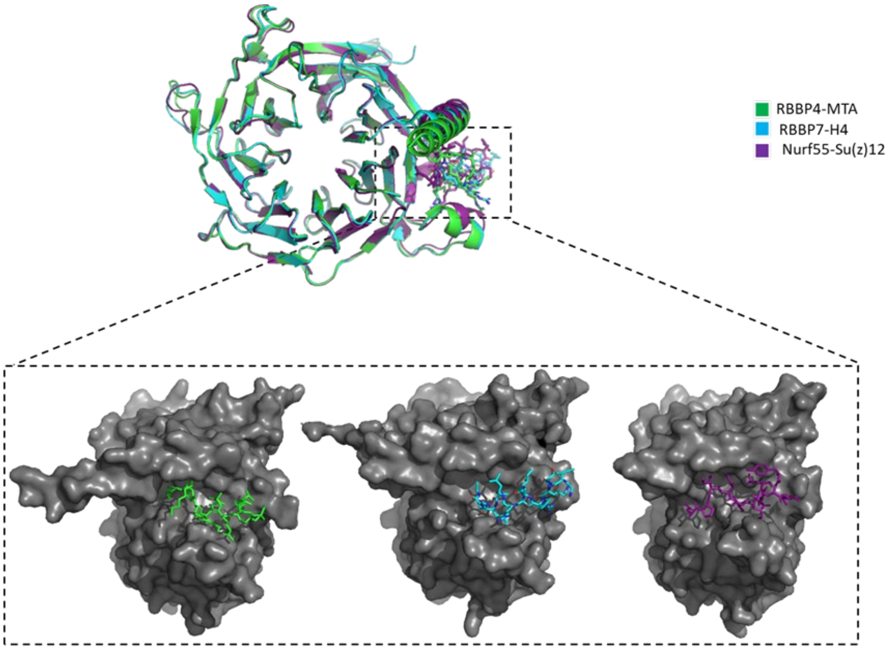


**Figure S1. Crystal structures of RBBP4 and RBBP7 in complex with interacting peptides.** (A) Structural alignment of RBBP4 in complex with BCL11A, FOG-1, H3 and PHF6 (PDB code: 5VTB, 2XU7 and 4R7A, respectively) and Nurf55 in complex with histone H3 peptide (PDB code: 2YBA). (B) Structural alignment of RBBP4 in complex with MTA1 (PDB code: 4PBY), RBBP7 in complex with histone H4 (PDB code: 3CFV), and Nurf55 in complex with Su(z)12 (PDB code: 2YB8). All crystal structures were downloaded from RCSB Protein Data Bank and analyzed using PyMOL.

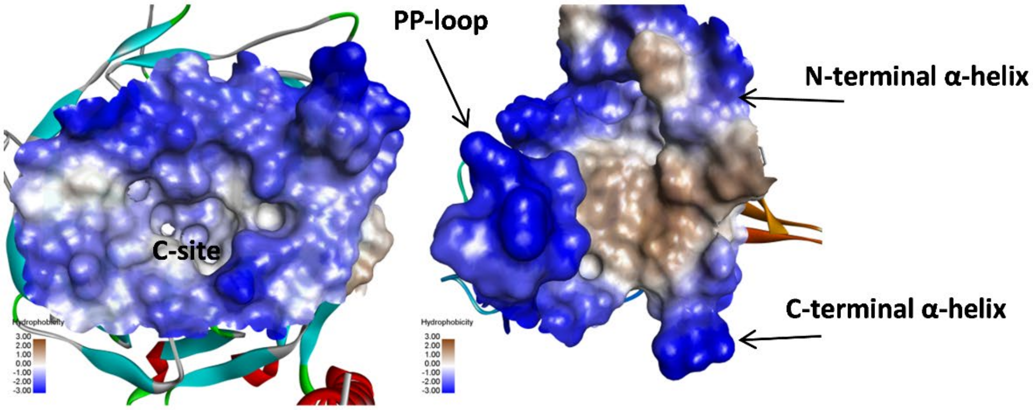


**Figure S2. Binding pockets of RBBP4.** The negative charges (blue) and hydrophobic surface (gray) in the protein-binding pockets of the RBBP4 are presented. C-site on the *left* and side binding pocket on the *right.*

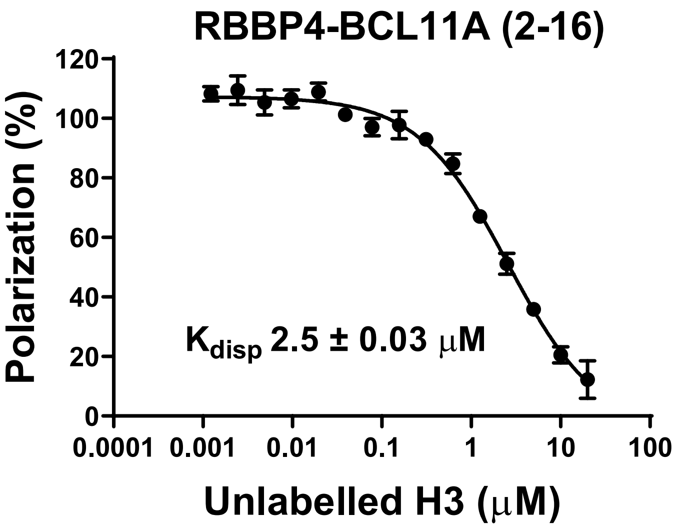


**Figure S3. H3 (1-21) peptide competes with FITC-BCL11A for binding to RBBP4 top pocket.** The histone H3 peptide (1-21) competes with and displaces FITC-BCL11A (2-16) peptide from RBBP4 top pocket. All experiments were performed in triplicate, and data are shown as the mean ± standard deviation.

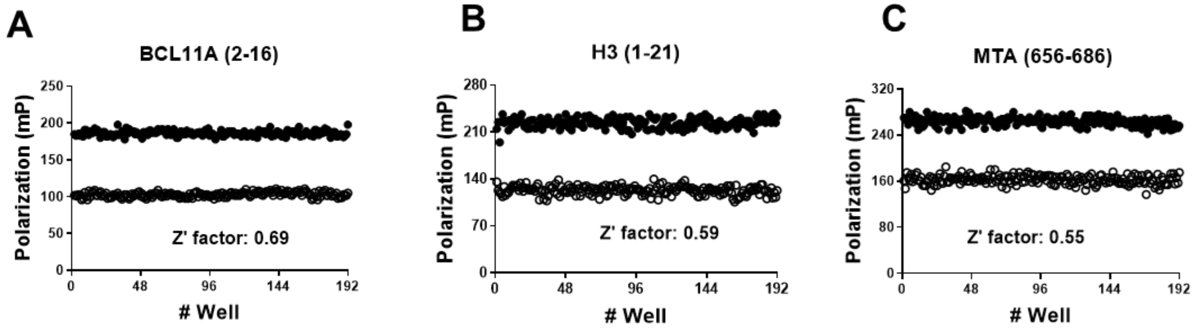


**Figure S4. Z′-factor determination.** Suitability of the peptide displacement assays for screening was determined by calculating the Z′-factor using (A) BCL11A (2-16) (0.69), and (B) H3 (1-21) (0.59) for probing the top pocket, and (C) MTA1 656-686 (0.55) for probing the side pocket of RBBP4. For all cases, 192 samples included only the FITC-labelled peptide (●), and 192 samples included both labelled and competing unlabeled (○) peptides.

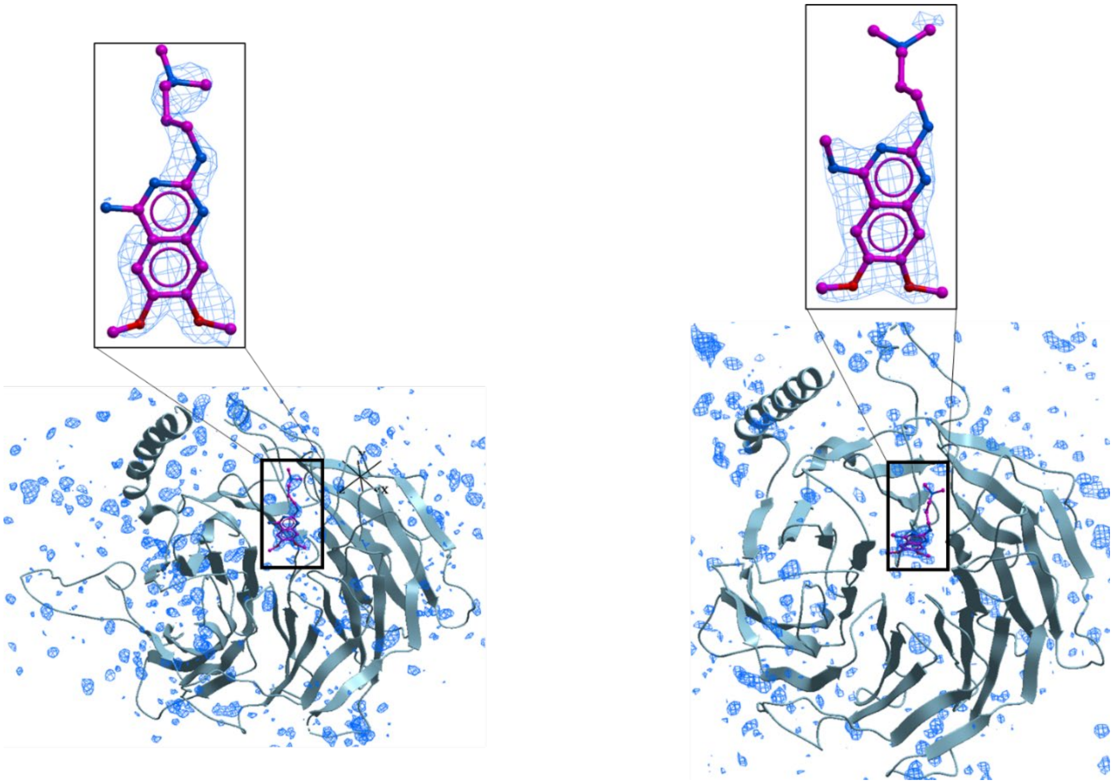

**Figure S5. Representation of ligand binding to RBBP4.** Extended view of the mFo-DFc electron density contoured at level 3.0 (inserts contoured at 2.5) with Molsoft's ICM (blue mesh) with a proposed ligand binding pose (magenta sticks). Left: crystallographic molecule A. Right: crystallographic molecule B. (PDB ID: 7M40).

**Supplementary Materials and Methods.**

**Chemistry and Compound Purity**

All reagents were purchased from commercial vendors and used without further purification. Volatiles were removed under reduced pressure by rotary evaporation or by using the V-10 solvent evaporator system by Biotage®. Very high boiling point (6000 rpm, 0 mbar, 56 ºC), mixed volatile (7000 rpm, 30 mbar, 36 ºC) and volatile (6000 rpm, 30 mbar, 36 ºC) methods were used to evaporate solvents. The yields given refer to chromatographically purified and spectroscopically pure compounds.

*Flash and Preparative Column Chromatography*

Compounds were purified using a Biotage Isolera One system by normal phase chromatography using Biotage® SNAP KP-Sil or Sfär Silica D columns (Part No.: FSKO-1107/FSRD-0445) or by reverse-phase chromatography using Biotage® SNAP KP-C18-HS or Sfär C18 D (Part No.: FSLO-1118/FSUD-040).

If additional purification was required, compounds were purified by solid phase extraction (SPE) using Biotage Isolute Flash SCX-2 cation exchange cartridges (Part No.: 532-0050-C and 456-0200-D). Products were washed with 2 cartridge volumes of MeOH and eluted with a solution of MeOH and $NH_4OH$ (9:1 v/v).

Preparative chromatography was carried out using a Waters 2767 injector with the collector attached to PDA UV/Vis and SQD mass detectors. An XSelect CSH Prep C18 5µm OBD 19 mm x 100 mm (Part No.: 186005421) or Xselect CSH Prep C18 5µm 10 mm x 100 mm (Part No.: 186005415) column was used for purification.
Final compounds were dried using the Labconco™ Benchtop FreeZone™ Freeze-Dry System (4.5 L Model).

*Nuclear Magnetic Resonance (NMR)*

$^{1}H$ and proton-decoupled $^{19}F$ NMRs were recorded on a Bruker Avance-III 500 MHz spectrometer at ambient temperature. Residual protons of $CDCl_3$, DMSO-$d_6$ and $CD_3OD$ solvents were used as internal references. Spectral data are reported as follows: chemical shift (δ in ppm), multiplicity (br = broad, s = singlet, d = doublet, dd = doublet of doublets, m = multiplet), coupling constants (*J* in Hz) and proton integration.

*Compound Purity Determination*

Conducted by UV absorbance at 254 nm during tandem liquid chromatography/mass spectrometry (LCMS) using a Waters Acquity separations module. **All final compounds had a purity of ≥95% as determined using this method`.**

*Low Resolution Mass Spectrometry (LRMS)*
Conducted in positive ion mode using a Waters Acquity SQD mass spectrometer (electrospray ionization source) fitted with a PDA detector. Mobile phase A consisted of 0.1% formic acid in water, while mobile phase B consisted of 0.1% formic acid in acetonitrile. The gradient that was followed is presented in the table below.

| **Time (min)** | **Flow (mL/min)** | **Column 1,2** | | **Column 3** | |
|---|---|---|---|---|---|
| | | **%A** | **%B** | **%A** | **%B** |
| Initial | 0.4 | 90 | 10 | 98 | 2 |
| 1.8 | 0.4 | 5 | 95 | 5 | 95 |
| 2.3 | 0.4 | 5 | 95 | 5 | 95 |
| 2.5 | 0.4 | 90 | 10 | 98 | 2 |
| 3 | 0.4 | 90 | 10 | 98 | 2 |
| 5 | 0 | 90 | 10 | 98 | 2 |

Column 1: Acquity UPLC CSH C18 (2.1 x 50 mm, 130 Å, 1.7 µm. Part No. 186005296) or
Column 2: Acquity UPLC BEH C8 (2.1 x 50 mm, 130 Å, 1.7 µm. Part No. 186002877) or
Column 3: Acquity UPLC HSS T3 (2.1 x 50 mm, 100 Å, 1.8 µm. Part No. 186003538).
All were used with column temperature maintained at 25 °C.

*High Resolution Mass Spectrometry (HRMS)*
Conducted using a Waters Synapt G2-S quadrupole-time-of-flight (QTOF) hybrid mass spectrometer system coupled with an Acquity ultra-performance liquid chromatography (UPLC) system.

*Chromatographic Separations*
Carried out on an Acquity UPLC CSH C18 (2.1 x 50 mm, 130 Å, 1.7 µm. Part No. 186005296) or Acquity UPLC BEH C8 (2.1 x 50 mm, 130 Å, 1.7 µm. Part No. 186002877) or Acquity UPLC HSS T3 (2.1 x 50 mm, 100 Å, 1.8 µm. Part No. 186003538). The mobile phase was 0.1% formic acid in water (solvent A) and 0.1% formic acid in acetonitrile (solvent B). Leucine Enkephalin was used as lock mass. MassLynx 4.1 was used for data analysis.

**Z′-factor calculation:**
The quality and robustness of the screening assays were verified by the standard Z′-factor determination. Z′-factors were calculated using 192 samples with FITC-labeled-peptides (corresponding to 80% of polarization signal in the binding assay) and 192 samples using unlabeled peptides in the optimized assay conditions (Supplementary Fig. 4) in 384-well format using an Agilent Bravo automated liquid-handling robot. Final DMSO concentration was 2%. Z′-factor was calculated as previously described by Zhang et al[1].

1. Zhang, J. H.; Chung, T. D. Y.; Oldenburg, K. R., A simple statistical parameter for use in evaluation and validation of high throughput screening assays. *J Biomol Screen* **1999,** *4* (2), 67-73.

**High Resolution Accurate Mass Analysis Report**

| No. | OICR ID | CF [M+H] | Calculated [M+H] | Observed [M+H] | ΔMass | Error (ppm) |
|---|---|---|---|---|---|---|
| 1 | OICR0017251A01 | C21H35N6O2 | 403.2821 | 403.2819 | -0.0002 | -0.5 |
| 2 | OICR0018633A01 | C20H33N6O2 | 389.2665 | 389.2660 | -0.0005 | -1.3 |
| 3 | OICR0018634A01 | C22H37N6O2 | 417.2978 | 417.2975 | -0.0003 | -0.7 |
| 4 | OICR0018635A01 | C22H37N6O2 | 417.2978 | 417.2975 | -0.0003 | -0.7 |
| 5 | OICR0018636A01 | C22H35N6O2 | 415.2821 | 415.2818 | -0.0003 | -0.7 |
| 6 | OICR0018637A01 | C23H37N6O2 | 429.2978 | 429.2972 | -0.0006 | -1.4 |
| 7 | OICR0019018A01 | C24H39N6O2 | 443.3134 | 443.3125 | -0.0009 | -2.0 |
| 8 | OICR0018721A01 | C23H37N6O3 | 445.2927 | 445.2917 | -0.0010 | -2.2 |
| 9 | OICR0018632A01 | C19H30N5O2 | 360.2400 | 360.2397 | -0.0003 | -0.8 |
| 10 | OICR0019016A01 | C20H32N5O2 | 374.2556 | 374.2549 | -0.0007 | -1.9 |
| 11 | OICR0018716A01 | C19H30N5O3 | 376.2349 | 376.2344 | -0.0005 | -1.3 |
| 12 | OICR0019093A01 | C19H31N6O2 | 375.2508 | 375.2502 | -0.0006 | -1.6 |
| 13 | OICR0018631A01 | C20H33N6O2 | 389.2665 | 389.2658 | -0.0007 | -1.8 |
| 14 | OICR0018718A01 | C17H28N5O2 | 334.2243 | 334.2238 | -0.0005 | -1.5 |
| 15 | OICR0018745A01 | C15H24N5O2 | 306.1930 | 306.1924 | -0.0006 | -2.0 |
| 16 | OICR0019102A01 | C18H29N6O2 | 361.2352 | 361.2350 | -0.0002 | -0.6 |
| 17 | OICR0018629A01 | C20H33N6O2 | 389.2665 | 389.2656 | -0.0009 | -2.3 |
| 18 | OICR0019099A01 | C19H31N6O2 | 375.2508 | 375.2507 | -0.0001 | -0.3 |
| 19 | OICR0018627A01 | C21H35N6O2 | 403.2821 | 403.2814 | -0.0007 | -1.7 |
| 20 | OICR0018628A01 | C21H35N6O2 | 403.2821 | 403.2812 | -0.0009 | -2.2 |
| 21 | OICR0018630A01 | C22H37N6O2 | 417.2978 | 417.2970 | -0.0008 | -1.9 |
| 22 | OICR0019101A01 | C21H35N6O2 | 403.2821 | 403.2817 | -0.0004 | -1.0 |

| | | | | | | |
|---|---|---|---|---|---|---|
| 23 | OICR0019100A01 | C20H33N6O2 | 389.2665 | 389.2663 | -0.0002 | -0.5 |
| 24 | OICR0019098A01 | C20H33N6O2 | 389.2665 | 389.2665 | 0.0000 | 0.0 |

**Experimental conditions**

| | |
|---|---|
| Mass analyzer | Waters Synapt G2-S Q-TOF mass spectrometer |
| Sample probe | |
| Ionization mode | ESI+ |
| Capillary voltage | 1.0 kV |
| Cone voltage | 25 V |
| Source temperature | 150 °C |
| Desolvation temperature | 500 °C |
| Cone gas flow | 150 L $h^{-1}$ |
| Desolvation gas flow | 500 L $h^{-1}$ |
| Scan time | 0.3 s |
| Reference probe | |
| Capillary voltage | 2.5 kV |
| Scan time | 0.3 s |
| Collision energy | 22 V |
| Reference substance (ions) | Leucine-Enkephalin (m/z 221.0926 and 556.2771) |

| | | | |
|---|---|---|---|
| Liquid chromatography | Waters ACQUITY UPLC I-Class system | | |
| Column | Waters ACQUITY UPLC HSS T3 column (2.1 × 100 mm, 1.8 µm at 40 °C) | | |
| Mobile phase A | 0.1% formic acid in water | | |
| Mobile phase B | 0.1% formic acid in acetonitrile | | |
| Gradient | Time/min | A% | Flow (µL/min) |
| | 0.0 | 95 | 400 |
| | 2.0 | 5 | 400 |
| | 2.5 | 5 | 400 |
| | 3.0 | 95 | 400 |
| | 4.0 | 95 | 400 |

### NMR spectra of products

17251

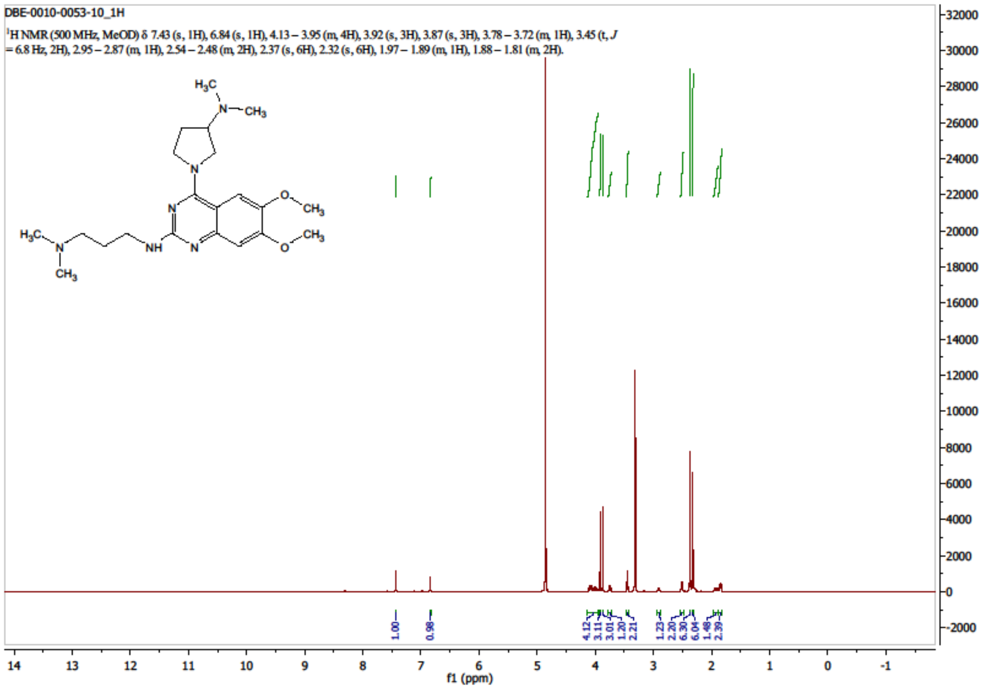


18633

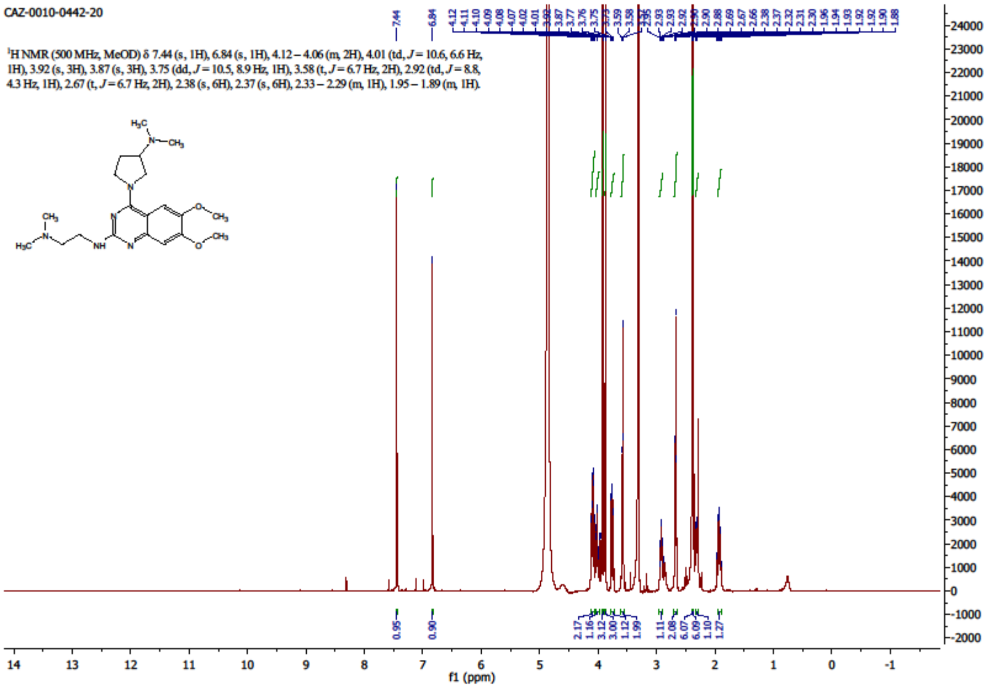

18634

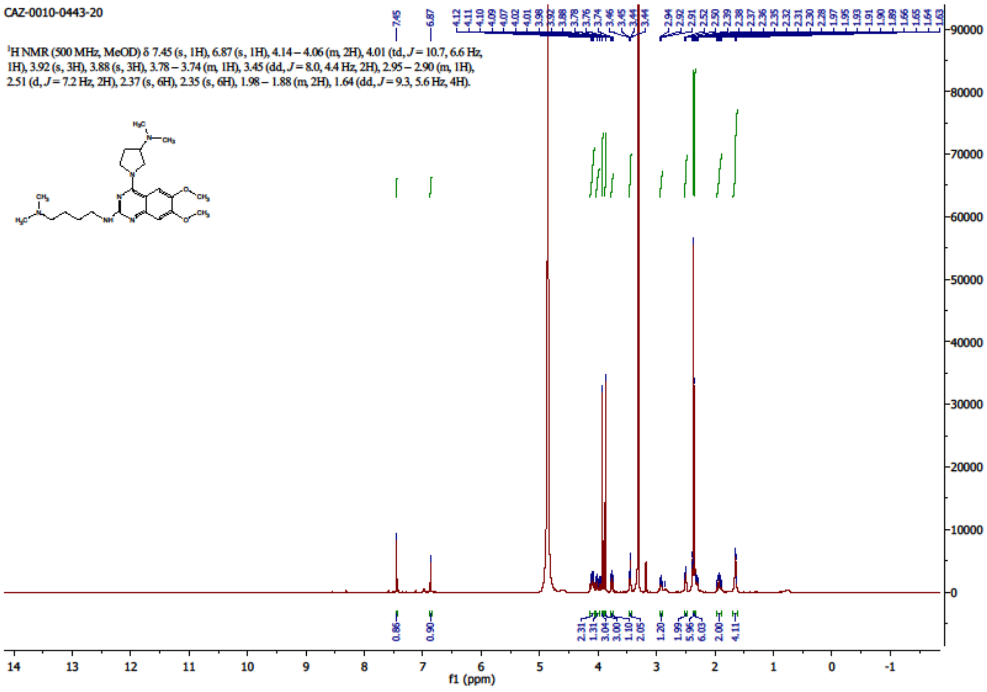
CAZ-0010-0443-20
$^1$H NMR (500 MHz, MeOD) δ 7.45 (s, 1H), 6.87 (s, 1H), 4.14 – 4.06 (m, 2H), 4.01 (td, $J$ = 10.7, 6.6 Hz, 1H), 3.92 (s, 3H), 3.88 (s, 3H), 3.78 – 3.74 (m, 1H), 3.45 (dd, $J$ = 8.0, 4.4 Hz, 2H), 2.95 – 2.90 (m, 1H), 2.51 (d, $J$ = 7.2 Hz, 2H), 2.37 (s, 6H), 2.35 (s, 6H), 1.98 – 1.88 (m, 2H), 1.64 (dd, $J$ = 9.3, 5.6 Hz, 4H).
f1 (ppm)

18635

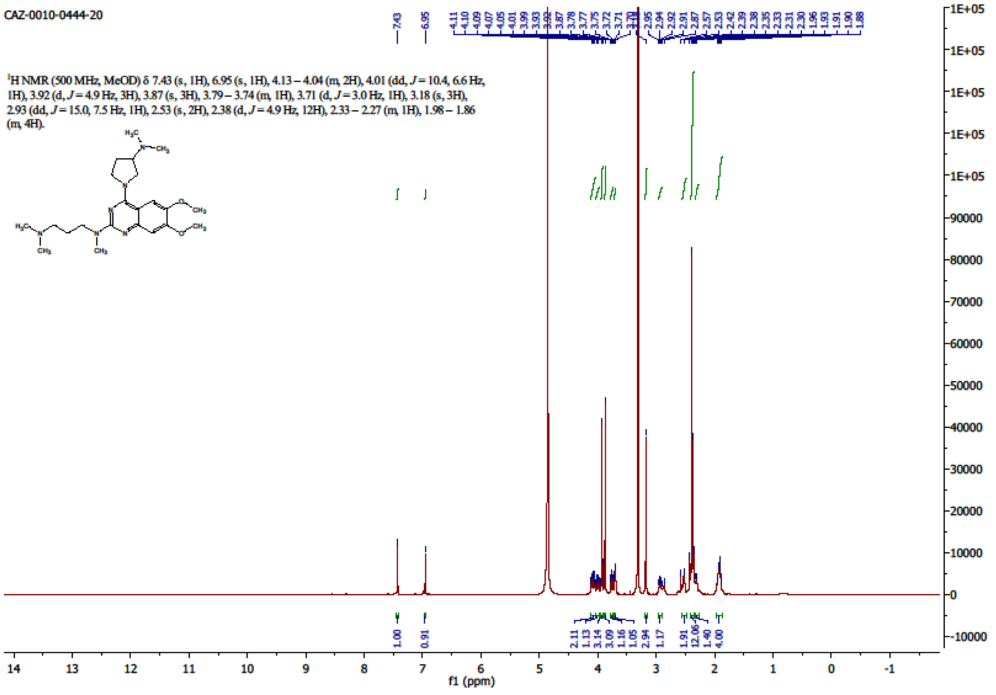
CAZ-0010-0444-20
$^1$H NMR (500 MHz, MeOD) δ 7.43 (s, 1H), 6.95 (s, 1H), 4.13 – 4.04 (m, 2H), 4.01 (dd, $J$ = 10.4, 6.6 Hz, 1H), 3.92 (d, $J$ = 4.9 Hz, 3H), 3.87 (s, 3H), 3.79 – 3.74 (m, 1H), 3.71 (d, $J$ = 3.0 Hz, 1H), 3.18 (s, 3H), 2.93 (dd, $J$ = 15.0, 7.5 Hz, 1H), 2.53 (s, 2H), 2.38 (d, $J$ = 4.9 Hz, 12H), 2.33 – 2.27 (m, 1H), 1.98 – 1.86 (m, 4H).
f1 (ppm)

18636

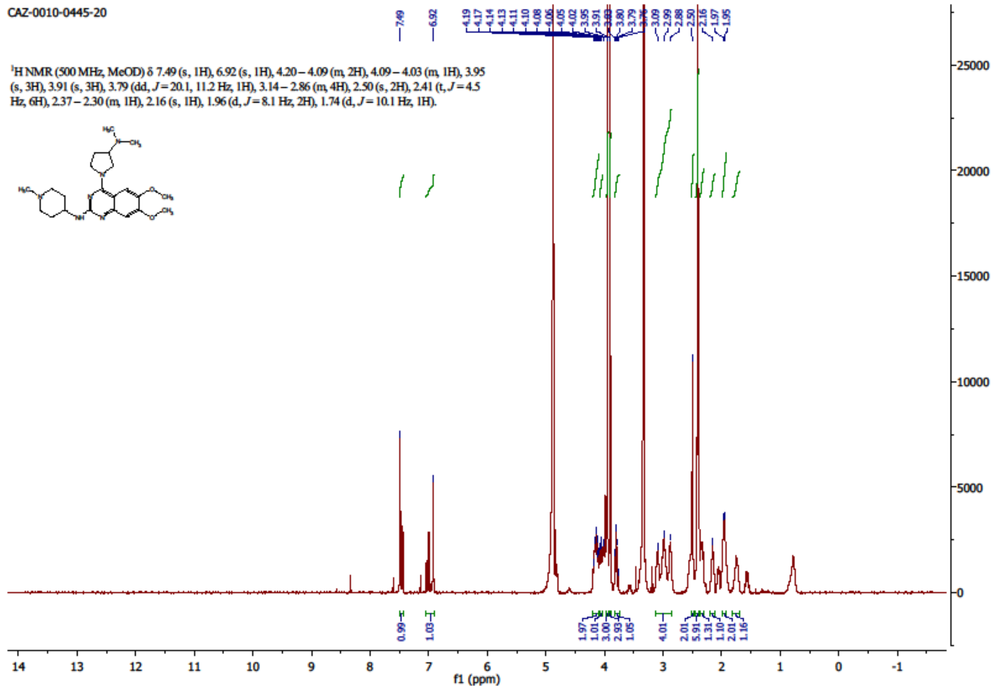
CAZ-0010-0445-20
¹H NMR (500 MHz, MeOD) δ 7.49 (s, 1H), 6.92 (s, 1H), 4.20 – 4.09 (m, 2H), 4.09 – 4.03 (m, 1H), 3.95 (s, 3H), 3.91 (s, 3H), 3.79 (dd, J = 20.1, 11.2 Hz, 1H), 3.14 – 2.86 (m, 4H), 2.50 (s, 2H), 2.41 (t, J = 4.5 Hz, 6H), 2.37 – 2.30 (m, 1H), 2.16 (s, 1H), 1.96 (d, J = 8.1 Hz, 2H), 1.74 (d, J = 10.1 Hz, 1H).
f1 (ppm)

18637

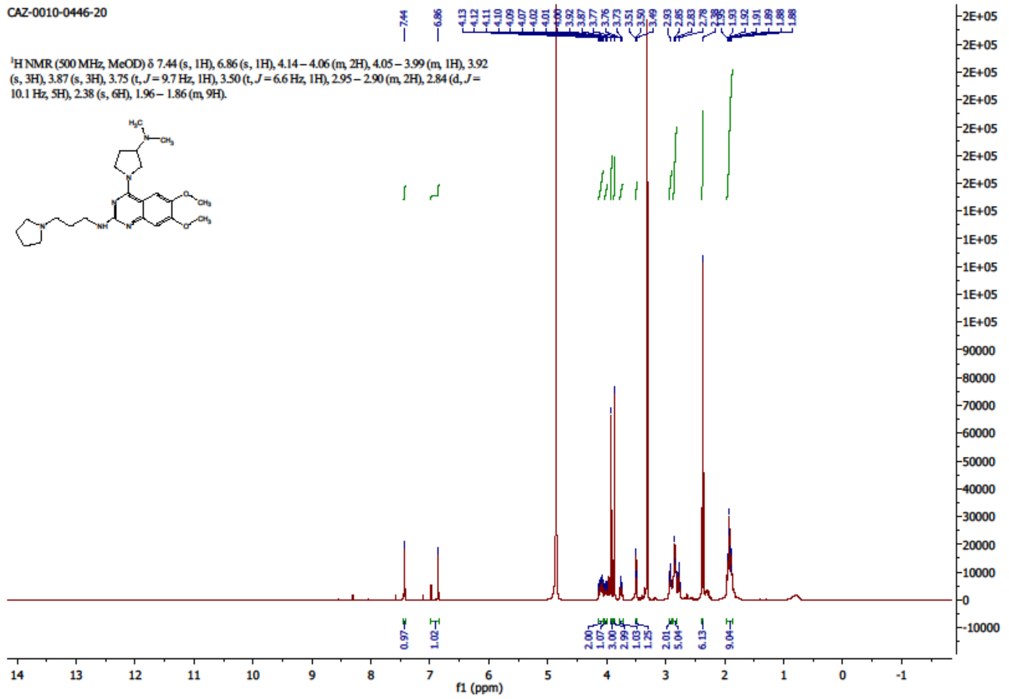
CAZ-0010-0446-20
¹H NMR (500 MHz, MeOD) δ 7.44 (s, 1H), 6.86 (s, 1H), 4.14 – 4.06 (m, 2H), 4.05 – 3.99 (m, 1H), 3.92 (s, 3H), 3.87 (s, 3H), 3.75 (t, J = 9.7 Hz, 1H), 3.50 (t, J = 6.6 Hz, 1H), 2.95 – 2.90 (m, 2H), 2.84 (d, J = 10.1 Hz, 5H), 2.38 (s, 6H), 1.96 – 1.86 (m, 9H).
f1 (ppm)

19018

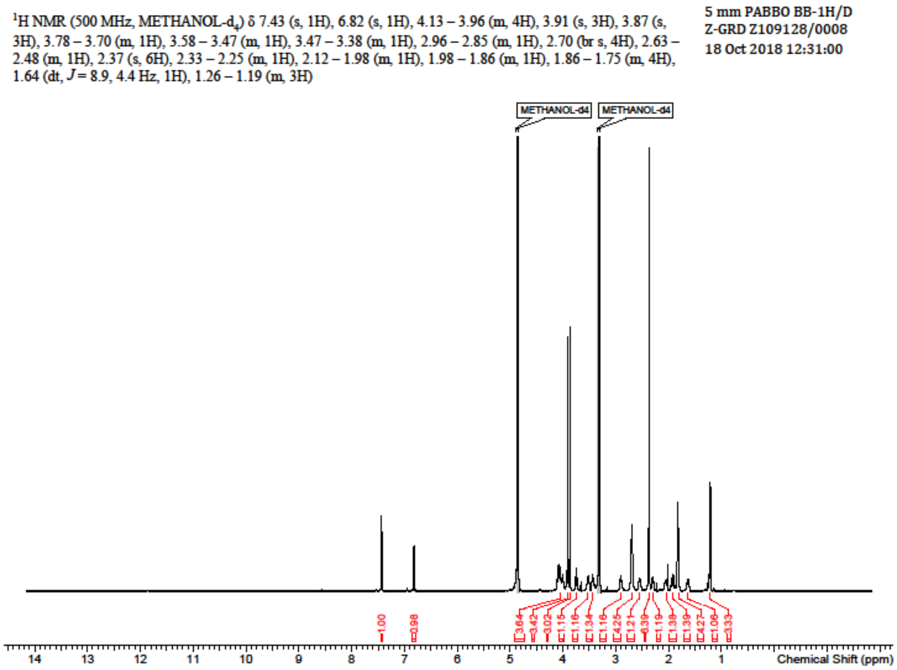

¹H NMR (500 MHz, METHANOL-d4) δ 7.43 (s, 1H), 6.82 (s, 1H), 4.13 – 3.96 (m, 4H), 3.91 (s, 3H), 3.87 (s, 3H), 3.78 – 3.70 (m, 1H), 3.58 – 3.47 (m, 1H), 3.47 – 3.38 (m, 1H), 2.96 – 2.85 (m, 1H), 2.70 (br s, 4H), 2.63 – 2.48 (m, 1H), 2.37 (s, 6H), 2.33 – 2.25 (m, 1H), 2.12 – 1.98 (m, 1H), 1.98 – 1.86 (m, 1H), 1.86 – 1.75 (m, 4H), 1.64 (dt, J = 8.9, 4.4 Hz, 1H), 1.26 – 1.19 (m, 3H)
5 mm PABBO BB-1H/D
Z-GRD Z109128/0008
18 Oct 2018 12:31:00
METHANOL-d4
METHANOL-d4
Chemical Shift (ppm)


18721

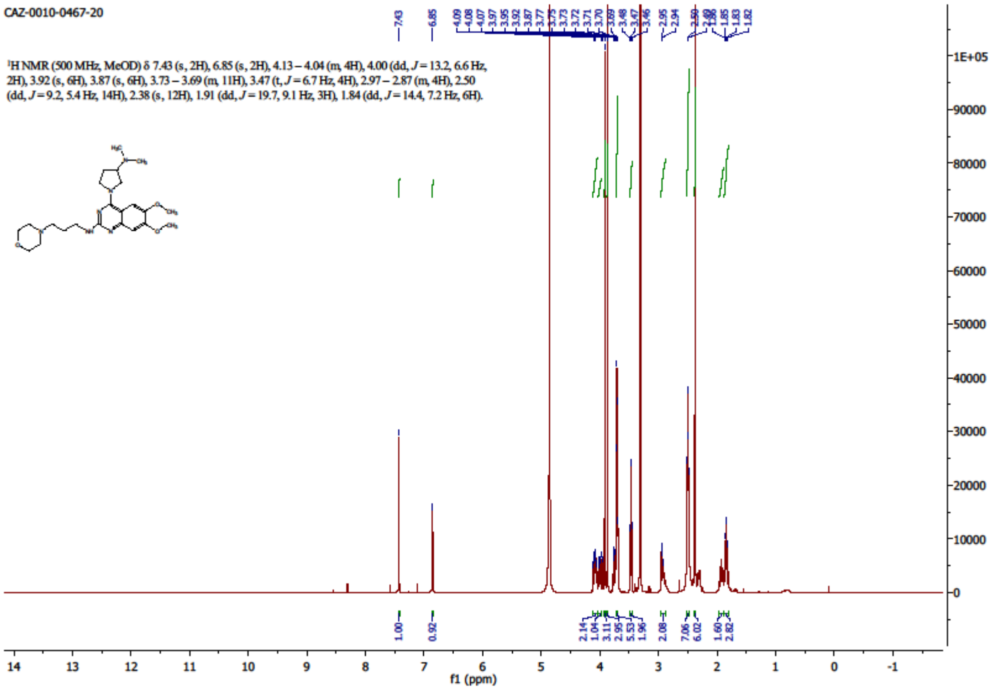

CAZ-0010-0467-20
¹H NMR (500 MHz, MeOD) δ 7.43 (s, 2H), 6.85 (s, 2H), 4.13 – 4.04 (m, 4H), 4.00 (dd, J = 13.2, 6.6 Hz, 2H), 3.92 (s, 6H), 3.87 (s, 6H), 3.73 – 3.69 (m, 11H), 3.47 (t, J = 6.7 Hz, 4H), 2.97 – 2.87 (m, 4H), 2.50 (dd, J = 9.2, 5.4 Hz, 14H), 2.38 (s, 12H), 1.91 (dd, J = 19.7, 9.1 Hz, 3H), 1.84 (dd, J = 14.4, 7.2 Hz, 6H).
f1 (ppm)

18632

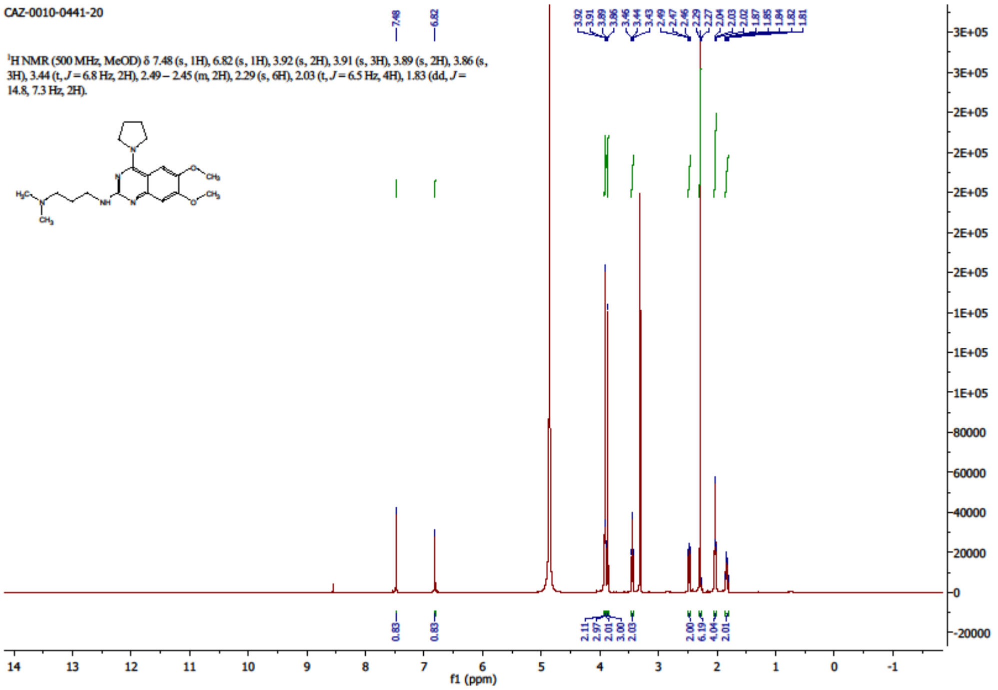


19016

$^1$H NMR (500 MHz, METHANOL-$d_4$) δ 7.06 (s, 1H), 6.86 (s, 1H), 3.92 (s, 3H), 3.87 (s, 3H), 3.57 (br s, 4H), 3.51 – 3.34 (m, 2H), 2.51 – 2.41 (m, 2H), 2.28 (s, 6H), 1.90 – 1.80 (m, 3H), 1.80 – 1.73 (m, 6H)

5 mm PABBO BB-1H/D
Z-GRD Z109128/0008
11 Oct 2018 10:21:52

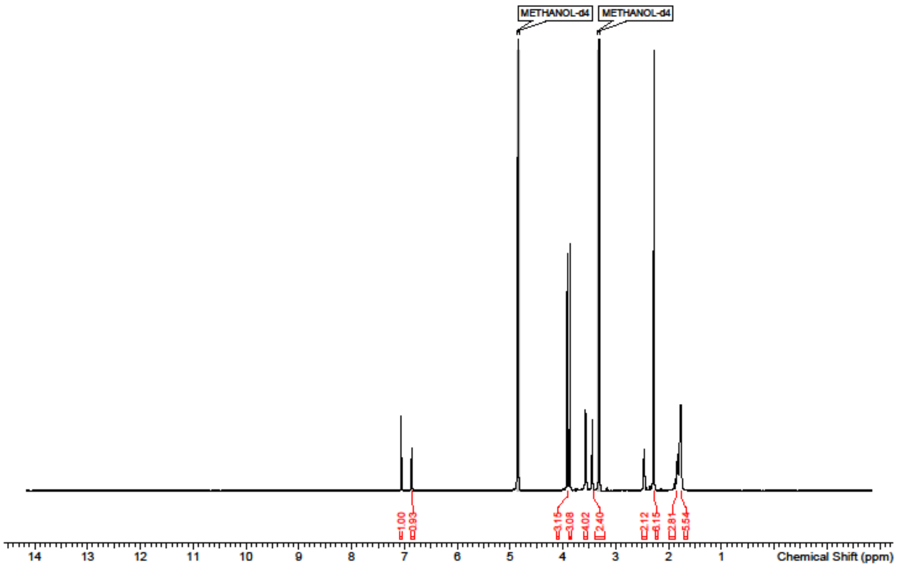

18716

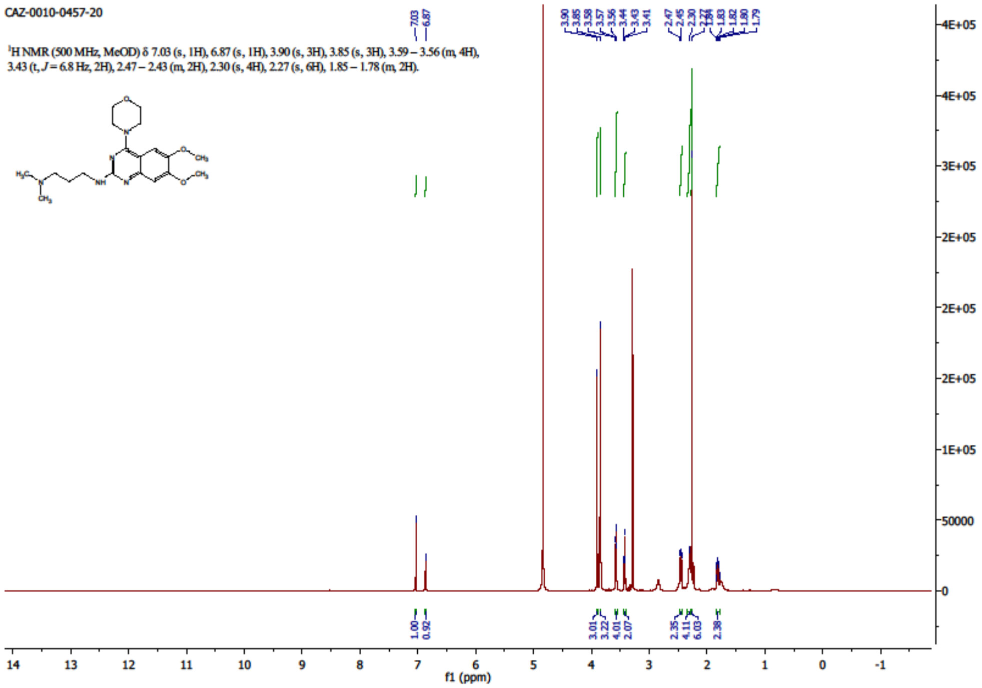
CAZ-0010-0457-20
¹H NMR (500 MHz, MeOD) δ 7.03 (s, 1H), 6.87 (s, 1H), 3.90 (s, 3H), 3.85 (s, 3H), 3.59 – 3.56 (m, 4H), 3.43 (t, J = 6.8 Hz, 2H), 2.47 – 2.43 (m, 2H), 2.30 (s, 4H), 2.27 (s, 6H), 1.85 – 1.78 (m, 2H).
f1 (ppm)

19093

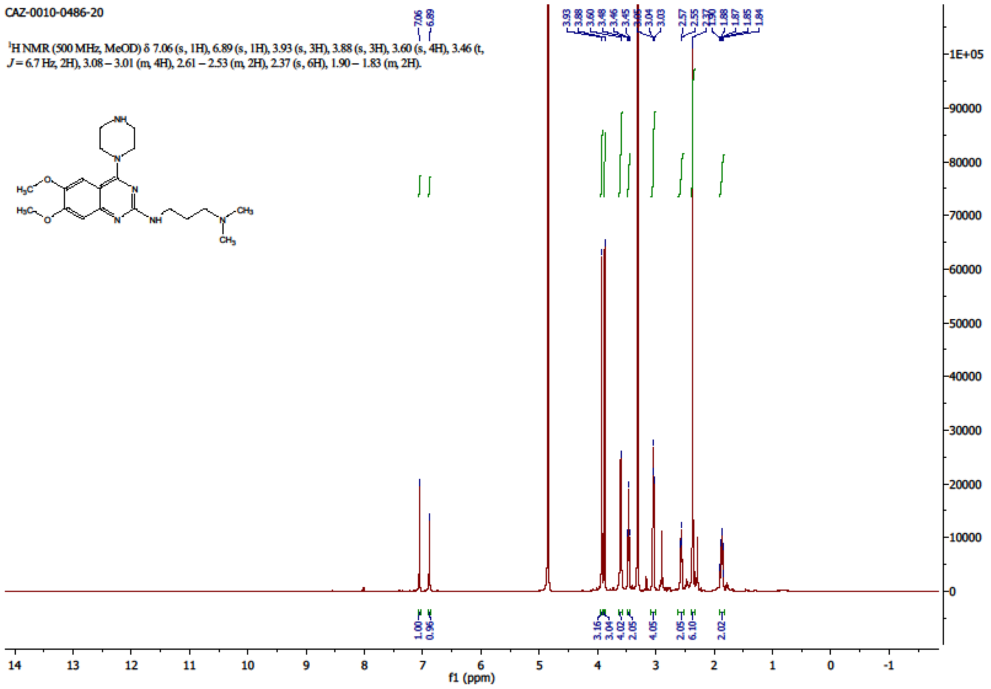
CAZ-0010-0486-20
¹H NMR (500 MHz, MeOD) δ 7.06 (s, 1H), 6.89 (s, 1H), 3.93 (s, 3H), 3.88 (s, 3H), 3.60 (s, 4H), 3.46 (t, J = 6.7 Hz, 2H), 3.08 – 3.01 (m, 4H), 2.61 – 2.53 (m, 2H), 2.37 (s, 6H), 1.90 – 1.83 (m, 2H).
f1 (ppm)

18631

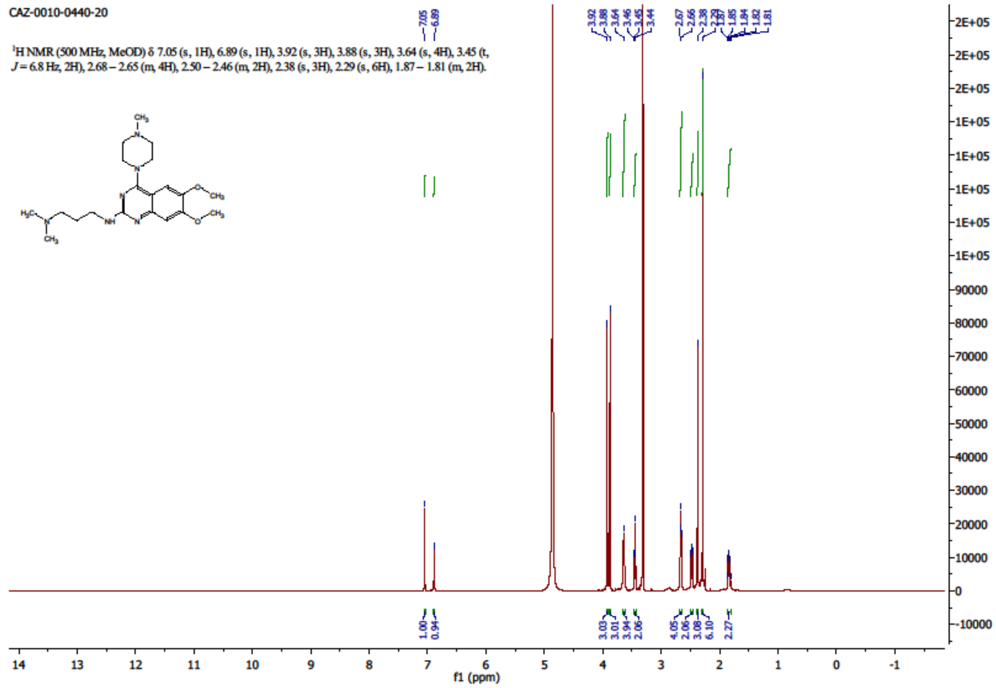
CAZ-0010-0440-20
1H NMR (500 MHz, MeOD) δ 7.05 (s, 1H), 6.89 (s, 1H), 3.92 (s, 3H), 3.88 (s, 3H), 3.64 (s, 4H), 3.45 (t, J = 6.8 Hz, 2H), 2.68 – 2.65 (m, 4H), 2.50 – 2.46 (m, 2H), 2.38 (s, 3H), 2.29 (s, 6H), 1.87 – 1.81 (m, 2H).
f1 (ppm)

18718

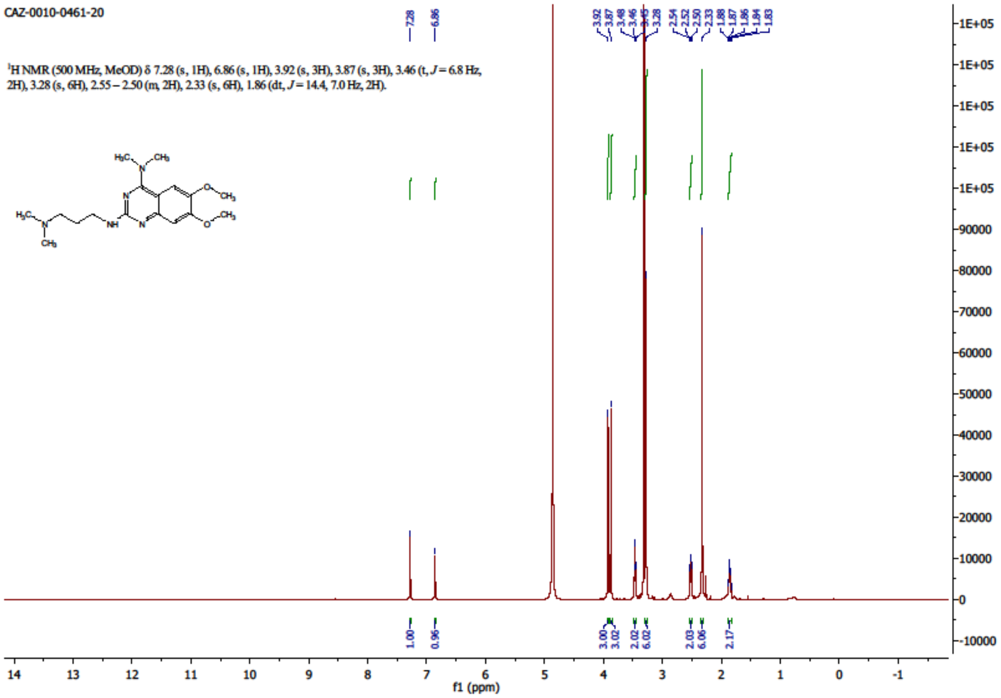
CAZ-0010-0461-20
1H NMR (500 MHz, MeOD) δ 7.28 (s, 1H), 6.86 (s, 1H), 3.92 (s, 3H), 3.87 (s, 3H), 3.46 (t, J = 6.8 Hz, 2H), 3.28 (s, 6H), 2.55 – 2.50 (m, 2H), 2.33 (s, 6H), 1.86 (dt, J = 14.4, 7.0 Hz, 2H).
f1 (ppm)

18745

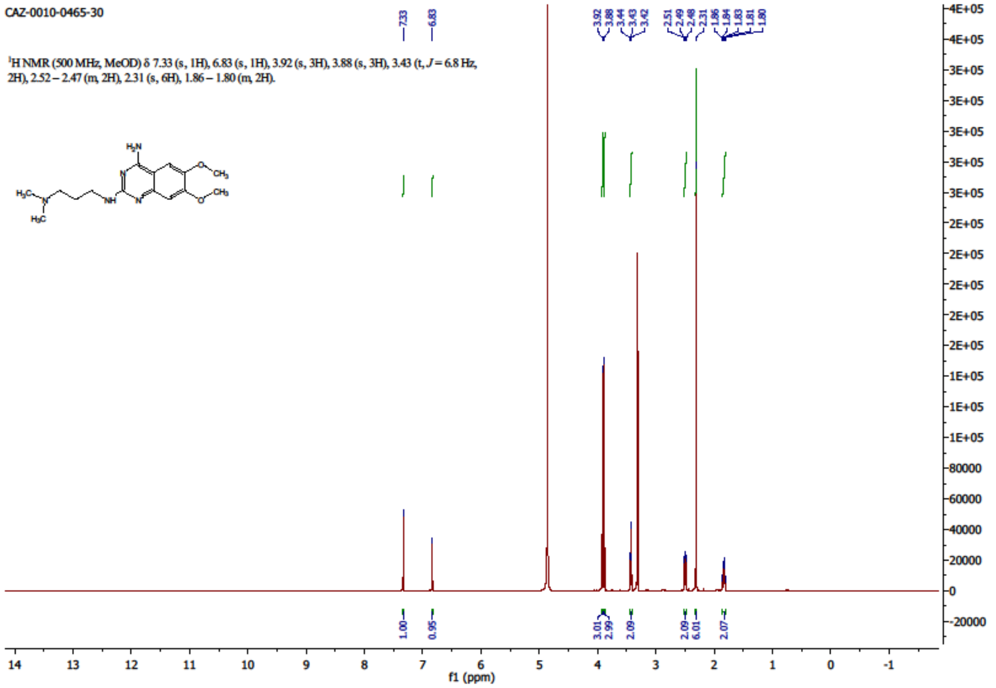
CAZ-0010-0465-30
¹H NMR (500 MHz, MeOD) δ 7.33 (s, 1H), 6.83 (s, 1H), 3.92 (s, 3H), 3.88 (s, 3H), 3.43 (t, J = 6.8 Hz, 2H), 2.52 – 2.47 (m, 2H), 2.31 (s, 6H), 1.86 – 1.80 (m, 2H).
f1 (ppm)

19102

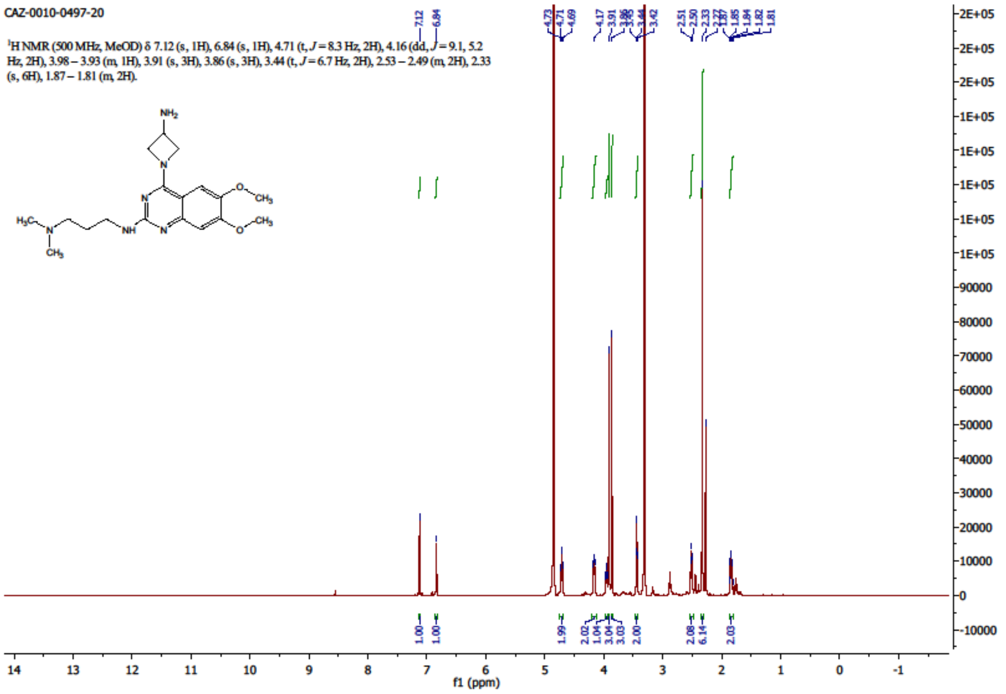
CAZ-0010-0497-20
¹H NMR (500 MHz, MeOD) δ 7.12 (s, 1H), 6.84 (s, 1H), 4.71 (t, J = 8.3 Hz, 2H), 4.16 (dd, J = 9.1, 5.2 Hz, 2H), 3.98 – 3.93 (m, 1H), 3.91 (s, 3H), 3.86 (s, 3H), 3.44 (t, J = 6.7 Hz, 2H), 2.53 – 2.49 (m, 2H), 2.33 (s, 6H), 1.87 – 1.81 (m, 2H).
f1 (ppm)

18629

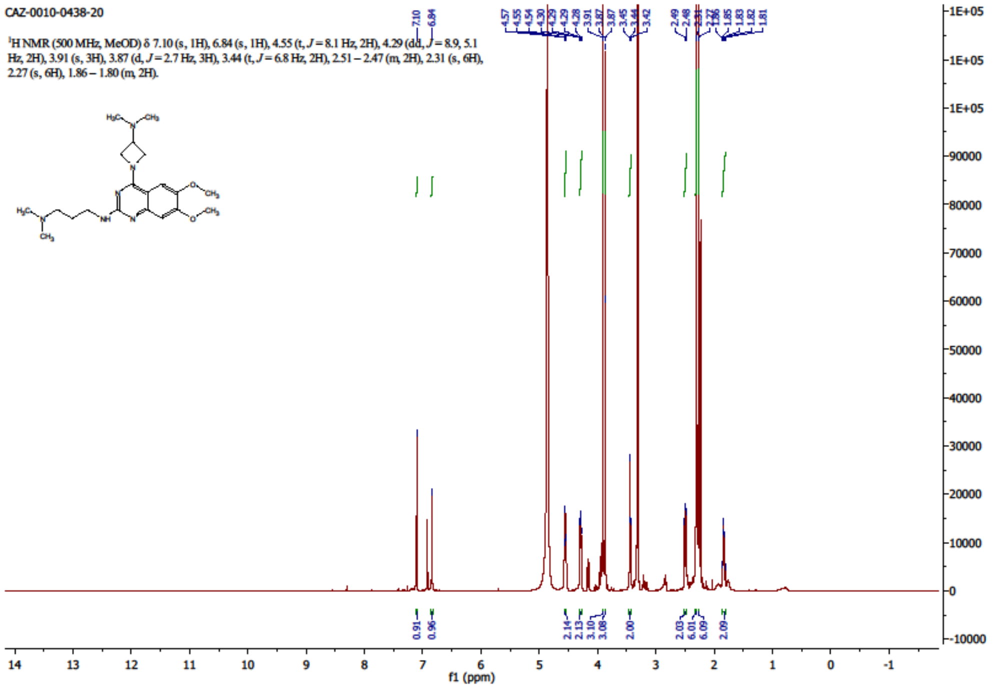
CAZ-0010-0438-20
¹H NMR (500 MHz, MeOD) δ 7.10 (s, 1H), 6.84 (s, 1H), 4.55 (t, J = 8.1 Hz, 2H), 4.29 (dd, J = 8.9, 5.1 Hz, 2H), 3.91 (s, 3H), 3.87 (d, J = 2.7 Hz, 3H), 3.44 (t, J = 6.8 Hz, 2H), 2.51 – 2.47 (m, 2H), 2.31 (s, 6H), 2.27 (s, 6H), 1.86 – 1.80 (m, 2H).
f1 (ppm)

19099

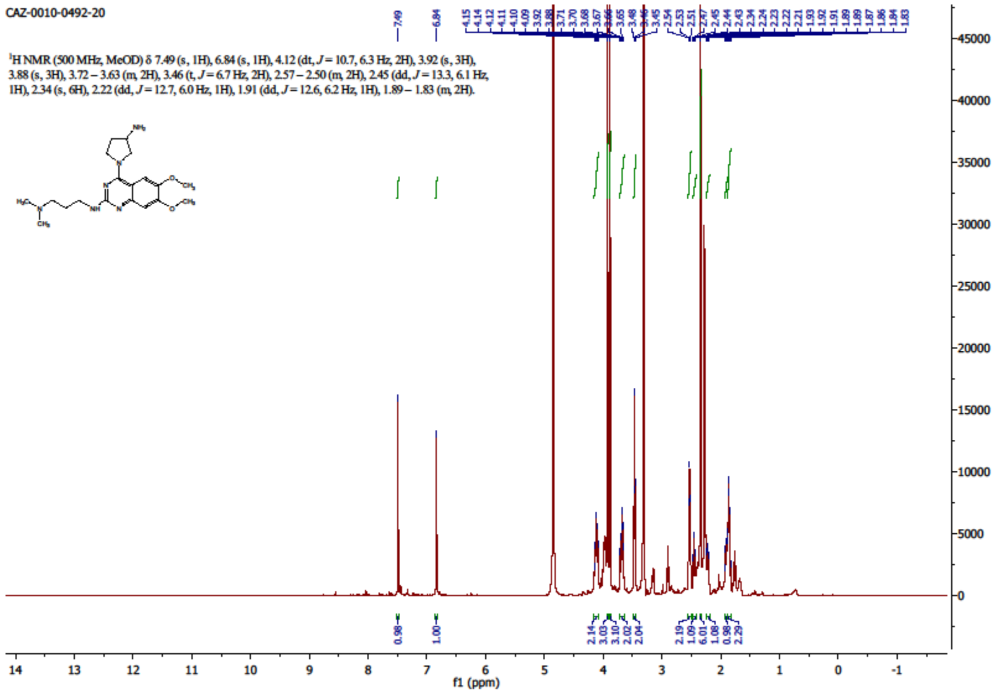
CAZ-0010-0492-20
¹H NMR (500 MHz, MeOD) δ 7.49 (s, 1H), 6.84 (s, 1H), 4.12 (dt, J = 10.7, 6.3 Hz, 2H), 3.92 (s, 3H), 3.88 (s, 3H), 3.72 – 3.63 (m, 2H), 3.46 (t, J = 6.7 Hz, 2H), 2.57 – 2.50 (m, 2H), 2.45 (dd, J = 13.3, 6.1 Hz, 1H), 2.34 (s, 6H), 2.22 (dd, J = 12.7, 6.0 Hz, 1H), 1.91 (dd, J = 12.6, 6.2 Hz, 1H), 1.89 – 1.83 (m, 2H).
f1 (ppm)

18627

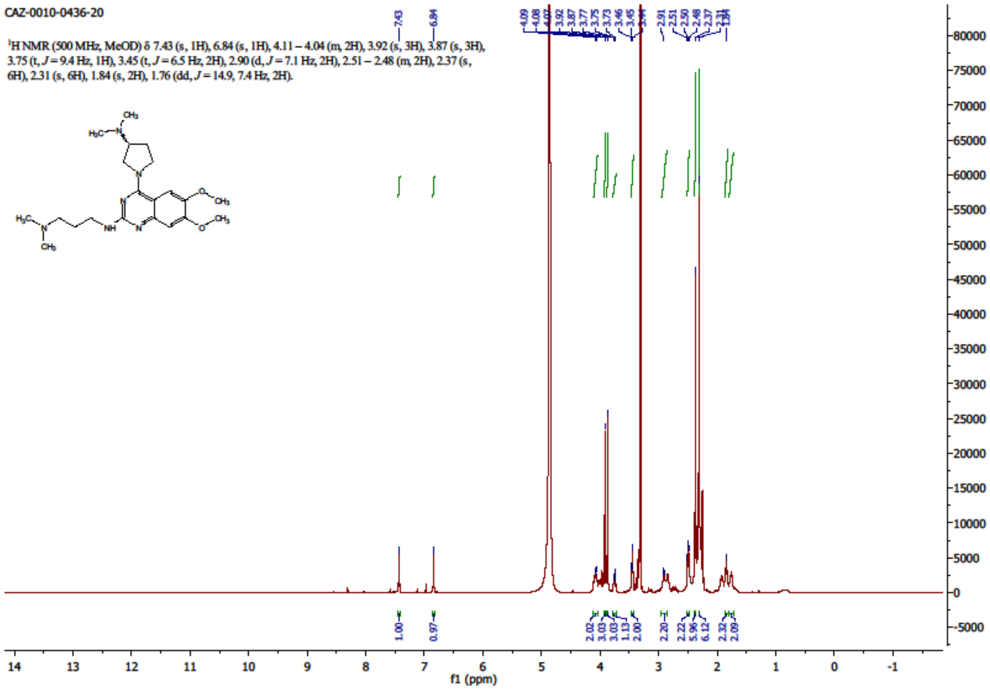
CAZ-0010-0436-20
1H NMR (500 MHz, MeOD) δ 7.43 (s, 1H), 6.84 (s, 1H), 4.11 – 4.04 (m, 2H), 3.92 (s, 3H), 3.87 (s, 3H), 3.75 (t, J = 9.4 Hz, 1H), 3.45 (t, J = 6.5 Hz, 2H), 2.90 (d, J = 7.1 Hz, 2H), 2.51 – 2.48 (m, 2H), 2.37 (s, 6H), 2.31 (s, 6H), 1.84 (s, 2H), 1.76 (dd, J = 14.9, 7.4 Hz, 2H).
f1 (ppm)

18628

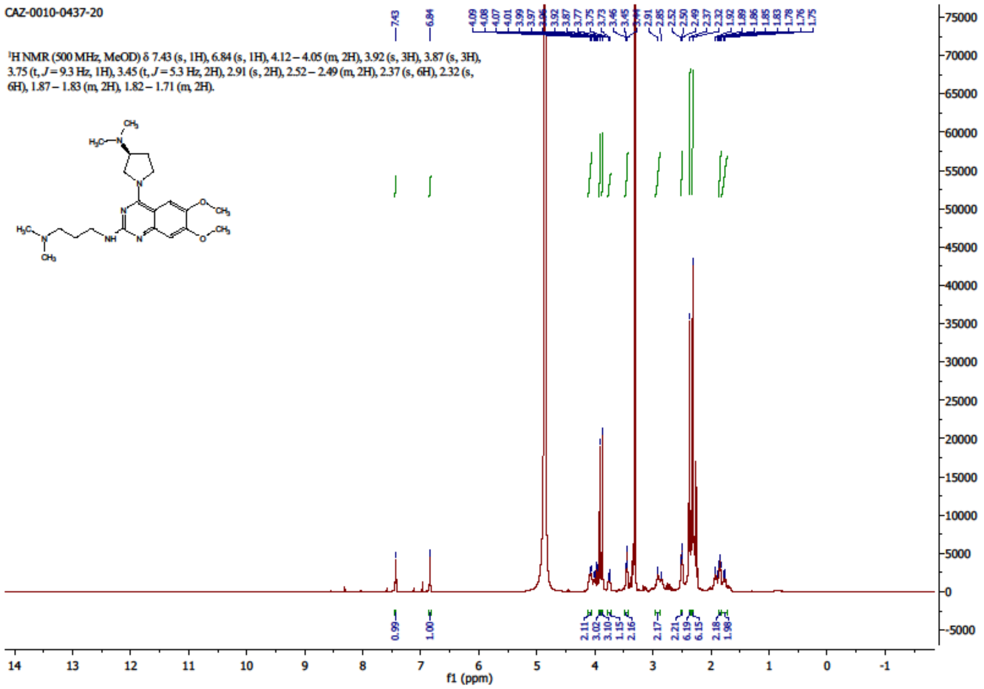
CAZ-0010-0437-20
1H NMR (500 MHz, MeOD) δ 7.43 (s, 1H), 6.84 (s, 1H), 4.12 – 4.05 (m, 2H), 3.92 (s, 3H), 3.87 (s, 3H), 3.75 (t, J = 9.3 Hz, 1H), 3.45 (t, J = 5.3 Hz, 2H), 2.91 (s, 2H), 2.52 – 2.49 (m, 2H), 2.37 (s, 6H), 2.32 (s, 6H), 1.87 – 1.83 (m, 2H), 1.82 – 1.71 (m, 2H).
f1 (ppm)

18630

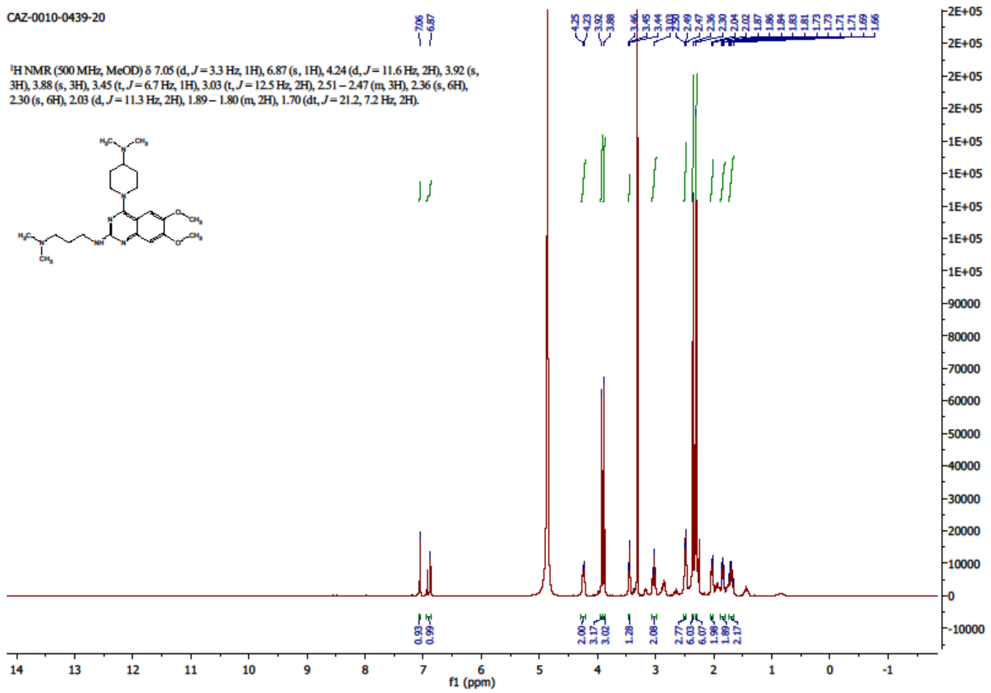
CAZ-0010-0439-20
¹H NMR (500 MHz, MeOD) δ 7.05 (d, J = 3.3 Hz, 1H), 6.87 (s, 1H), 4.24 (d, J = 11.6 Hz, 2H), 3.92 (s, 3H), 3.88 (s, 3H), 3.45 (t, J = 6.7 Hz, 1H), 3.03 (t, J = 12.5 Hz, 2H), 2.51 – 2.47 (m, 3H), 2.36 (s, 6H), 2.30 (s, 6H), 2.03 (d, J = 11.3 Hz, 2H), 1.89 – 1.80 (m, 2H), 1.70 (dt, J = 21.2, 7.2 Hz, 2H).
f1 (ppm)

19101

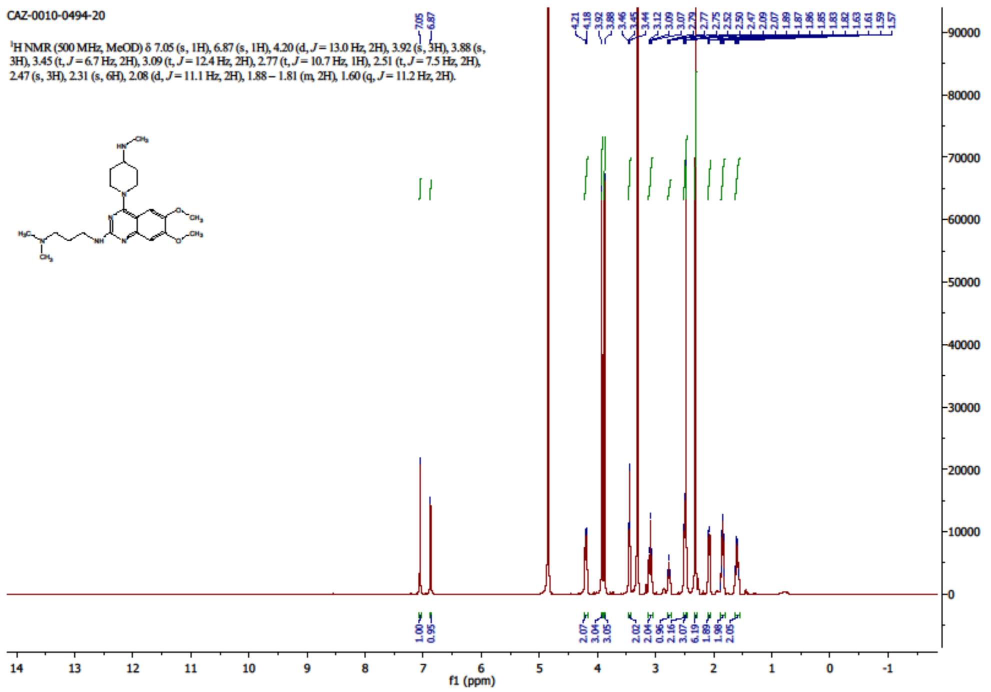
CAZ-0010-0494-20
¹H NMR (500 MHz, MeOD) δ 7.05 (s, 1H), 6.87 (s, 1H), 4.20 (d, J = 13.0 Hz, 2H), 3.92 (s, 3H), 3.88 (s, 3H), 3.45 (t, J = 6.7 Hz, 2H), 3.09 (t, J = 12.4 Hz, 2H), 2.77 (t, J = 10.7 Hz, 1H), 2.51 (t, J = 7.5 Hz, 2H), 2.47 (s, 3H), 2.31 (s, 6H), 2.08 (d, J = 11.1 Hz, 2H), 1.88 – 1.81 (m, 2H), 1.60 (q, J = 11.2 Hz, 2H).
f1 (ppm)

19100

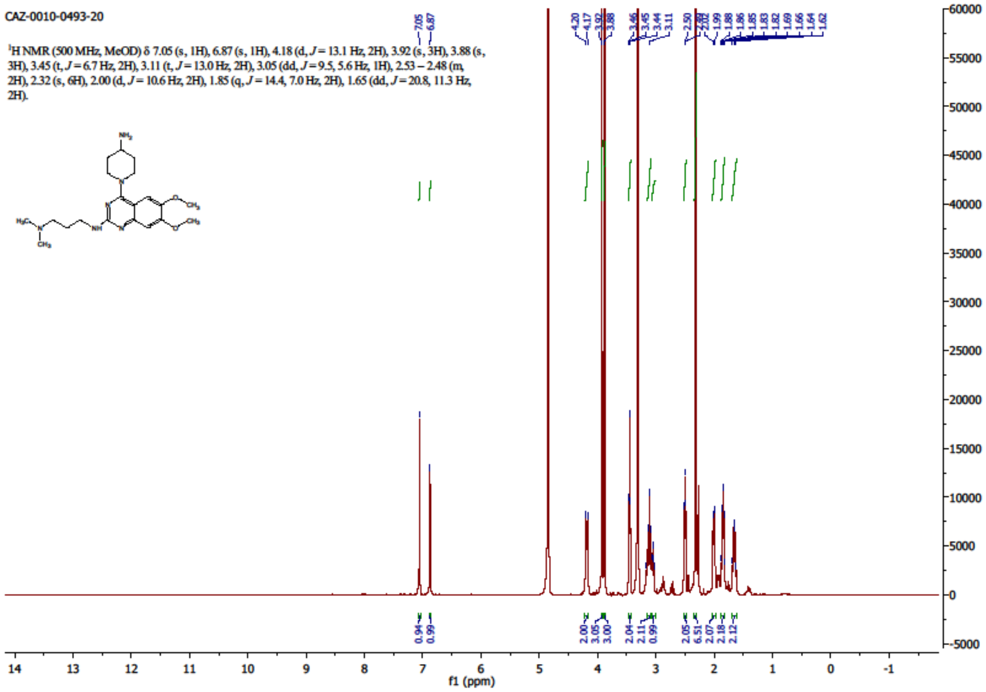
CAZ-0010-0493-20
1H NMR (500 MHz, MeOD) δ 7.05 (s, 1H), 6.87 (s, 1H), 4.18 (d, J = 13.1 Hz, 2H), 3.92 (s, 3H), 3.88 (s, 3H), 3.45 (t, J = 6.7 Hz, 2H), 3.11 (t, J = 13.0 Hz, 2H), 3.05 (dd, J = 9.5, 5.6 Hz, 1H), 2.53 – 2.48 (m, 2H), 2.32 (s, 6H), 2.00 (d, J = 10.6 Hz, 2H), 1.85 (q, J = 14.4, 7.0 Hz, 2H), 1.65 (dd, J = 20.8, 11.3 Hz, 2H).
f1 (ppm)

19098

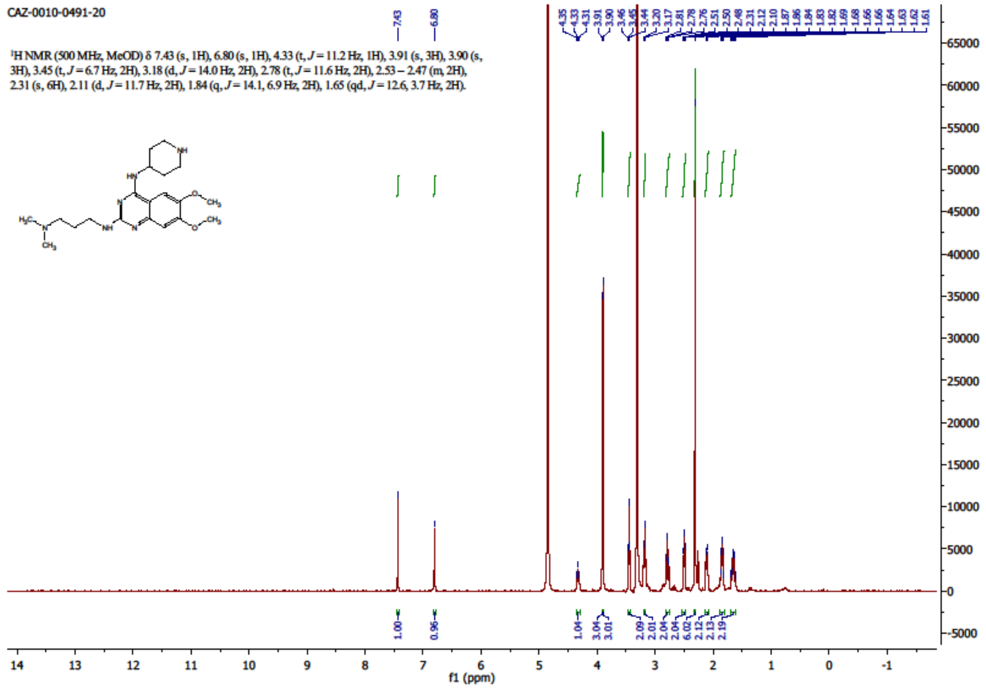
CAZ-0010-0491-20
1H NMR (500 MHz, MeOD) δ 7.43 (s, 1H), 6.80 (s, 1H), 4.33 (t, J = 11.2 Hz, 1H), 3.91 (s, 3H), 3.90 (s, 3H), 3.45 (t, J = 6.7 Hz, 2H), 3.18 (d, J = 14.0 Hz, 2H), 2.78 (t, J = 11.6 Hz, 2H), 2.53 – 2.47 (m, 2H), 2.31 (s, 6H), 2.11 (d, J = 11.7 Hz, 2H), 1.84 (q, J = 14.1, 6.9 Hz, 2H), 1.65 (qd, J = 12.6, 3.7 Hz, 2H).
f1 (ppm)

### HPLC chromatograms

17251

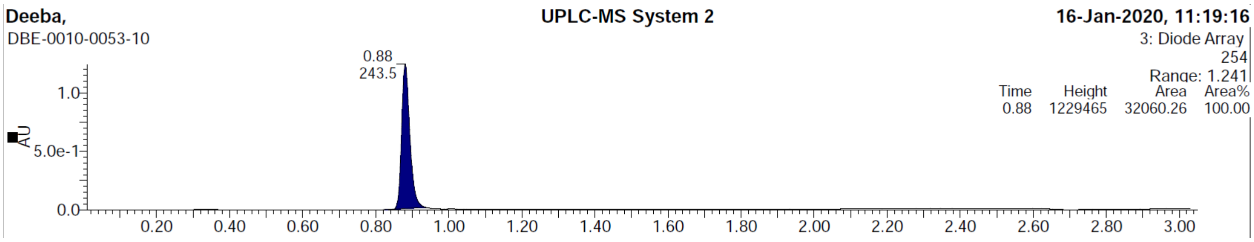


18633

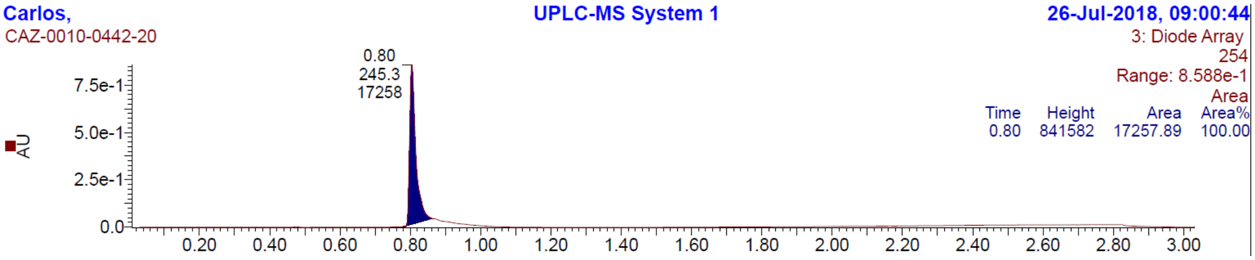


18634

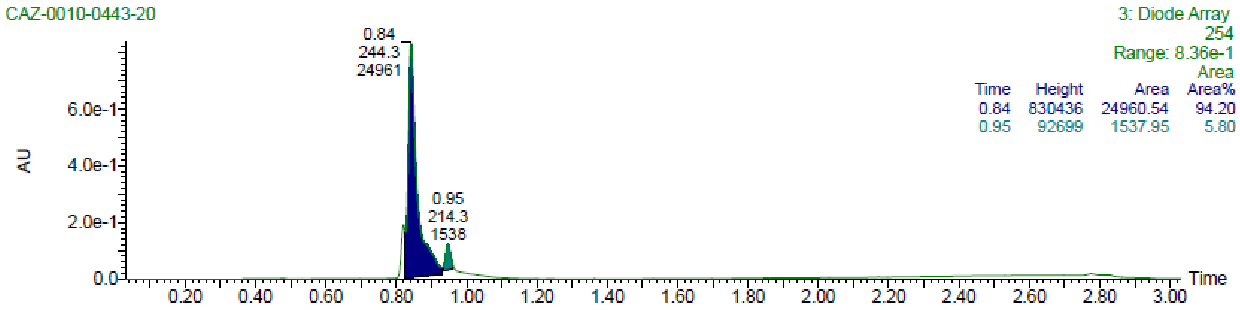


18635

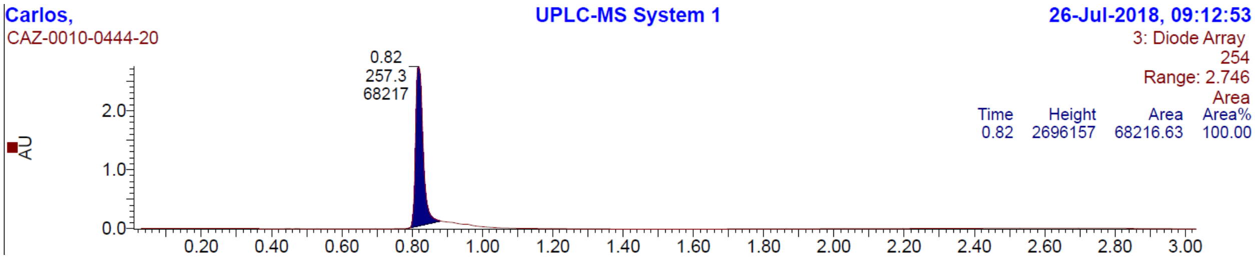


18636

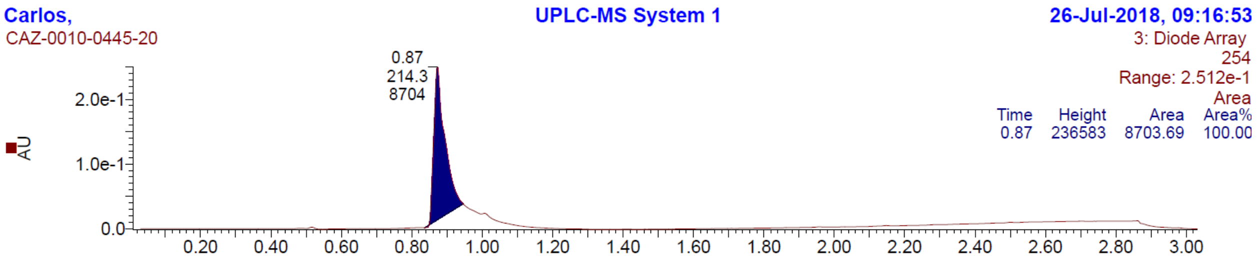


18637

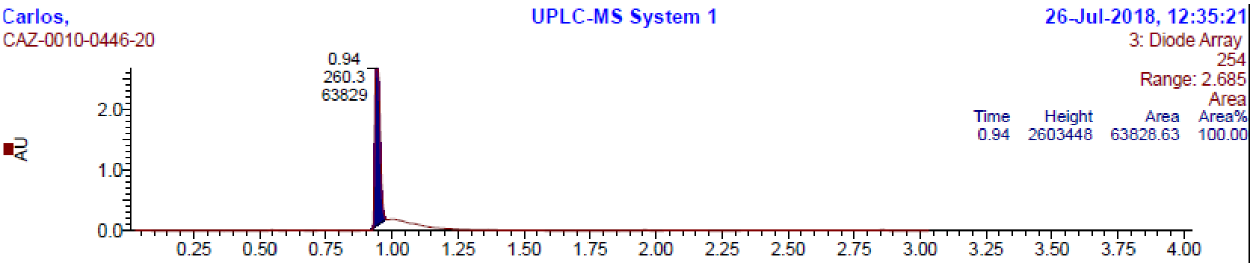


19018

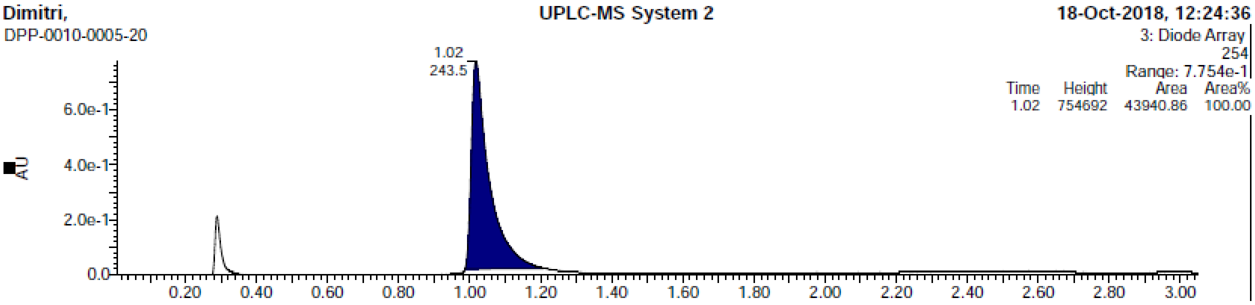


18721

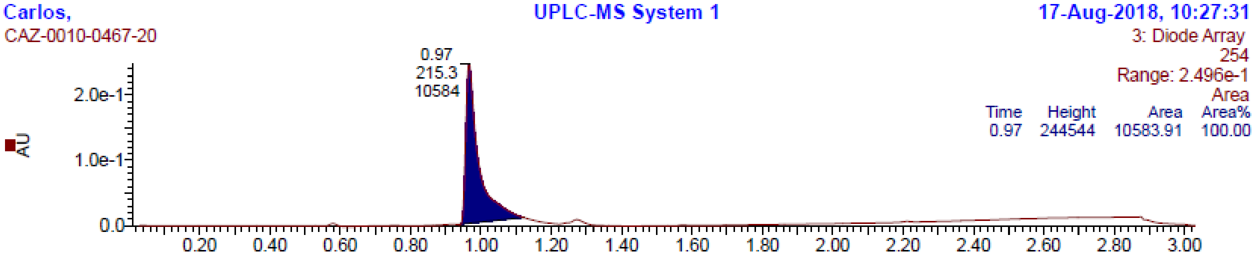


18632

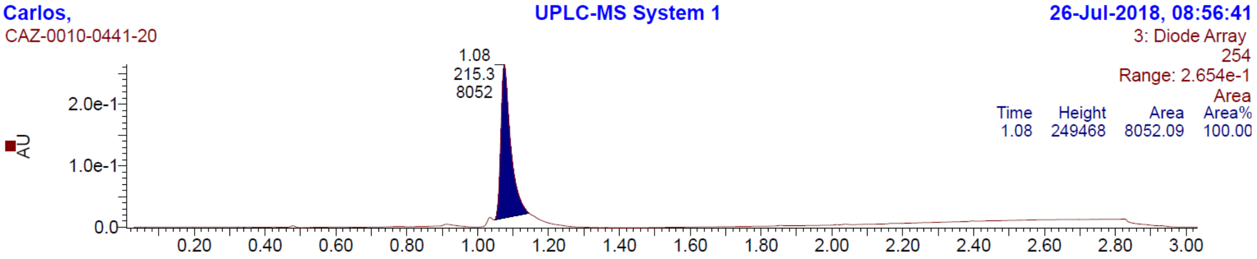


19016

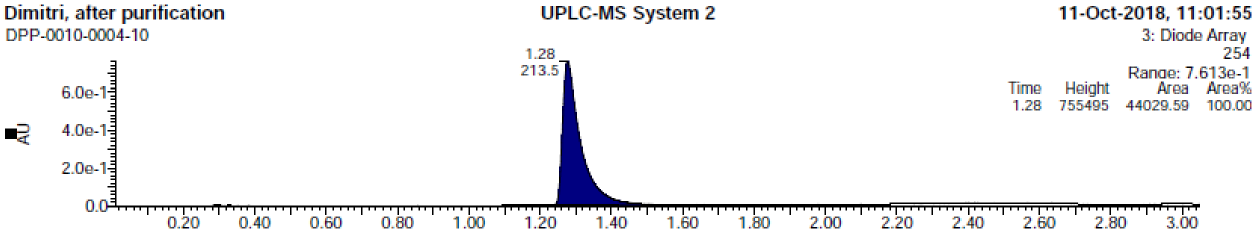


18716

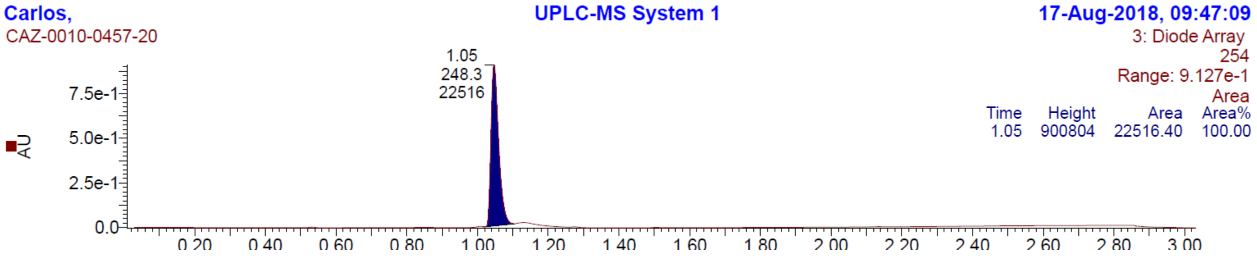


19093

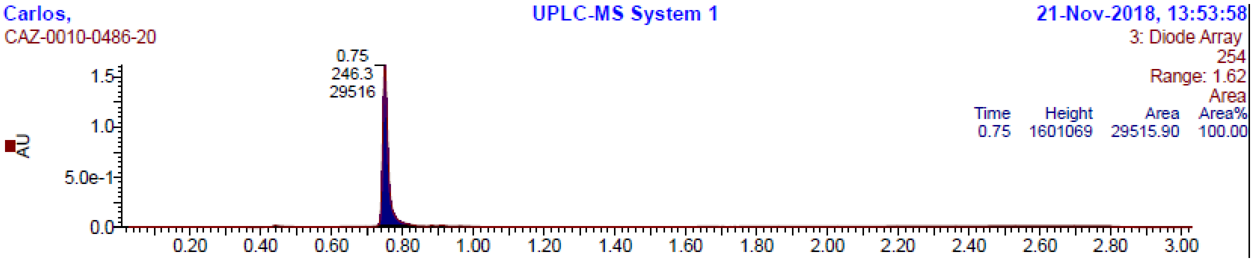


18631

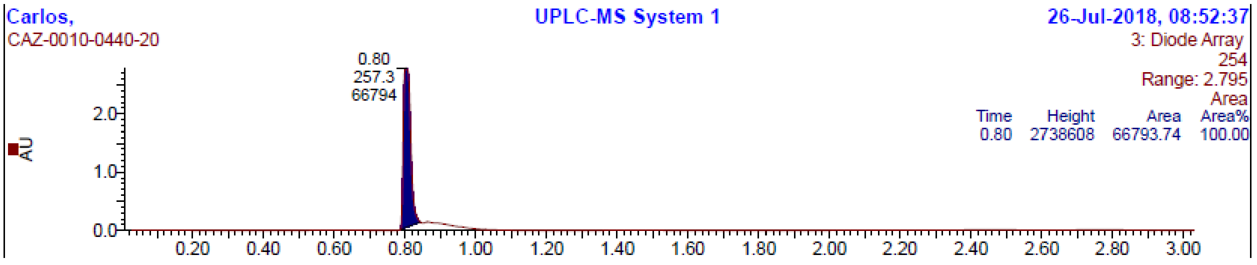


18718

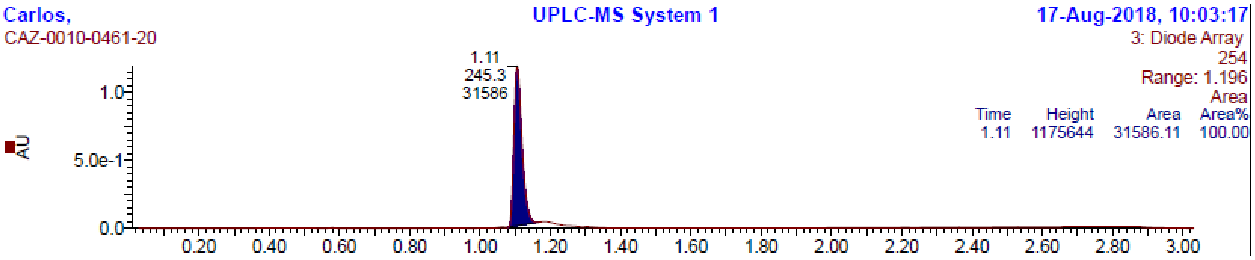


18745

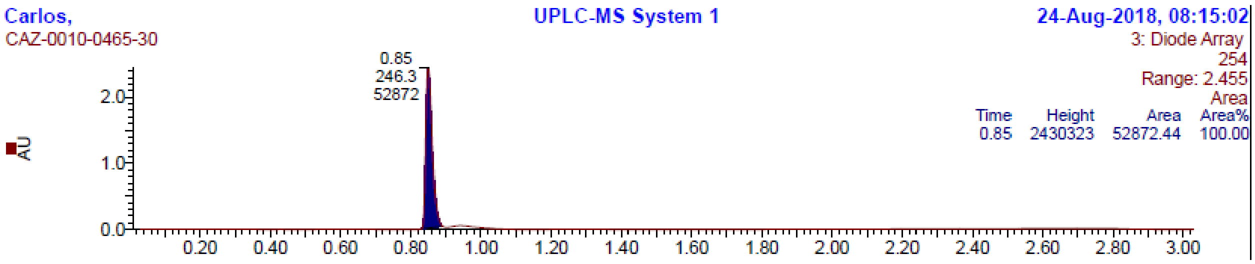


19102

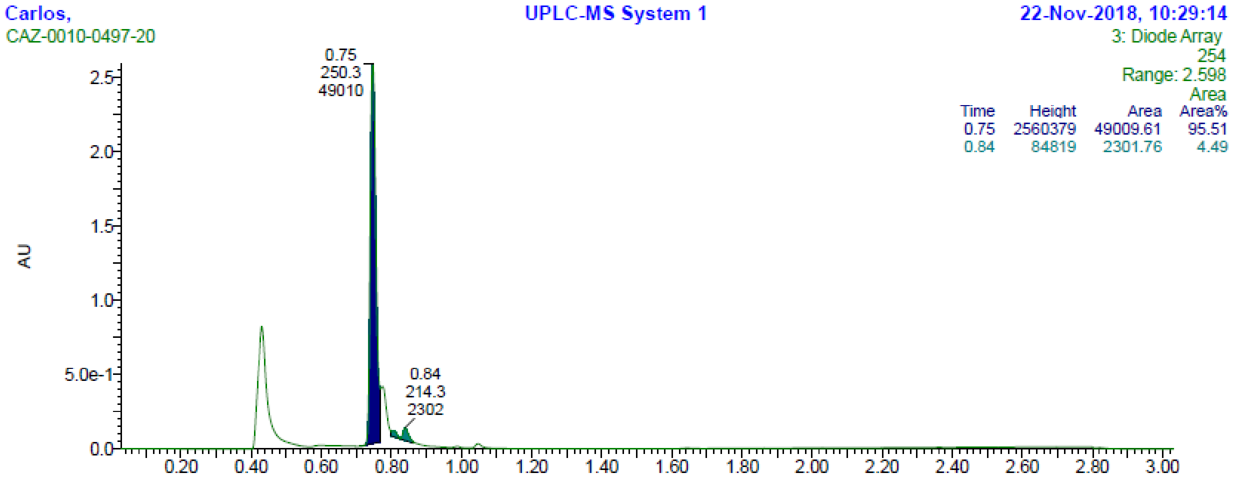


18629

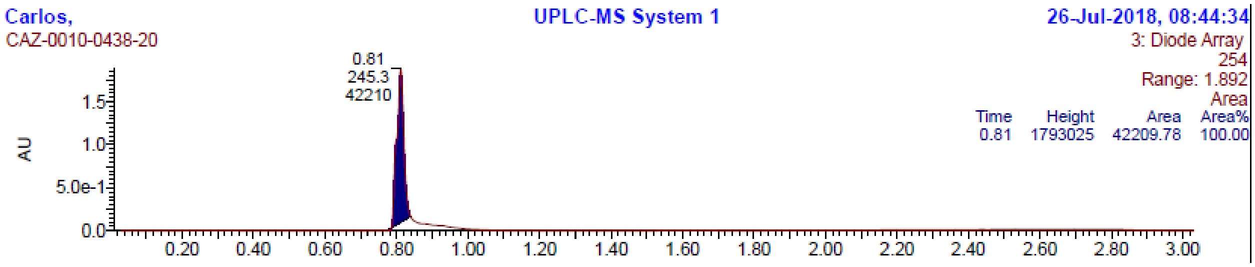


19099

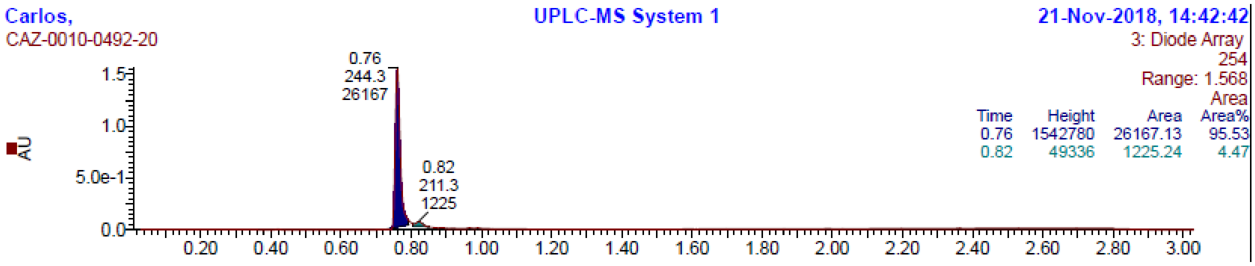


18627

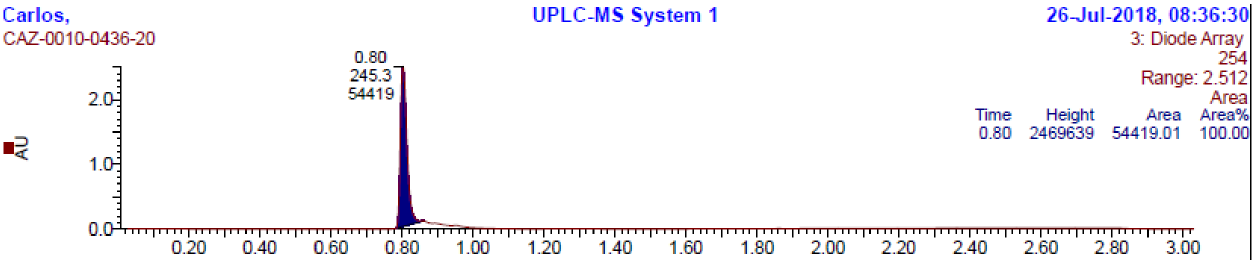


18628

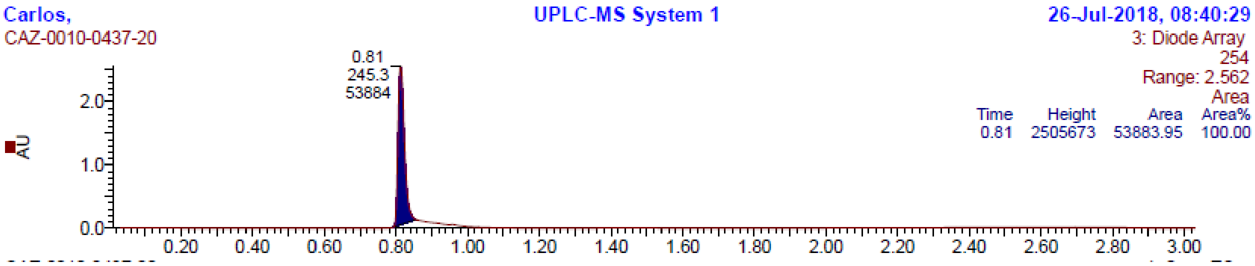


18630

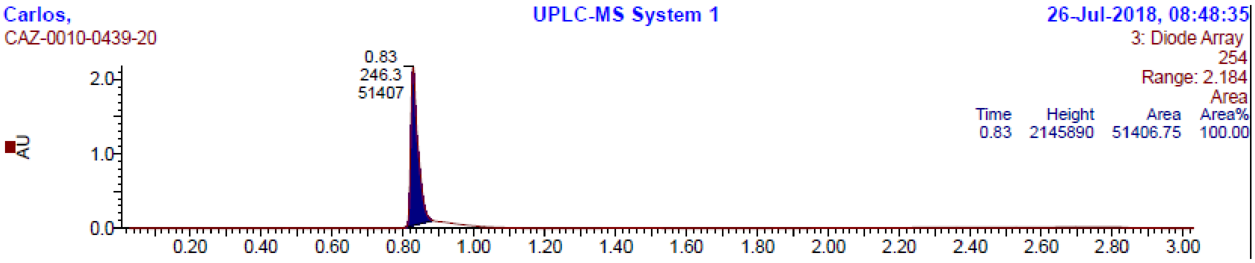


19101

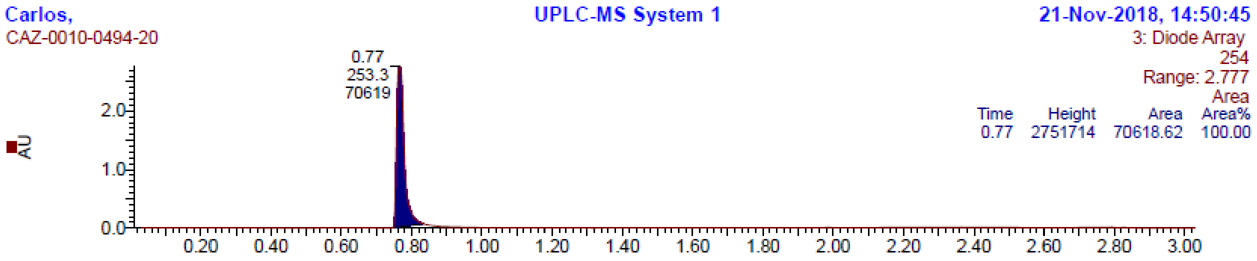
Carlos,
CAZ-0010-0494-20
UPLC-MS System 1
21-Nov-2018, 14:50:45
3: Diode Array
254
Range: 2.777
Area
Time Height Area Area%
0.77 2751714 70618.62 100.00
0.77
253.3
70619
AU

19100

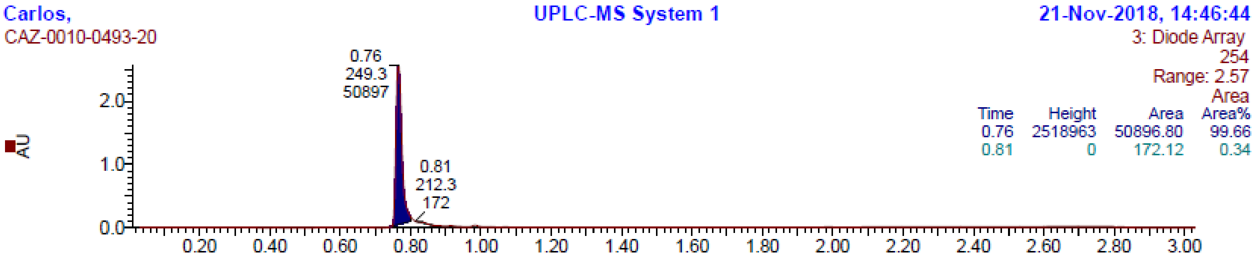
Carlos,
CAZ-0010-0493-20
UPLC-MS System 1
21-Nov-2018, 14:46:44
3: Diode Array
254
Range: 2.57
Area
Time Height Area Area%
0.76 2518963 50896.80 99.66
0.81 0 172.12 0.34
0.76
249.3
50897
0.81
212.3
172
AU

19098

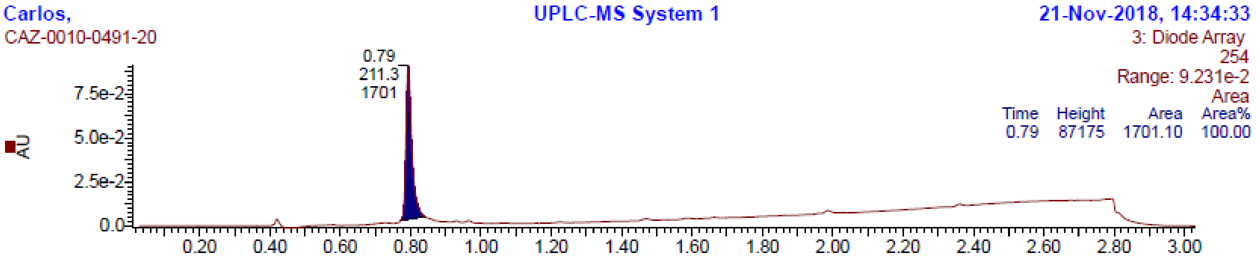
Carlos,
CAZ-0010-0491-20
UPLC-MS System 1
21-Nov-2018, 14:34:33
3: Diode Array
254
Range: 9.231e-2
Area
Time Height Area Area%
0.79 87175 1701.10 100.00
0.79
211.3
1701
AU
